# Spatial isotope tracing maps compartmentalized nutrient use across the adult fruit fly

**DOI:** 10.64898/2026.10.06.757091

**Authors:** Sun Jin Moon, Kerui Huang, Jenna L. E. Blum, Connie New, Yooha Park, Yuchen Chen, Hiroshi Nishida, Veronika Pister, Md Ausrafuggaman Nahid, Yanhui Hu, Jason Tennessen, Ernest Fraenkel, John M. Asara, Shawn M. Davidson, Norbert Perrimon

## Abstract

*Drosophila melanogaster* is a powerful model for studying systemic metabolism. Although metabolism is compartmentalized across tissues, prior work in adult *Drosophila* has mostly measured it in dissected tissues rather than *in situ*, limiting the spatial information that can be gained. Here, we combine whole-fly massspectrometry imaging with stable-isotope tracing and flux modeling to quantify metabolic activities across the adult fruit fly in a spatially resolved manner. Across 68 tracer-fed Oregon-R female flies and 187,709 pixels, we map 81 metabolites onto eight annotated regions, including muscle, fat body, gut, and nervous system. Glucose labeling and fitted relative fluxes indicate higher glycolytic and tricarboxylic acid cycle turnover in thoracic muscle than in abdominal fat body; proline labels the nervous system and eye; glutamine the abdominal reproductive system and gut. Alanine carries glucose-derived carbon through the hemolymph and transfers nitrogen into glutamate. In flies bearing yorkie-driven gut tumors, alanine contribution to malate rises in gut and abdomen, and reducing gut alanine uptake lowers tumor burden. Together, this work establishes a framework for quantitative spatial metabolic analysis, showing compartmentalized nutrient use across the adult fruit fly.

## Main

Metabolism is highly compartmentalized at multiple biological scales (*1*). At the organismal level, maintenance of systemic metabolic homeostasis requires coordination among multiple tissues that respond to nutrient and environmental changes (*2*, *3*). Disruption of this coordination contributes to conditions such as cancer, aging, and neurodegeneration. Although conventional metabolomics and isotope tracing techniques have revealed metabolic states and activities in isolated tissues and whole animals, sample homogenization prevents access to spatial information (*4–6*). This limits the direct assessment of local tissue metabolism and its interaction with the environment. Thus, systemic metabolic studies need approaches that preserve spatial metabolic information.

Recent advances in mass-spectrometry imaging (MSI) have enabled spatial metabolomics, allowing metabolites (*7*), lipids (*8*, *9*), glycogen (*10*), and other small molecules to be mapped directly in tissue orwhole animal sections (*11–13*). Combining stable isotope tracing with MSI has allowed the direct assessment of spatially resolved metabolic pathway activities in kidney, intestine, brain, and tumors *in situ* (*13–16*).

*Drosophila melanogaster*, a powerful model for studying systemic metabolism, offers a particularly favorable system for this approach. Its relatively small adult body size (∼ 3 × 2 mm in length and height) places the major metabolic tissues, such as muscle, fat body, gut, and others, within a single section, making simultaneous, spatially resolved measurement of the whole animal feasible (*17–19*). MSI has been applied to whole-body fly sections to map lipid classes (*20*, *21*) and to dissected brains (*22–24*). Cryo-OrbiSIMS has been used to map hydrocarbons and wax esters on adult fly cuticle (*25*). Multi-isotope imaging mass spectrometry has also quantified lipid turnover in individual lipid droplets in fly (*26*). However, these studies have primarily focused on lipid distributions and turnover, and have not resolved tissue-level metabolic activities in central carbon metabolism. Computational approaches, such as tissue-level metabolic network reconstruction and flux analysis, have enabled quantitative investigation of tissue-level metabolism, but these methods rely on indirect gene expression measurements (*27*). Bulk metabolomics and isotope tracing have also been applied to dissected anatomical regions that serve as proxies for tissue-enriched compartments, such as the thorax for muscle and the abdomen for fat body (*28–30*). However, these approaches do not delineate tissue-level metabolite profiles or activities from bulk samples.

To address these limitations, we combine whole-fly MSI with stable-isotope tracing and flux analyses to quantitate spatially resolved metabolic activities across the adult fruit fly. We first establish a spatial metabolic atlas that resolves eight annotated regions, including muscle, fat body, gut, and the central nervous system. We then quantify how carbon and nitrogen sources contribute to the tricarboxylic acid (TCA) cycle and glutamate synthesis across tissues. By measuring nutrient labeling in hemolymph, we link circulating hemolymph nutrients to tissue metabolism. ^13^C-metabolic flux analysis and direct contribution analysis are further used for quantitative evaluation. Finally, we apply this framework to evaluate how metabolic activities vary across a *yorkie*-driven gut tumor bearing fly model. We find an increased alanine contribution to malate in the gut and abdominal tissues, and gut-specific knockdown of the amino acid transporter *path* reduces tumor burden and extends lifespan. Together, these results establish a framework for quantitative spatial metabolic analysis in whole fly.

## RESULTS

### Building a Fly-Iso-Imaging workflow

To investigate the compartmentalized metabolic activities in the adult fruit fly, we established the Fly-Iso-Imaging (FISO) pipeline (**Fig. 1A**, see **Methods**). Building on the Iso-Imaging workflow (*14*), the pipeline begins with the administration of stable isotopic tracers to Oregon-R female flies, supplemented with Allura Red dye to account for potential feeding variability and help localize gut lumen (*31*). We then embedded tracer-fed flies in a fly-specific optimized cryosectioning medium and processed them for matrix-assisted laser desorption ionization mass-spectrometry imaging (MALDI-MSI). Under these optimized conditions, we consistently detected Allura Red dye signals in sagitally sectioned samples (**Fig. S1A-F, Table S1**). Additionally, the weighted fractional labeling (wFL) of input tracer reached approximately 20 to 30%, while TCA cycle intermediates, such as citrate and malate, reached 10 to 15% across whole flies fed with [U-^13^C_6_]glucose tracer (**Fig. S1G**). These ranges are consistent with those reported in the original mammalian Iso-Imaging workflow, supporting the feasibility of using our pipeline to evaluate spatial labeling of metabolites in the adult fruit fly.

**Figure 1.**
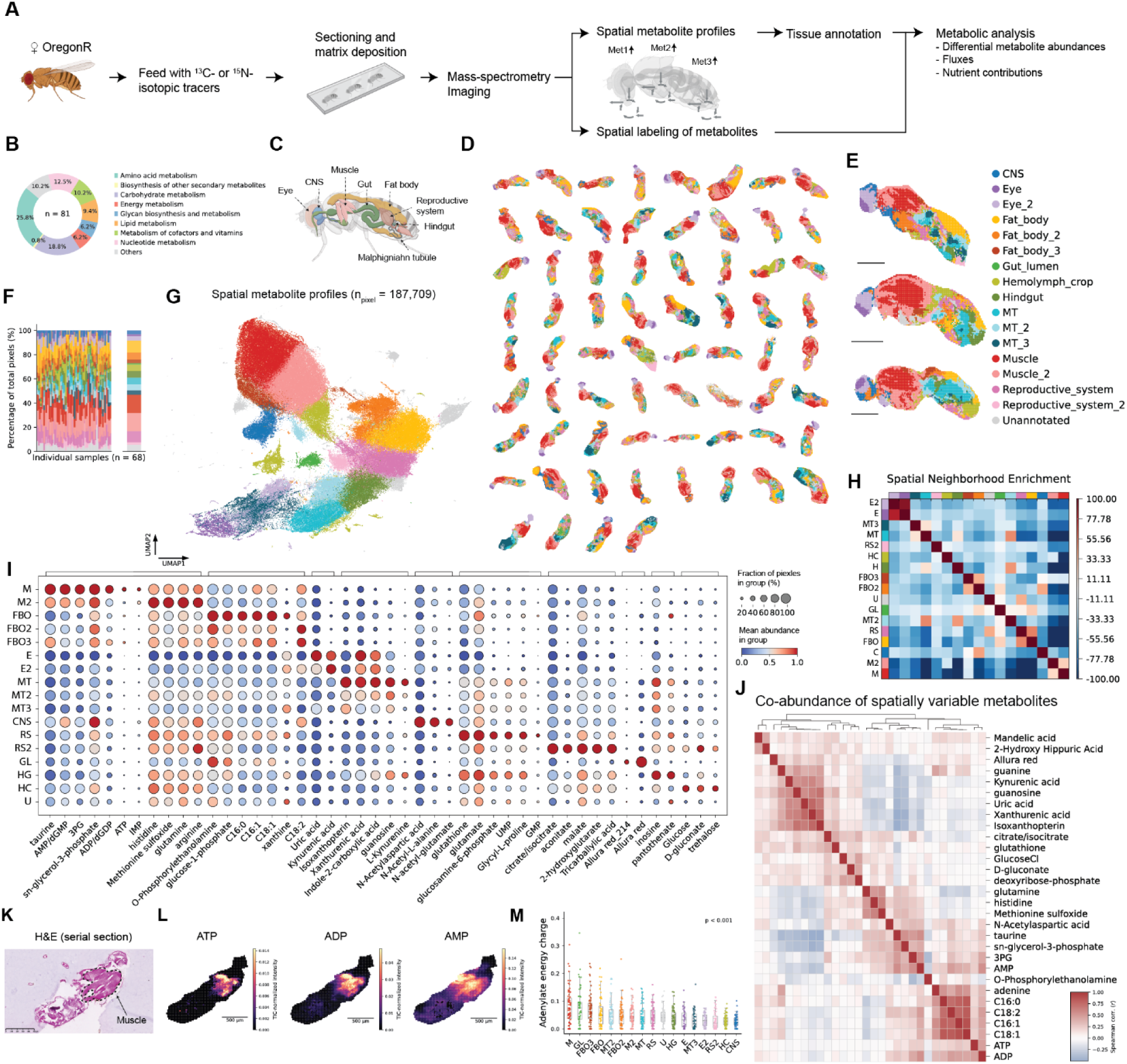
Establishment of a spatially resolved metabolic atlas of the adult fruit fly. **(A)** Schematic of the Fly-Iso-Imaging (FISO) workflow. **(B)** KEGG pathway classification of the metabolites included in the spatial metabolic atlas. **(C)** Schematic of major tissues in the adult fly sagittal section. **(D)** Spatial maps of tissue annotations across all 68 sagittal sections. **(E)** Representative tissue annotation maps from individual sections, colored as in the key. Scale bars, 500 µm. **(F)** Proportions of pixels assigned to each annotated tissue in individual sagittal sections and their mean across sections. **(G)** Uniform manifold approximation and projection (UMAP) of spatial metabolite profiles from 187,709 pixels across 68 flies, colored by annotated tissue. **(H)** Spatial neighborhood enrichment among annotated tissues. **(I)** Dot plot of tissue-enriched metabolites. Dot size indicates the percentage of pixels with nonzero metabolite signal within each annotated tissue, and color indicates mean normalized metabolite abundance. **(J)** Spearman correlation heatmap of the 30 metabolites with the highest Moran’s I values across all pixels. **(K)** Representative hematoxylin and eosin (H&E)-stained serial section corresponding to the spatial metabolite images. Muscle is outlined. **(L)** Representative spatial maps of ATP, ADP, and AMP intensity. Scale bars, 500 µm. **(M)** Quantification of adenylate energy charge (AEC), calculated as (ATP + 0.5×ADP)/(ATP + ADP + AMP), using TIC-normalized intensity of each adenylate. Tissues are ordered by mean AEC. Statistical significance was assessed by one-way ANOVA; the corresponding *P* value is shown. Abbreviations used in (H), (I), and (M): C, CNS; E, Eye; FBO, Fat_body; GL, Gut/lumen; HC, Hemolymph_crop; H, Hindgut; MT, Malpighian tubule; M, Muscle; RS, Reproductive_system; U, Unannotated. Numbered suffixes denote subclusters (e.g., FBO2, Fat_body_2; E2, Eye_2).

Next, we assessed the spatially resolved metabolite profiles. Quality control analysis revealed that approximately 80 metabolites consistently showed relatively high abundance and coverage **(Fig. S2A, B)**. Bulk liquid chromatography-tandem mass spectrometry (LC-MS/MS) also detected fifty of these metabolites, and their relative abundances correlated positively between the two platforms (r = 0.48), leaving most of the variance in MALDI intensity unexplained by bulk abundance (**Fig. S2C, D)**. To determine whether this metabolite coverage was sufficient to resolve anatomical regions, we performed unsupervised clustering analysis. We found that a single sagittal section, containing approximately 3,000 pixels, was sufficient to infer spatial clusters that closely corresponded to fruit fly anatomy (**Fig. S3A-F**). Moreover, metabolites enriched in each cluster showed higher overlap with metabolites enriched in anatomically corresponding dissected regions, independently measured by LC-MS/MS (*27*, *32*) (**Fig. S4**). Cluster 3 and the thorax were both enriched with glutamine, methionine sulfoxide, and adenosine monophosphate (AMP), while cluster 1 and the abdomen were enriched with uridine monophosphate (UMP), xanthine, and kynurenine. We quantified this overlap using the Jaccard similarity index for the top 20 enriched metabolites of each pair, and found that corresponding cluster–region pairs scored higher than non-corresponding ones, a descriptive comparison that was not tested against a permutation null. Together, these results indicate that spatially distributed metabolites from a single sagittal section can resolve major anatomical regions.

### Establishing a spatially resolved fly metabolic atlas

To build a spatial metabolic atlas, we scaled up the FISO pipeline to profile 68 biologically independent sagittal sections from Oregon-R female flies, fed with 10 ^13^C and 4 ^15^N nitrogen stable isotope tracers (**Table S2**). Because individual sagittal sections vary in exactly where they cut through the anatomy, we aggregated all datasets to capture this variation and increase the robustness of tissue region annotation. In total, we extracted 81 metabolites, including the exogenous marker Allura Red, across 187,709 pixels (see **Methods**). Similar to the single-section analysis, metabolites with higher abundances showed greater spatial coverage (Spearman ρ = 0.96, p < 0.0001; e.g., O-phosphorylethanolamine, taurine, and histidine). Kyoto Encyclopedia of Genes and Genomes (KEGG) classification analysis further showed that these metabolites spanned diverse pathways, with amino acid metabolism (26%), carbohydrate metabolism (19%), and nucleotide metabolism (13%) comprising the largest classes (**Fig. 1B**).

Next, we performed unsupervised clustering analysis on the spatial metabolite profiles (*33*) (see **Methods**). The resulting clusters formed spatial domains that broadly recapitulated the anatomical organization of the adult fly (**Fig. S5**-**8**). Using three criteria (i) overall enriched-metabolite profiles and their agreement with previously reported regional metabolite profiles (metabolic evidence), (ii) anatomical location and spatial organization (spatial evidence), and (iii) reproducibility across the 68 sections (consistency evidence) we annotated the clusters as eight regions: central nervous system (CNS), eye, fat body (FBO), gut/lumen, hindgut, Malpighian tubules (MT), muscle, and reproductive system (RS), with hemolymph/crop (a fluid and food-store class rather than a tissue) and unannotated pixels retained as two further labels; of these, gut/lumen contains luminal contents rather than cells alone, and all classes are provisional annotations of metabolite clusters (**Fig. 1C-E**, **Fig. S9–S15,** and **Tables S3-18**, see **Methods**). We retained subclusters with distinct enriched-metabolite profiles and spatial organization as separate subregions, such as the thoracic and head fat body regions labeled Fat_body_2 (FBO2) and Fat_body_3 (FBO3). We use these names throughout, including in the flux analysis. Overall, muscle accounted for the largest fraction of annotated pixels (30%), followed by fat body (19%) and Malpighian tubules (14%) (**Fig. 1F, G)**. To test whether the annotated tissues captured expected spatial relationships, we performed neighborhood enrichment analysis (*34*) (**Fig. 1H**). Physically adjacent regions had higher neighborhood scores (e.g., hindgut and Malpighian tubules), whereas anatomically separated regions of the same tissue type scored lower (e.g., abdominal versus thoracic fat body), indicating that the tissue annotations preserved spatial organization.

We next examined the metabolite profiles of the annotated tissues, which exhibited distinct enrichment patterns (**Fig. 1I**). Muscle was enriched for energy-related metabolites (e.g., ATP, ADP, and AMP), consistent with its high energetic demand, while the fat body class was enriched for lipid-associated metabolites (C16:0, C16:1, and C18:0). Eye and MT were enriched for purine-degradation products (uric acid, xanthine), with the eye also enriched for kynurenine-pathway and pteridine metabolites, the CNS for N-acetyl-aspartate, gut/lumen for Allura Red dye, and the reproductive system for nucleotides and TCA cycle metabolites. Co-abundance analysis of the most spatially variable metabolites further showed that functionally related metabolites grouped together, although co-abundance across whole-animal pixels also reflects shared tissue localization (**Fig. 1J**). Moreover, metabolite ratios, such as uric acid/xanthine, kynurenic acid/kynurenine, and glutamate/glutamine, further distinguished tissues (**Fig. S16A-I**). For instance, adenylate energy charge (AEC) (*35*), ratio calculated from the measured adenylate signals, was highest in muscle, followed by gut/lumen, whose pixels include luminal contents rather than cytoplasm alone, and Fat_body_3 (**Fig. 1K-M**).

Finally, we compared our tissue-enriched metabolite sets with dissected tissue metabolomics from FlyMet using the Jaccard similarity index score (*36*) (**Fig. S17A**). Consistently, the annotated tissues showed concordant enrichment patterns with their dissected counterparts: CNS with brain, muscle with muscle, and eye and MT with their respective dissected tissues (**Fig. S17B, C**). Together, the spatially resolved metabolic atlas reveals regional differences in metabolite abundance, and metabolite ratios in the adult fruit fly.

### Estimating tissue-level metabolic activities in the adult fruit fly

Because metabolite abundance or ratios do not directly report pathway activity (*37–40*), we next sought to quantify metabolic activities across tissues. We use metabolic activity in two distinct senses that should not be conflated: fractional ^13^C incorporation into a pathway intermediate, which depends jointly on flux, pool size and precursor enrichment. Later, we use fluxes fitted by ^13^C-metabolic flux analysis, which we estimate relative to the glucose input flux of the same tissue and are therefore partitioning ratios rather than absolute rates. Given the central role of glycolysis and TCA cycle in energy metabolism, we aimed to quantify their activities. We first analyzed the mass isotopomer distributions (MIDs) of sn-glycerol-3-phosphate (G3P), citrate, glutamate, and malate in a fly fed with [U-^13^C_6_]glucose (**Fig. 2A**). We used G3P to reflect glycolytic carbon incorporation, since M+3 G3P can arise only through triose phosphate, although its labeling also depends on G3P pool size and glycerolipid synthesis. We used citrate, glutamate, and malate labeling to reflect downstream TCA cycle metabolism. The observed isotopologue patterns were consistent with the expected patterns of [U-^13^C_6_]glucose catabolism, including M+1 labeling of TCA cycle intermediates that may reflect extensive CO₂ recycling *in vivo* (**Fig. S18A, B)** (*41*). Glucose-derived labeling of both G3P and TCA cycle metabolites was anatomically compartmentalized, with labeling highest in the thorax, followed by the head, and lowest in the abdomen, where unlabeled storage pools and luminal tracers also contribute (**Fig. S19A, B**). The evaluation of tissue-level labeling patterns further revealed that M+3 G3P, M+2 citrate, M+1 or M+2 glutamate and malate labeling varied significantly across tissues (one-way ANOVA, *P* < 0.05; *n* = 4), with muscle, CNS, and eye more highly labeled than other tissues (**Fig. 2B** and **Fig. S20**).

**Figure 2.**
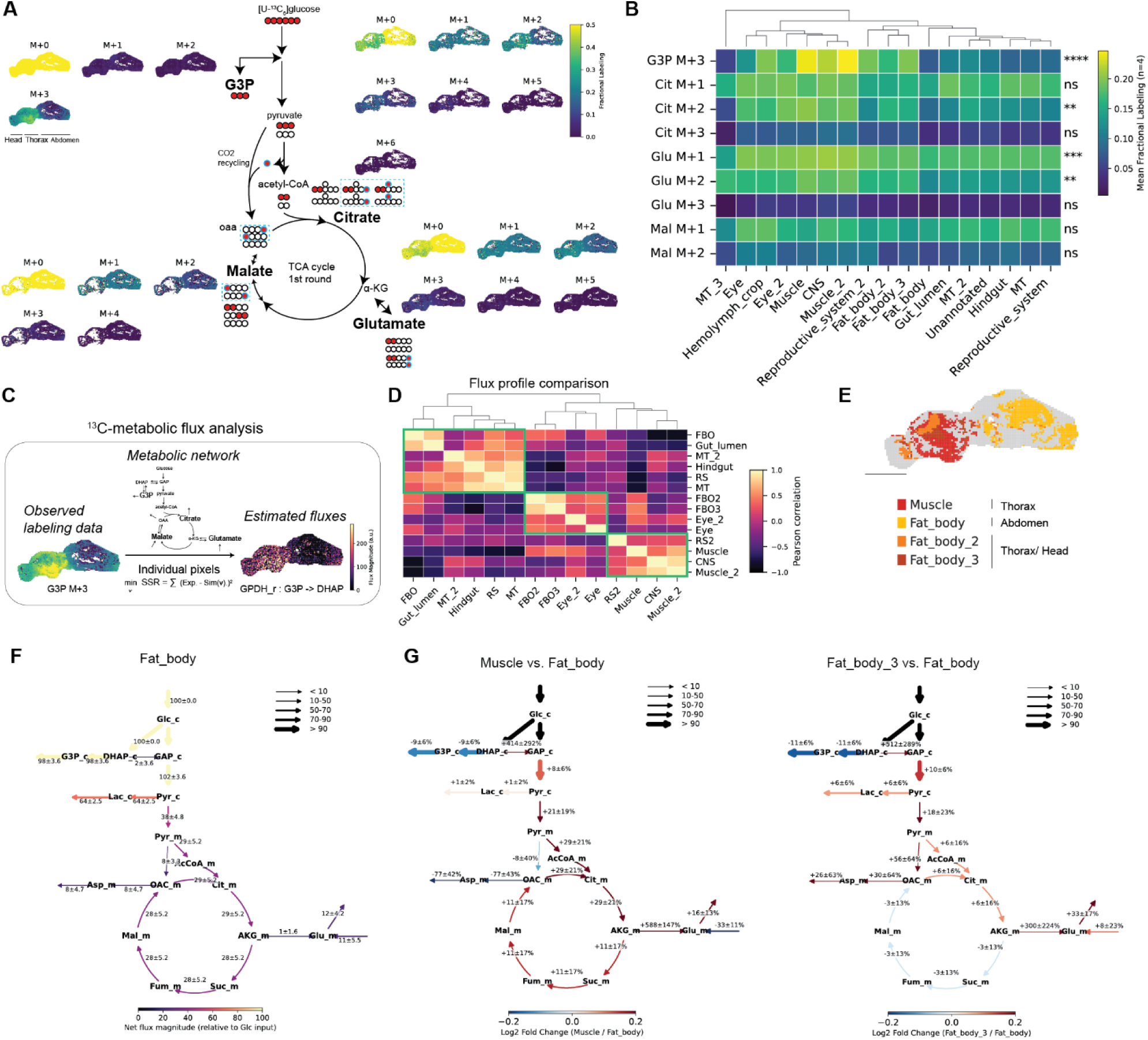
Estimation of spatially resolved metabolic fluxes in the adult fruit fly. **(A)** Representative spatial maps of isotopologues and expected labeling patterns of G3P, citrate, malate, and glutamate in flies fed [U-^13^C_6_]glucose. Color indicates isotopologue fractional labeling. **(B)** Mean fractional labeling for selected isotopologues across annotated tissues. One-way ANOVA was performed for each metabolite isotopologue. Significance is indicated as ns, not significant; \**P* < 0.05; \*\**P* < 0.01; \*\*\**P* < 0.001; \*\*\*\**P* < 0.0001. **(C)** Schematic of ^13^C-metabolic flux analysis using spatially resolved mass isotopomer distribution data. **(D)** Pairwise Pearson correlations between tissue-level flux profiles following z-score standardization. Tissues were ordered by hierarchical clustering. **(E)** Spatial map of the annotated fat body and muscle in the abdomen, thorax, and head. **(F)** Estimated flux map for Fat_body. Arrow direction indicates net flux direction, and arrow width indicates net flux magnitude relative to glucose input. Values indicate mean estimated flux ± SD across flies (n = 4); the SD is the between-fly scatter of the fitted values, not a confidence interval on the flux. **(G-H)** Comparison of estimated fluxes in **(G)** Muscle and **(H)** Fat_body_3 relative to Fat_body. Fat_body was used as the reference for both comparisons. Arrow color indicates the log_2_ fold change relative to Fat_body, and arrow width indicates net flux magnitude in Fat_body. Values indicate the mean percentage difference in flux ± SD across flies (n = 4).

Next, to estimate intracellular metabolic fluxes based on measured spatial labeling patterns of metabolites, we performed ^13^C-metabolic flux analysis (MFA) (*42*) (**Fig. 2C**). Flux vectors share the same stoichiometric constraints and a common normalization to glucose input. We compared simulated labeling patterns from the best-fitted fluxes with the experimentally measured patterns, and showed the minimized residual errors (**Fig. S21**). Moreover, correlation analysis across the 46 estimated fluxes revealed distinct flux profiles among tissues (**Fig. 2D**). Muscle correlated positively with thoracic or head fat body (FBO2, *r* = 0.30; FBO3, *r* = 0.48) and CNS (*r* = 0.39), but negatively with those of abdominal fat body (FBO, *r* = -0.52) and reproductive system (RS, *r* = -0.82). Abdominal fat body, on the other hand, correlated more strongly with abdominal tissues such as gut (*r* = 0.77) and RS (*r* = 0.49) than with thoracic or head fat body (*r* = 0.31 and 0.38, respectively). Spatial mapping showed that thoracic muscle was closer to FBO2 and FBO3 than to abdominal fat body, suggesting that similarities in flux profiles may also reflect spatial tissue organization (**Fig. 2E**).

To examine individual reactions underlying the tissue-level flux associations, we compared each tissue’s flux estimates. Fluxes through triose-phosphate isomerase (TPI), aspartate amino transferase, pyruvate carboxylase, and glutamate dehydrogenase (GLUD) were among those that differed by more than 100 % from the mean for the same reaction in at least one tissue (**Fig. S22A)**. Because thoracic muscle and abdominal fat body were the two most abundant annotated tissues in our dataset, we further compared their net fluxes by generating flux maps (**Fig. 2F-G**). Compared with abdominal fat body, muscle showed trends toward higher glycolytic flux, including conversion of glyceraldehyde 3-phosphate (GAP) to pyruvate (approximately 8% higher), and higher TCA cycle turnover, including mitochondrial isocitrate dehydrogenase (IDH_m) and mitochondrial malate dehydrogenase (MDH_m) fluxes (approximately 30% and 11% higher, respectively); fluxes are relative to glucose input, and the error bars show variability across flies (n = 4) rather than confidence intervals on the flux estimates. Consistent with their similar overall flux profiles, FBO2 and FBO3, which predominantly localized to thoracic and head regions, also showed higher flux through GAP to pyruvate conversion (approximately 10% and 11% higher, respectively) and IDH_m flux (approximately 6% and 10% higher, respectively) relative to abdominal fat body (**Fig. 2H** and **S22B**).

We further examined the two branch points in our metabolic network - the fate of triose phosphate in the cytosol and the source of glutamate in the mitochondria. In the cytosol, muscle, FBO2 and FBO3 showed higher TPI flux relative to G3P dehydrogenase (G3PDH) than abdominal fat body, suggesting that they direct triose phosphate toward glycolysis rather than toward G3P, a precursor for *de novo* synthesis of glycerolipids (**Fig. S22C**). In the mitochondria, abdominal fat body showed higher glutaminase (GLND) flux relative to GLUD, suggesting that its glutamate is supplied more from glutamine than from α-ketoglutarate within the modeled network (**Fig. S22D**). Although the network also included aspartate aminotransferase, its net flux was directed toward glutamate consumption and was therefore not included in this comparison of glutamate-forming routes. Altogether, these results reveal distinct estimated flux profiles across tissues and suggest that thoracic and head fat body resembles muscle more than fat abdominal fat body, with trends toward higher glycolytic and TCA cycle fluxes.

### Nutrient preference to fuel the TCA cycle across tissues in the adult fruit fly

In *Drosophila*, little is known about the contribution of circulating nutrients to TCA cycle substrates across tissues (*43*). Based on regional differences in glucose utilization, we hypothesized that tissues differ in their nutrient preferences for fueling the TCA cycle (**Fig. 3A**). To test this, we fed female Oregon-R flies with ^13^C-labeled nutrients and used labeling of the TCA cycle intermediate malate as a readout (**Fig. 3B**). This readout depends on the entry point: nutrients entering at pyruvate or oxaloacetate (glucose, fructose, lactate, alanine), or at alpha-ketoglutarate (glutamine, proline), label malate within one span of the cycle. By contrast, nutrients entering as acetyl-CoA (leucine, oleate) label malate only after condensation with oxaloacetate and a further turn, and are therefore expected to give lower malate labeling at equal flux. Glucose and fructose labeled malate more strongly in muscle, CNS, and eye than abdominal tissues, as raw labeling that is not normalized to hemolymph tracing enrichment. Among other nutrients, [U-^13^C_5_]glutamine labeled malate more strongly in the reproductive system and hindgut, whereas [U-^13^C_5_]proline labeled it most strongly in the CNS and eye (**Fig. 3C, D**). [U-^13^C_3_]alanine also labeled it in MT, hindgut, and muscle. Citrate, leucine, and oleate labeled it across tissues, although their lower observed labeling cannot be separated from limited uptake at the concentrations fed (**Table S2**). These panels report weighted fractional labeling of malate rather than contribution: they are not normalized to the hemolymph enrichment each tracer reached and do not yet account for interconversion among circulating nutrients (**Fig. S23**), both of which are addressed below.

**Figure 3.**
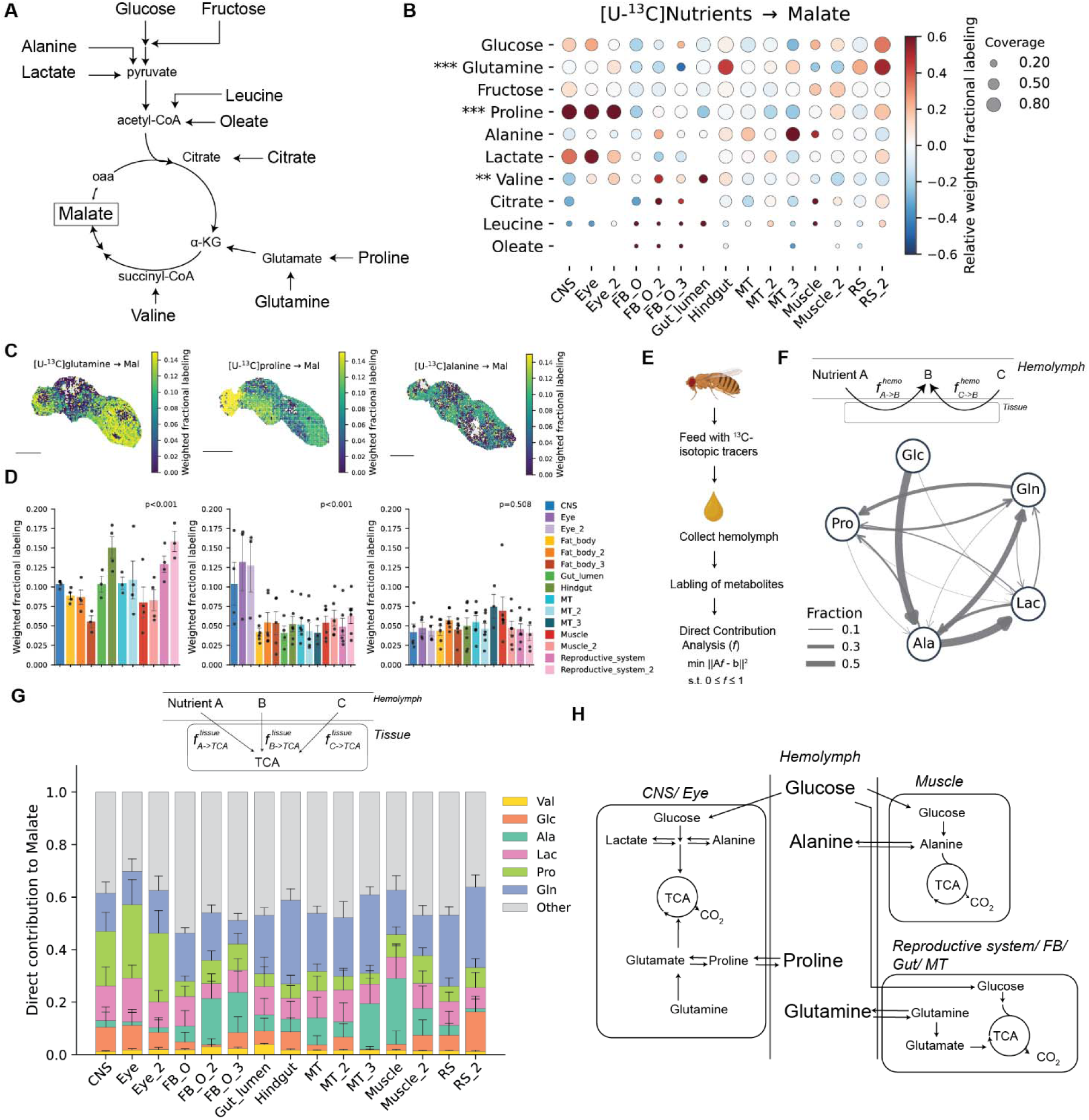
Nutrient preferences for fueling the TCA cycle across tissues in the adult fruit fly. **(A)** Schematic of pathways from input nutrients to the TCA cycle metabolites. **(B)** Weighted fractional labeling (wFL;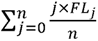) of malate from the indicated ^13^C-labeled nutrients across annotated tissues. Color indicates the relative deviation of wFL from the median across tissues for each nutrient; dot size represents the mean coverage fraction. One-way ANOVA was performed across tissues; \*\**P* < 0.01; \*\*\**P* < 0.001. **(C)** Representative spatial maps of wFL of malate from [U-^13^C_5_]glutamine, [U-^13^C_5_]proline, and [U-^13^C_3_]alanine. Scale bars, 500 µm. **(D)** Quantification of wFL of malate from the spatial map shown in C. Bars indicate mean ± SEM and points represent biological replicates. Tissues with at least three biological replicates were included. One-way ANOVA was performed, with the corresponding *P* values shown. **(E)** Workflow for measuring hemolymph nutrient interconversion and performing direct contribution analysis. **(F)** Network of estimated direct fractional contributions of circulating nutrients to other nutrients in hemolymph. Edge width indicates fractional contribution. **(G)** Estimated fractional contributions of circulating nutrients to malate across annotated tissues. Stacked bars indicate mean fractional contributions from 5,000 Monte Carlo iterations, in which hemolymph and tissue labeling were resampled from their means and SEMs, with error bars indicating SD; this reflects uncertainty propagated from the measured labeling values, not uncertainty in the selected nutrient set. **(H)** Model of tissue preference for circulating nutrient utilization.

As nutrients are readily exchanged among tissues (*44*, *45*), we calculated the extent of interconversion among circulating nutrients in hemolymph, analogous to human blood (*44*). After feeding flies with ^13^C-labeled tracers, we extracted hemolymph and measured ^13^C labeling of both input and other circulating nutrients (**Fig. 3E**). While glucose labeled other circulating nutrients most extensively, alanine also labeled circulating nutrients, including lactate, glutamine, and proline, to a substantial degree (**Fig. S23A, B**). Direct contribution analysis on circulating nutrients indicated that alanine, derived from glucose, contributes to circulating lactate and glutamine (**Fig. 3F, and S23C**). Because alanine aminotransferase and lactate dehydrogenase both operate at near equilibrium (*46*), these estimates report carbon exchange among the alanine, pyruvate and lactate pools and do not on their own establish net carbon transfer. Together, these results suggest that alanine serves as a key circulating carbon carrier among the nutrients measured here; we did not include trehalose, one of the main circulating sugars in the adult fly (*30*), in the contribution model.

Next, because circulating nutrients supply carbon to tissue energy metabolism through the TCA cycle (*47*), we quantified their contributions to TCA cycle metabolites across tissues (**Fig. 3G, and S24A, B**). The analysis apportioned glucose-derived carbon to the TCA cycle via circulating alanine in muscle, fat body, and the MT. Because pyruvate formed by local glycolysis and pyruvate formed from imported alanine are isotopically indistinguishable, this routing is a model apportionment rather than a direct measurement. Glutamine was a major direct contributor in the gut, whose pixels include luminal contents and associated microbiota, the MT, and the reproductive system, while proline was a major contributor in the CNS and eye. Altogether, these results reveal preferential nutrient utilization across tissues, when each nutrient is supplied alone after a 4-hr fast (see **Methods** and **Table S2**), with glucose, alanine, glutamine, and proline serving as distinct direct and indirect TCA contributors (**Fig. 3H**).

### Nitrogen sources for glutamate and its derivatives

Glutamate is at the intersection of carbon and nitrogen metabolism, accepting amine nitrogen from circulating amino acids through transamination and dehydrogenase reactions (*48*) (**Fig. 4A**). To evaluate the nitrogen source of glutamate and its related nitrogenous products, we fed flies four different ^15^N-labeled amino acids and quantified the ^15^N-labeling of glutamate, N-acetyl-aspartate (NAA), and glutathione (**Fig. 4B-D and S25A-C**). All four nitrogen sources labeled glutamate across the whole body without detectable tissue specificity; one-way ANOVA across annotated tissues was non-significant for every donor (P = 0.997 for proline, P = 1.000 for alanine and glutamine; **Fig. 4B**). Among the four nitrogen donors, [^15^N]alanine labeled glutamate nitrogen most substantially in raw ^15^N-labeling; the hemolymph enrichment of each tracer was measured later in separate experiments.

**Figure 4.**
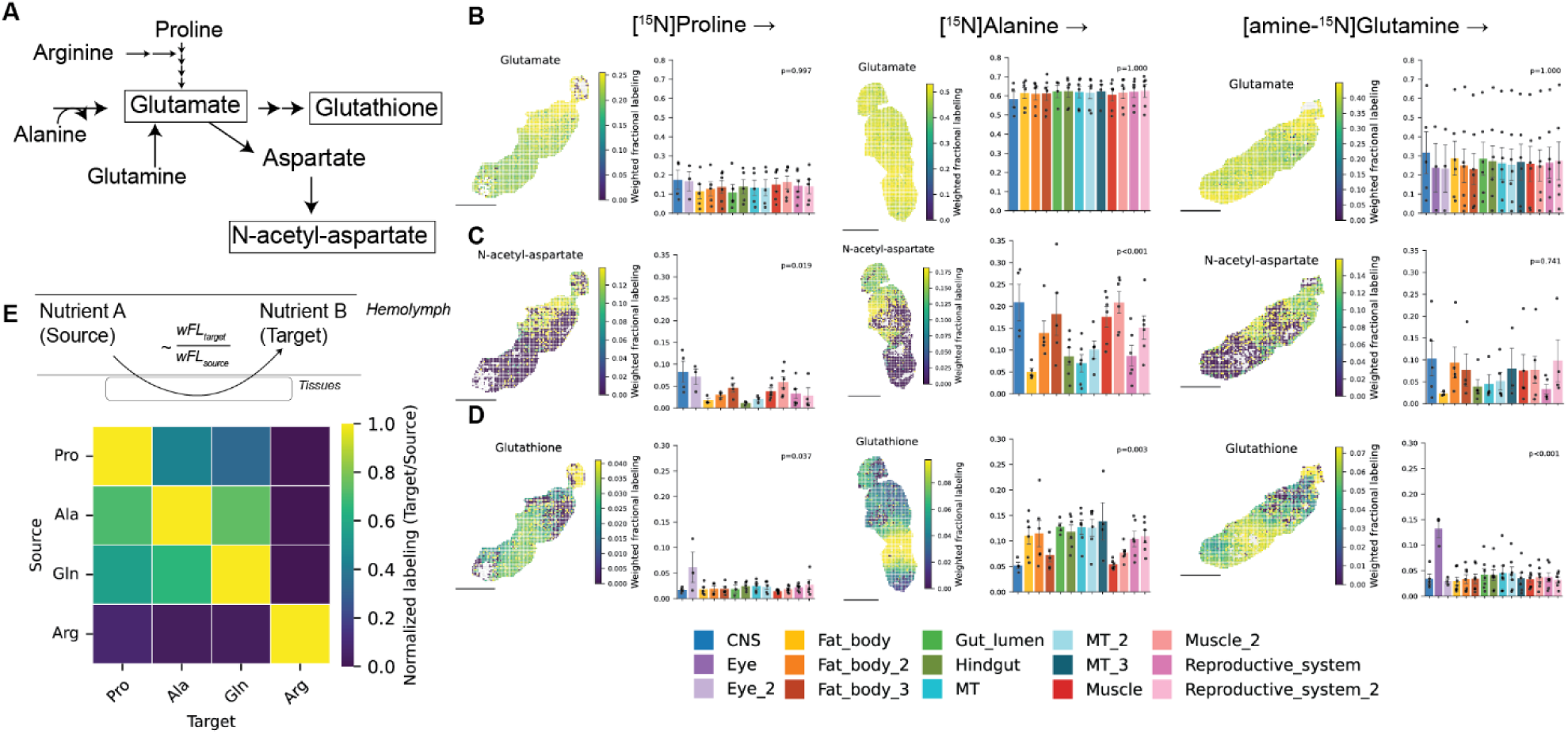
Nitrogen sources for glutamate and its derivatives. **(A)** Schematic showing the amino acid nitrogen contributions to glutamate, N-acetyl-aspartate, and glutathione. **(B-D)** Representative spatial maps and quantification of weighted fractional labeling (wFL) from the indicated ^15^N-labeled amino acid tracers in **(B)** glutamate, **(C)** N-acetyl-aspartate, and **(D)** glutathione. For the ^15^N tracers the wFL denominator is the number of nitrogen atoms in the metabolite (one for glutamate and N-acetyl aspartate, three for glutathione). Each map has an independently scaled color bar; labeling values should be read from the corresponding axis rather than comparing colors between maps. Bars show mean ± SEM, and points represent biological replicates. Tissues with at least three biological replicates were included. One-way ANOVA was performed across tissues, with the corresponding *P* values shown. Scale bars, 500 µm. **(E)** Heatmap of normalized ^15^N labeling of circulating nutrients in the hemolymph after feeding the indicated ^15^N-labeled amino acid tracers. Color indicates wFL of target circulating nutrients divided by the wFL of source nutrient in hemolymph.

In contrast to glutamate, ^15^N-labeling to NAA and glutathione were more tissue specific. [^15^N]proline and [^15^N]alanine labeled NAA nitrogen predominantly in the head and thorax, with the CNS among the most strongly labeled tissues (one-way ANOVA across tissues, P=0.019 and P < 0.001, respectively), whereas glutamine showed no significant tissue difference (P = 0.741) (**Fig. 4C**). Proline and glutamine labeled glutathione nitrogen most strongly in the eye, while alanine labeled most in the abdominal tissues such as gut and MT (**Fig. 4D**). Given that [^15^N]alanine labeled glutamate nitrogen most extensively across tissues, we asked whether alanine similarly labeled other circulating nutrients. After feeding flies ^15^N-labeled tracers, we extracted hemolymph and measured ^15^N-labeling of both the input tracer and the other circulating nutrients. Indeed, [^15^N]alanine labeled other circulating nutrients more extensively than glutamine, proline, or arginine (**Fig. 4E and S25D-F**). Together, these results show that alanine nitrogen exchanges extensively with other circulating amino acids and reaches NAA and glutathione in a spatially compartmentalized manner; because transamination is reversible, we did not determine net nitrogen transfer.

### Systemic rewiring of alanine utilization in a *Drosophila* gut-tumor model

To determine whether alanine utilization changes in disease, we applied the FISO framework to a *Drosophila* gut-tumor model. Expression of an activated form of *yorkie* (*yki*), the *Drosophila* ortholog of human *YAP1*, in intestinal stem cells drives gut over-proliferation and wasting of peripheral organs (*49–51*). After feeding [U-^13^C_3_]alanine to flies bearing yki-activated gut tumors (hereafter, *yki* flies), we measured both spatially resolved metabolite profiles and labeling patterns of malate (**Fig. 5A, B**). At the whole-fly level, 27 metabolites were significantly altered (p_adj_ < 0.05 and |log_2_FC| > 1), with 10 decreased and 17 increased (**Fig. S26A**). The most significantly decreased metabolites included ADP, glutamate, glutathione, and free fatty acids, while trehalose and allantoin were among the most increased metabolites, consistent with previous metabolomics studies (*52*, *53*).

**Figure 5.**
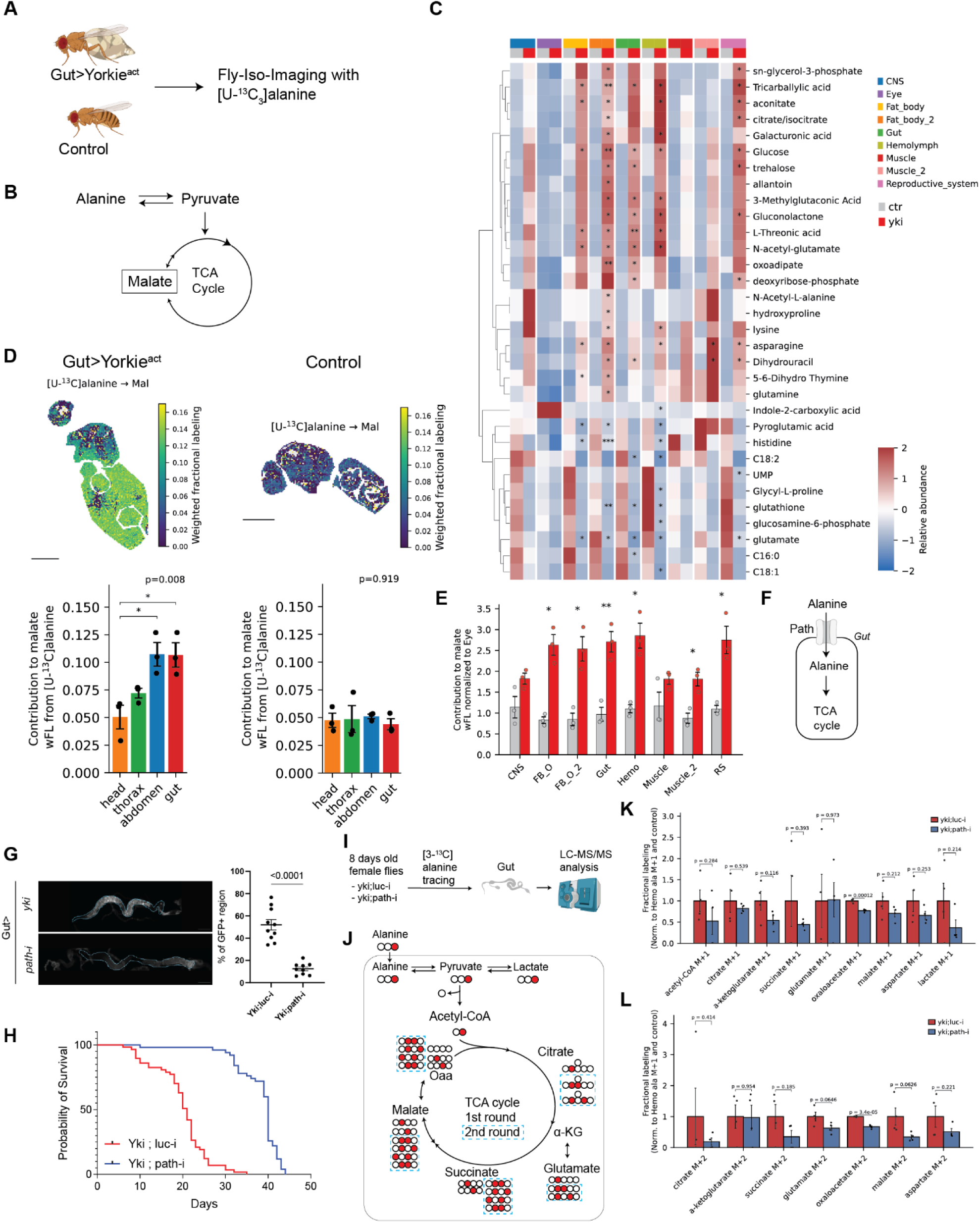
Systemic rewiring of alanine utilization in a *Drosophila* gut-tumor model. **(A)** Experimental design comparing control and *yki* flies analyzed by Fly-Iso-Imaging after feeding [U-^13^C_3_]alanine. **(B)** Schematic of alanine contribution to malate through pyruvate and the TCA cycle. **(C)** Relative metabolite abundance across annotated tissues in control and *yki* flies. Significantly perturbed metabolites (false discovery rate, FDR < 0.05) in at least one tissue are shown. Asterisks indicate significance for comparisons between genotypes within each tissue (\**P* < 0.05, \*\**P* < 0.01, \*\*\**P* < 0.001). **(D)** Representative spatial maps and quantification of malate weighted fractional labeling (wFL) from [U-^13^C_3_]alanine in *yki* and control flies. Bars show mean ± SEM and dots represent biological replicates (n = 3 flies per genotype). Differences among regions were assessed by one-way ANOVA separately within each genotype; corresponding *P* values are shown; asterisks mark pairwise comparisons against the head within *yki* flies (\**P* < 0.05). Scale bars, 500 µm. **(E)** Alanine contribution to malate across annotated tissues in control and *yki* flies, expressed as the malate wFL of each tissue divided by that of the eye in the same fly; the eye is therefore fixed at one by construction and is not shown. Bars show mean ± SEM and dots represent biological replicates (n = 3). Significance was assessed for each tissue between genotypes (\**P* < 0.05, \*\**P* < 0.01). Because every tissue is referenced to the eye, this panel reports the distribution of labeling between tissues and not its absolute magnitude. **(F)** Schematic of gut-specific knockdown of *path* in *yki* flies. **(G)** Representative gut images and quantification of the percentage of gut area positive for GFP driven by *esg*-GAL4 following gut-specific *path* knockdown. Bars show mean ± SEM and dots represent individual guts. A two-sided unpaired Student’s *t*-test was performed, with the corresponding *P* value shown. **(H)** Survival curves of *yki;luc-i* and *yki;path-i* flies. **(I)** Experimental design for [3-^13^C]alanine tracing experiments for *yki;luc-i* and *yki;path-i* flies. **(J)** Schematic of the expected ^13^C-labeling patterns generated from [3-^13^C]alanine in the TCA cycle **(K-L)** Fractional labeling of **(K)** M+1 and **(L)** M+2 isotopologues of the indicated gut metabolites in *yki;luc-i* and *yki;path-i* flies after [3-^13^C]alanine feeding, corrected for natural isotope abundance, divided by the hemolymph alanine M+1 fractional labeling of the same fly, and normalized to the *yki;luc-i* control. Bars show mean ± SEM and dots represent biological replicates (n = 4). A two-sided unpaired Student’s *t*-test was performed, with the corresponding *P* values shown.

We next performed clustering analysis and tissue annotations (**Fig. S26B-D**, and see **Supplementary Text**). Consistent with previous observations, hemolymph clusters accounted for nearly 30% of all pixels in sectioned *yki* flies, about three times higher than the control, although the atlas could not separate hemolymph from crop contents (**Fig. S26E**). Moreover, many tissue-enriched metabolites identified in the fly metabolic atlas were found in the same annotated tissues in both control and *yki* flies, supporting the resulting tissue annotations (**Fig. S26F-H**). At the tissue level, metabolite perturbations varied across regions (**Fig. 5C**). The eye was largely unaffected, whereas free fatty acid–and glutathione-associated metabolites were decreased in the CNS, abdominal fat body, gut, and reproductive system. Muscle showed the strongest depletion of histidine in the tissue panel. Conversely, sn-glycerol-3-phosphate, trehalose, and allantoin were elevated across most tissues, except the eye (**Fig. 5C**); adenine nucleotides were among the metabolites decreased at the whole-body level (**Fig. S26A**) but were not retained in the tissue-level panel.

Because alanine substantially labeled circulating amino acids and TCA cycle metabolites in wild-type flies, we next performed [U-^13^C_3_]alanine tracing to evaluate its contribution to TCA cycle metabolites across anatomical regions (**Fig. S27A**). Compared with controls, the alanine contribution to malate was significantly elevated in the abdomen and gut of *yki* flies relative to that in head (three flies per genotype; we did not normalize imaging values to hemolymph alanine enrichment) (**Fig. 5D and S27B**). At the annotated tissue level, alanine-derived malate labeling, relative to the eye within each genotype, was approximately threefold higher in *yki* flies than in controls in both fat body clusters, the gut, the hemolymph compartment, and the reproductive system, and approximately two-fold higher in the second muscle cluster, whereas the CNS and muscle did not differ significantly between genotypes (**Fig. 5E**). The increased contribution in fat body is consistent with our earlier finding that alanine serves as a substrate for trehalose synthesis, analogous to mammalian gluconeogenesis (*53*), although malate labeling alone cannot separate alanine oxidation from gluconeogenic export.

We next asked whether decreasing alanine utilization in the gut could ameliorate the disease phenotype. Using the Gal4-UAS system, we knocked down *path*, a SLC36-family proton-coupled amino acid transporter that transports alanine and, to a lesser extent, glycine and proline, specifically in the gut (*54*) (**Fig. 5F**). Gut-specific knockdown of *path* with a RNAi line decreased tumor burden, as reflected by a more than 50% reduction in *esg*-driven green fluorescent protein (GFP) signal, and extended lifespan (**Fig. 5G, H**). Furthermore, we performed [3-^13^C]alanine tracing experiments in *yki* flies with gut-specific *path* knockdown (*yki;path-i*) or the luciferase-RNAi control (*yki;luc-i*) **(Fig. 5I)**. We expected decreased M+1 and M+2 fractional labeling of TCA cycle metabolites in the gut of *path*-knockdown *yki* flies **(Fig. 5J)**. Indeed, we observed a trend of decrease in labeling of TCA cycle metabolites **(Fig. 5K,L** and **Fig. S28A-C)**. Compared with *yki;luc-i* control flies, normalized M+1 fractional labeling of acetyl-CoA, citrate, succinate, and malate decreased by approximately 48, 18, 56, and 29%, respectively, in *yki;path-i* flies (**Fig. 5K**). Similarly, M+2 labeling of citrate, succinate, malate, which requires more than one turn of the cycle or carbon recycling, decreased by approximately 81, 65, and 66%, respectively (**Fig. 5L**). As acetyl-CoA carries at most one ^13^C from [3-^13^C]alanine, we do not report its M+2 labeling fraction.

In parallel, we fed *yki* flies β-chloroalanine, an inhibitor of alanine aminotransferase, the enzyme that interconverts alanine and pyruvate (*55*) (**Fig. S29A**). β-chloroalanine treatment decreased tumor burden in *yki* flies, as reflected by reduced GFP signals (**Fig. S29B**). [3-^13^C]alanine tracing similarly revealed a trend of decrease in labeling of TCA cycle metabolites, particularly M+1 labeling (**Fig. S30-31**). Specifically, normalized M+1 fractional labeling of acetyl-CoA, citrate, succinate, and malate decreased by approximately 57, 44, 55, and 20%, respectively. Note that, as β-chloroalanine is a mechanism-based inhibitor of pyridoxal-5′-phosphate–dependent enzymes and is not selective for alanine aminotransferase, this result is supporting rather than independent evidence. Altogether, these results show that our framework can resolve systemic metabolic perturbations in a *Drosophila* gut-tumor model, revealing region-specific rewiring of alanine contribution to TCA cycle metabolites and its link to tumor burden.

## DISCUSSION

Metabolism is highly compartmentalized at the organismal level, yet direct assessment of metabolic activity and nutrient use across the whole adult *Drosophila* has remained challenging. Here, using spatial isotope tracing and flux analyses, we established the Fly-Iso-Imaging (FISO) framework, building a whole-body spatial metabolic atlas of the adult fruit fly and quantifying metabolic activity and nutrient utilization in a spatially resolved manner. By preserving anatomical context, our framework resolves regional differences in relative metabolite abundance and nutrient use that are difficult to distinguish in dissected samples.

We found compartmentalization of proline utilization. In several insects, including the Colorado beetle, mosquitoes, and the tsetse fly, proline has been reported to serve as a fuel for muscle energy metabolism (*56*, *57*). In resting *Drosophila* fed proline as the only nutrient after a fast, however, we found that proline fueled the TCA cycle more strongly in head regions, including the brain and eye, than in thoracic muscle; whether flight activates proline oxidation in fly flight muscle, as reported for these insects, was not tested here. Even within the thorax region, the proline contribution to malate was higher in the annotated nervous system than in the muscle. Measurements of dissected thoraces cannot readily separate these contributions because the samples contain both muscle and nonmuscle tissues. These findings highlight the importance of resolving metabolic activities spatially to distinguish tissue-specific nutrient utilization that would otherwise be obscured by bulk tissue measurements.

We also found that circulating alanine carries glucose-derived carbon and participates in nitrogen exchange across tissues. Alanine-mediated inter-tissue metabolite communication has been described in *Drosophila*, where glucose-derived alanine is transferred from glia to neurons and converted back to pyruvate to support mitochondrial metabolism (*58*, *59*). Our results indicate that glucose also fueled the TCA cycle through circulating alanine in subsets of muscle, Malpighian tubules, and the fat body, being the functional analog of mammalian liver and adipose tissue. This extends the role of alanine beyond brain metabolism and the classical mammalian glucose–alanine cycle (*60*). Additionally, our results deviate from mammalian systems, in which circulating lactate serves as a major glucose derived carbon source for the TCA cycle (*47*). These comparisons suggest that the relative roles of circulating carbon carriers may vary with species and physiological conditions.

Beyond normal physiology, our framework revealed regional changes in nutrient utilization associated with gut tumors. In a *Drosophila* gut-tumor model, we found that tumor growth was accompanied by changes in metabolite profiles across tissues and by an increased alanine contribution to malate in the gut and surrounding abdominal tissues. These findings link the regional use of circulating nutrients with a model of tumor–host metabolic communication. Previous studies have implicated alanine in host gluconeogenesis in tumor-bearing mice, zebrafish, and flies (*53*, *55*, *61*), and in supplying carbon to pancreatic tumor cells (*62*). Given that alanine serves as both an oxidative carbon source and a major systemic nitrogen donor in our study, its increased utilization may help meet the altered carbon and nitrogen demands of both tumor and host tissues. Further studies are needed to determine how alanine-derived carbon and nitrogen contribute to tumor growth and how other carbon and nitrogen sources contribute to this gut-tumor model.

Finally, this framework has potential applications beyond the findings reported here. It could be adapted to investigate compartmentalized nutrient use in other systems, including vector–host interactions. MSI has been used to map lipid distributions in *Ixodes* ticks and adjacent guinea pig skin at the bite site (*63*), and our approach could help further delineate nutrient use and metabolic activity at such interfaces. Our framework also provides direct experimental measurements that could be used to evaluate predictions and constrain tissue-specific genome-scale metabolic models of *Drosophila* (*27*). Moreover, it could be integrated with spatial transcriptomic data in *Drosophila* to link metabolic gene expression with metabolic activity (*18*).

This study has limitations. First, we gave each tracer as a single purified nutrient after a fast (**Table S2**), so the labeling patterns report nutrient use under that feeding state rather than on a mixed diet. The atlas pools 68 flies across 14 such diets. Second, tumor-model comparisons rest on three biological replicates per genotype, so tissue-level differences that did not reach significance may reflect limited power rather than absence of effect. Third, we assigned tissue annotation to metabolite clusters by enriched-metabolite profiles, location and reproducibility rather than by co-registered histology, so the classes are provisional, gut/lumen and hemolymph classes contain non-tissue material, and boundaries are not exact. Fourth, we fit the metabolic flux analysis to one tracer and five measured mass isotopomer distributions per tissue. The fluxes are relative to glucose input, several branch fluxes are not uniquely identified by these data, and we could not measure tissue-specific net uptake or release rates *in vivo*. Fifth, the spatial resolution of current mass-spectrometry imaging does not reach the single-cell level, so annotated regions contain multiple cell types. Finally, we avoided fixation to preserve metabolite pools, so some regions were susceptible to deformation or localized metabolite leakage, particularly near the outer edge of the animal. Advances in imaging resolution, sample preparation and flux analysis will help address these limitations.

Together, this study establishes a framework for quantitative spatial metabolic analysis in the adult fruit fly. We anticipate that this approach will enable systematic investigation of metabolic compartmentalization across physiological and pathological states in *Drosophila*, and across other species.

## MATERIALS AND METHODS

### Fly stocks and maintenance

All flies were reared at 25 °C and 60% humidity with a 12-hour light/dark cycle on standard laboratory food. Standard laboratory food is made as follows: 12.7 g/liter deactivated yeast, 7.3 g/liter soy flour, 53.5 g/liter cornmeal, 0.4% agar, 4.2 g/liter malt, 5.6% corn syrup, 0.3% propionic acid, and 1% Tegosept/ethanol. The food was replaced every two days. For mass-spectrometry imaging experiments, five-day-old Oregon-R (OreR) female flies and eight-day-old female *yorkie*-activated gut tumor flies (*yki* flies) were used, unless otherwise noted in the main text. Bloomington Drosophila stock center: UAS-path RNAi (BDSC: 64029). UAS-luc RNAi (BDSC: 31603). Laboratory stocks: *w^1118^* and OreR.

### Gut tumor induction

Crosses were set up with *esg*-GAL4, *tub*-GAL80^ts^, UAS-GFP (hereafter *esg*^ts^) and UAS-*yki*^3SA^ at 18 °C to inactivate GAL4, as previously described (*49*). Following eclosion, adult flies were collected and kept at 18 °C. two- to five-day-old adult progeny were placed at 29 °C to induce the transgenes. Progeny from a cross between *esg*^ts^ and *w*^1118^ were used as controls. The food was replaced every two days. For the RNAi of *path* in *yki*^3SA^-tumor guts, UAS-*path* RNAi or UAS-*luc* RNAi (control) was crossed to *esg*-GAL4, *tub*-GAL80^ts^, UAS-GFP/CyO; UAS-*yki^3SA^/TM6B* at 18 °C. Gut morphology was measured after switching to 29 °C for 8 days. For inhibitor treatment, flies were transferred onto food containing inhibitors at day 0 (simultaneously with tumor induction). L-β-Chloroalanine hydrochloride (MedChemExpress, HY-107373A) was used as an alanine aminotransferase inhibitor at a final concentration of 250 µM in the food. Control flies received the same food prepared with the vehicle alone.

### Fly-Iso-Imaging workflow

Fly-Iso-Imaging was adapted from the previously described Iso-Imaging method for application to whole adult *Drosophila* (*14*). In brief, five-day-old adult female OreR flies or eight-day-old female *yki* flies were fed stable isotopic tracers together with Allura Red dye. Whole flies were embedded in a cryosectioning medium, sectioned, coated with matrix, and analyzed by matrix-assisted laser desorption ionization mass-spectrometry imaging (MALDI-MSI). Metabolite and isotopologue intensities were extracted using IsoScope (*14*). Metabolite intensity data were used for unsupervised clustering and tissue annotation. Mass isotopomer distribution (MID) was calculated and evaluated at the anatomical or annotated tissue level. ^13^C-metabolic flux analysis was performed on spatially resolved MID data at each pixel. Detailed methods for each step are described below.

### 13C-isotopic labeling experiments

^13^C-isotopic tracing experiments in fruit flies were performed as previously described (*27–29*, *64*). At around 9:00 AM, about 20 flies were transferred to an empty vial containing Whatman paper (MilliporeSigma, WHA10010155) with deionized water. After four hours, the flies were transferred to a new vial containing Whatman paper with ^13^C- or ^15^N-labeled tracers with 10 mM Allura Red dye (Sigma-Aldrich, 458848). Tracer concentrations are listed in **table S2**. After four to five hours of feeding, flies were briefly placed on a CO_2_ pad and quickly placed into the cryomold for subsequent cryosectioning, or the thorax, head, or abdomen was dissected for bulk liquid chromatography-tandem mass spectrometry (LC-MS/MS) analysis. The isotopic pseudo-steady state of TCA cycle metabolites over this time scale was confirmed in adult fruit flies (*28*). For [3-^13^C]alanine (Cambridge Isotope Laboratories, CLM-117) tracing experiments in *yki;luc-i* and *yki;path-i* female flies, the same feeding procedure was used. Guts were dissected from 15 flies per replicate for LC-MS/MS analysis.

### Tissue preparation

Embedding medium was prepared by combining 10% gelatin (Sigma-Aldrich, G1890) and 5% carboxymethyl cellulose (CMC; Sigma-Aldrich, C5678). The cryomold (Sakura, 4566) was half filled with the embedding medium. About nine flies were placed onto the mold in the sagittal orientation. The rest of the cryomold was filled with the medium, and the mold was placed in a 100% ethanol bath with dry ice. Once the embedding medium turned white and solidified, the cryomold was covered with aluminum foil. Frozen samples in the cryomold were stored intact at −80 °C until matrix-assisted laser desorption ionization mass-spectrometry imaging (MALDI-MSI) analysis. For each sample, ∼ 12-µm-thick sections were collected on a cryostat (Leica, CM3050S). The CT and OT temperatures were set at -22 °C. Sections for MALDI-MSI were thaw-mounted on indium tin oxide (ITO)-coated glass slides (Sigma-Aldrich, 576352) and desiccated under vacuum for 15 min.

### Matrix coating

Desiccated tissue sections mounted on ITO-coated glass slides were sprayed using an HTX TM sprayer (HTX Technologies) with 10 mg mL^−1^ N-(1-naphthyl)ethylenediamine dihydrochloride (NEDC; Sigma, 222488) dissolved in 70%:30% (v/v) methanol:water, followed by filtration (0.20-µm pore size). The sprayer temperature was set to 70 °C, with a flow rate of 0.1 mL min^−1^, velocity of 1,000 mm min^−1^, track spacing of 2 mm, pressure of 10 psi, and a gas flow rate of 3 L min^−1^. Ten passes of the matrix were applied to slides with 10 s of drying time between each pass.

### MALDI mass spectrometry

For MALDI Fourier transform ion cyclotron resonance (FT-ICR) mass spectrometry, matrix-coated slides were loaded into a slide adaptor (Bruker Daltonics) and then into a solariX XR FT-ICR mass spectrometer equipped with a 9.4 T magnet (Bruker Daltonics). The resolving power was 120,000 at *m/z* 500. Mass accuracy was calibrated on tissue within 1 ppm before starting the run. *m/z* 124.0068 (taurine), *m/z* 133.0136 (malate) and *m/z* 145.0611 (glutamate) were used as lock masses during the run. The minimum intensity threshold for peak picking during acquisition was set at 5 × 10^3^. The laser focus was set to small and the x–y raster width was set to 20 µm, using Smartbeam-II laser optics. A spectrum was accumulated from 200 laser shots at 1,500 Hz. Data collection was performed using Bruker FlexImaging software (Bruker Daltonics).

### MALDI-MS/MS

All MS/MS spectra were acquired in negative ionization mode using a MALDI source coupled to a quadrupole mass analyzer on the same 9.4 T solariX XR FT-ICR that was used throughout the study. Fresh-frozen tissue cryosections were coated with NEDC matrix and analyzed directly on-tissue without prior extraction. For each target metabolite, the [M−H]^−^ precursor ion was isolated using a quadrupole isolation window of 0.1–0.4 *m/z* to confirm the presence of only the desired precursor and to exclude isobaric interferences and ^13^C isotope contributions from neighboring species. Fragmentation was achieved by CID in the collision cell. The collision RF amplitude was set between 900 and 1,300 Vpp and the collision energy between 7 and 12 V, optimized per metabolite to maximize informative fragmentation while retaining residual precursor signal. Fragment ions were detected in the quadrupole mass analyzer. Experimental MS/MS spectra were then compared against reference spectra adopted from the Human Metabolome Database (HMDB). Spectral similarity was quantified using the cosine similarity score between the measured and reference spectra. All fragment ion identities were additionally validated by exact mass calculation using IUPAC 2016 monoisotopic atomic masses and cross-referenced against known compounds in PubChem and HMDB where applicable. Metabolites were considered positively identified (Metabolomics Standards Initiative level 2), as distinct from formula-level assignments made at ±5 ppm without on tissue MS/MS, which are reported as level 3, when: (i) the precursor [M−H]^−^ matched the theoretical exact mass within ±5 ppm; (ii) at least two diagnostic MS/MS fragment ions were observed with correct mass and expected relative intensity pattern; and (iii) the cosine similarity to the reference spectrum exceeded 0.85.

### Hemolymph collection and LC–MS

Hemolymph was collected as described previously (*49*, *65*). To collect hemolymph, thoraces of approximately 20 flies were pierced with a tungsten needle (Fine Science Tools). Next, the flies were placed in a perforated 0.5 mL Eppendorf tube within a 1.5-ml Eppendorf tube and then centrifuged at 2,348 × *g* at 4 °C. One microliter of the hemolymph was collected and transferred to a new tube. Processed hemolymph was flash frozen on dry ice and kept at 80 °C until metabolomic sample preparation. Hemolymph was resuspended in 80:20 methanol:water for LC-MS analysis. LC was performed on an Atlantis BEH Z-HILIC column (Waters) using a Dionex UltiMate 3000 UHPLC system (Thermo Fisher Scientific). Solvent A consisted of 95:5 water:acetonitrile with 20 mM ammonium acetate and 20 mM ammonium hydroxide at pH 9.4. Solvent B consisted of acetonitrile. The gradient used for metabolite separation was 0 min, 90% B; 2 min, 90% B; 3 min, 75% B; 7 min, 75% B; 8 min, 70% B; 9 min, 70% B; 10 min, 50% B; 12 min, 50% B; 13 min, 25% B; 14 min, 25% B; 16 min, 0% B; 21 min, 0% B; 21 min, 90% B; and 25 min, 90% B. MS was performed on an Orbitrap Exploris 240 mass spectrometer (Thermo Fisher Scientific) operated in negative ion mode over an *m/z* range of 70–1,000. Data were analyzed using El-MAVEN Software (Elucidata; elucidata.io) (*66*). For tracing experiments, the natural isotope abundance was corrected using AccuCor (*67*).

### Intracellular metabolite extraction and LC-MS analysis

Metabolite extraction from *Drosophila* was adapted from a published protocol (*27*, *32*). Twenty thoraces, heads, guts, or abdomens were dissected in ice-cold 0.9% normal saline (Thermo Fisher Scientific, 23-062125) after placing the flies under light CO_2_ anesthesia. Dissected tissues were immediately placed in 1.5 mL tubes on dry ice. Cold 80% (v/v) aqueous methanol (450 µL) was added to the tubes, and intracellular metabolites were extracted by homogenizing tissues with a pellet pestle for 1 minute (cycles of 20 s of grinding and 10 s of rest). Samples were centrifuged at 20,000 × *g* for 10 min in the cold room. Supernatants were transferred to a new 1.5 mL tube, dried by vacuum centrifugation (SpeedVac) and stored at 80 °C until mass-spectrometry analysis. Samples were re-suspended using 22 µL HPLC grade water for mass spectrometry. 5 to 7 μL was injected and analyzed using a hybrid 6500 QTRAP triple quadrupole mass spectrometer (AB/SCIEX) coupled to a Prominence UFLC HPLC system (Shimadzu) via selected reaction monitoring (SRM) of a total of 300 endogenous water soluble metabolites for steady-state analyses of samples. Some metabolites were targeted in both positive and negative ion mode for a total of 311 SRM transitions using positive/negative ion polarity switching. ESI voltage was +4950 V in positive ion mode and 4500 V in negative ion mode. The dwell time was 3 ms per SRM transition and the total cycle time was 1.55 s. Approximately 9 to 12 data points were acquired per detected metabolite. For targeted ^13^C flux analyses, isotopomers from ∼140 polar molecules were targeted with a total of 460 SRM transitions. Samples were delivered to the mass spectrometer via hydrophilic interaction chromatography (HILIC) using a 4.6 mm i.d. × 10 cm XBridge Amide column (Waters) at 400 μL min^−1^. Gradients were run starting from 85% buffer B (HPLC grade acetonitrile) to 42% B from 0 to 5 min; 42% B to 0% B from 5 to 16 min; 0% B was held from 16 to 24 min; 0% B to 85% B from 24 to 25 min; 85% B was held for 7 min to re-equilibrate the column. Buffer A consisted of 20 mM ammonium hydroxide/20 mM ammonium acetate (pH=9.0) in 95:5 water:acetonitrile. Peak areas from the total ion current for each metabolite SRM transition were integrated using MultiQuant v3.2 software (AB/SCIEX).

### Hematoxylin and eosin staining

Meyer’s hematoxylin (Sigma, MHS32) and eosin (Sigma, HT110332) were used for Hematoxylin and eosin (H&E) staining. For MALDI-MSI sections, the matrix was removed with ice-cold methanol for 5 min. Slides were then washed in PBS and water, stained with hematoxylin for 15 min, dehydrated in 95% ethanol for 30 s, and counterstained with eosin for 1 min. Sections were further dehydrated in 95% ethanol followed by absolute ethanol, washed in xylene for 2 min, and mounted with Cytoseal 60 mounting medium (Thermo Fisher Scientific, 8310-4). Whole-slide images were acquired on a Hamamatsu NanoZoomer and analyzed in NDP.view2 software (Hamamatsu). Anatomical regions of interest were manually annotated in NDP.view2 using the freehand region-drawing tool.

### Metabolite and isotopologue processing using IsoScope

Individual-fly MALDI-MSI datasets were converted to MATLAB .mat files containing an msi data structure and processed using IsoScope as previously described (*14*). Metabolites were defined by their molecular formulas using predefined negative-ion metabolite lists. Peak extraction was performed in negative-ion mode using a charge of −1 and a mass tolerance of 5 ppm, with isoType = 1 for ^13^C-tracing datasets and isoType = 2 for ^15^N-tracing datasets. Metabolite intensities were extracted at each spatial measurement location using the *msi_get_idata* function and assembled into matrices containing the X and Y coordinates and the intensity of each metabolite. This was equivalent to using the batch intensity module in IsoScope graphical user interface. For selected metabolites, isotopologue intensities from M+0 through M+n were extracted. Locations with no detected isotopologue signal were retained as missing values. For fractional labeling analyses, pixels with a total isotopologue intensity below 10^5^ were excluded.

### Quality control, normalization, and metabolite selection

Metabolite intensity datasets processed from IsoScope were imported into Python as AnnData objects (*33*). Rows represented individual MSI pixels, and columns represented metabolites. Quality control was performed separately for each fly. For each metabolite, spatial coverage was calculated as the percentage of pixels with nonzero intensity, and mean intensity was calculated across all pixels. Metabolites with a mean intensity of at least 10,000 were classified as excellent, good, or fair when detected in at least 80%, 50%, or 10% of pixels, respectively. Metabolites satisfying the criteria for at least fair classification were retained; others were excluded. Pixels with zero summed metabolite intensity were also removed. After filtering, individual flies retained between 20 and 95 metabolites, with a median of 76; this per-fly quality control precedes the across-fly filter. Total ion count normalization was performed by dividing each metabolite intensity by the summed intensity of the retained metabolites within the corresponding pixel (*8*). Allura Red was retained during filtering steps because it served as an exogenous feeding marker and provided a spatial reference for identifying gut lumen-associated regions.

### Aggregation of individual-fly MSI datasets

Metabolites were included if they passed the coverage and intensity criteria in at least 30% of the investigated flies. Because metabolites were extracted by molecular formula, entries such as citrate/isocitrate, AMP/dGMP, ADP/dGDP, leucine/isoleucine and the hexoses were represented as formula-level summed intensities. Likewise, hydroxyproline and N-acetyl-L-alanine had the same molecular formula and identical spatial intensity profiles; they were counted as one metabolite in the final atlas, while both entries were retained in the matrix used for primary dimensionality reduction and clustering analyses. Considering unresolved metabolites as single entries, 81 features were reported in the final atlas, of which 80 were endogenous metabolites. Allura Red dye, an exogenous feeding marker, was also retained. The 68 individual-fly AnnData objects were concatenated using an outer join, and intensity values for metabolites not included in an individual fly were filled with zeros.

### Dimensionality reduction, batch correction, and unsupervised clustering

We applied a log1p transformation to the normalized metabolite intensities and performed principal component analysis (PCA). Harmony algorithm was applied to the resulting principal components using individual fly identity as the batch variable, with a maximum of 20 Harmony iterations (*68*). Using the Harmony-corrected principal components, we constructed nearest-neighbor graphs with neighborhood sizes of 15, 30, and 50. For each setting, we examined the resulting metabolic structure and mapped the cluster patterns to the original spatial coordinates of individual sections. We selected 15 nearest neighbors because this setting retained smaller, spatially contiguous metabolic domains and limited mixing between neighboring regional patterns, whereas neighborhood sizes of 30 and 50 produced broader spatial mixing. Then, we generated a Uniform Manifold Approximation and Projection (UMAP) representation from the selected neighborhood graph using a minimum distance of 0.3 and a random seed of 42. UMAP was used for visualization and was not used directly for tissue assignment. Next, we performed Leiden clustering on the selected neighborhood graph at resolutions from 0.1 to 0.6 using the *igraph* package. These resolutions identified 26, 30, 31, 34, 37, and 37 clusters, respectively (**Fig. S5**).

We compared the resulting resolutions based on (i) changes in total cluster number across adjacent resolutions at a fixed seed and (ii) visual inspection of spatial cluster patterns across independent sections (**Fig. S6**). Lower resolutions captured the major spatial metabolic domains but merged several regional patterns. With seed 0, resolutions 0.5 and 0.6 both yielded 37 clusters and showed similar spatial organization on visual inspection. To assess reproducibility across random seeds, we also repeated Leiden clustering using five seeds (0–4) at each resolution while holding the neighbor graph fixed. Mean pairwise adjusted Rand indices were 0.80, 0.76, 0.66, 0.68, 0.63, and 0.65 for resolutions 0.1-0.6, respectively, indicating that seed-to-seed agreement declined modestly as resolution increased, and resolution 0.6 yielded 37–40 clusters across seeds. We selected resolution 0.6 because it provided a more subdivision of the dominant thoracic population into five regional subclusters, compared with four at resolution 0.5, while maintaining spatially coherent patterns across representative sections (**Fig. S7 and S8**). Of the 37 clusters obtained at resolution 0.6 with seed 0, 18 clusters each contained ≥1% of all pixels and together accounted for 96.7% of the dataset, whereas the remaining 19 clusters each contained <1% and together accounted for 3.3%. Fly coverage also varied, with 13 clusters represented in only 1–4 flies and together accounting for 0.33% of all pixels. Clusters were treated as initial metabolic groupings rather than distinct anatomical regions, and were subsequently evaluated through iterative tissue annotation, including consolidation or reassignment, as described below.

### Tissue annotation

The aim of this analysis was to annotate clusters identified by unsupervised clustering of spatial metabolite profiles as biologically meaningful tissues (**Fig. S9**). Because tissue sections were analyzed without fixation to minimize alterations in metabolite abundance and spatial distribution, Hematoxylin and eosin (H&E)-stained serial sections could not be precisely co-registered with the MSI sections at the pixel level and were therefore not used as the primary basis for annotation. In addition, because the whole adult *Drosophila* contains tissues with different physical properties, unfixed serial sections may undergo uneven cutting or displacement during sectioning and histological processing, limiting precise co-registration between the H&E-stained and MSI sections. Accordingly, we annotated tissues based on spatial metabolite profiles, cluster locations and patterns, and consistency across sections. The resulting tissue annotations do not imply that cluster boundaries correspond exactly to histological or cellular boundaries. The detailed procedure and results are provided in the Supplementary text (**Fig. S10-S15**, see **Supplementary text**).

### Annotation criteria

Clusters and subclusters were annotated using three primary criteria:

i. their enriched metabolite profiles were compared with reference metabolite profiles from four major anatomical regions characterized in the previous study;
ii. their spatial locations within the fly sections were qualitatively compared with known *Drosophila* anatomy; and
iii. the consistency of their spatial patterns was evaluated across the 68 sections.

For metabolite profile evaluation (criterion i), annotations were based on the overall pattern of enriched metabolites rather than on one or two metabolites treated as tissue-specific markers, because individual metabolites are not necessarily restricted to a single tissue. Cluster profiles were compared with metabolomics data previously obtained from dissected *Drosophila* head, thorax, gut, and abdomen without the gut (**Table S3**) (*27*). These reference profiles were used to assess whether the enriched metabolites of a cluster were consistent with its proposed anatomical location. For gut lumen-associated regions, the spatial distribution of Allura Red dye served as a marker.

For spatial evaluation (criterion ii), cluster assignments were mapped back to their original spatial coordinates. We considered the body compartment in which each cluster occurred, its relative position to the head, thorax, or abdomen, its spatial organization, and its relationship to neighboring clusters. Peripheral, circular, or broadly distributed patterns were compared qualitatively with the expected locations and organization of adult *Drosophila* tissues described in published anatomical and histological resources. These included histological descriptions of adult *Drosophila* sections (*69*), anatomical maps of peripheral tissues and nervous structures (*17*), whole-adult spatial transcriptomic maps (*18*), and adult anatomical figures from the online Atlas of *Drosophila* Development (https://www.sdbonline.org/sites/fly/atlas/00atlas.htm). Because histology precisely co-registered to the MALDI-MSI sections was not available, these resources were used to approximate tissue location rather than establish exact histological boundaries.

For consistency evaluation (criterion iii), we examined the number of independent sections containing each cluster or subcluster. Annotated tissues were not required to be present in all 68 sections because they could vary with sectioning orientation and the anatomical structures captured in each sagittal section. Clusters detected in only a small number of sections or representing a small proportion of the analyzed pixels were analyzed more cautiously, particularly when their spatial or metabolite profile evidence was not distinctive. Such clusters were left unannotated throughout subclustering analysis.

Clusters lacking sufficient support across the metabolic, spatial, and consistency criteria were left unannotated. Clusters with partial support received tentative tissue assignments. When further evaluation was needed, selected clusters and neighboring annotations underwent targeted subclustering, followed by re-evaluation using the same criteria. Final tissue labels were retained after review of the combined evidence, consolidation of working annotations, and standardization of tissue names (**Fig. S9**).

### Differential metabolite abundance analysis

At each clustering and subclustering stage, metabolites were ranked for each cluster using the Wilcoxon test on normalized metabolite intensities with the *rank_genes_groups* tool function in Scanpy. The highest-ranked metabolites for each cluster were reviewed with their scores, log_2_ fold changes, and adjusted values. Metabolite profiles were visualized using dot plots generated with the *rank_genes_groups_dotplot* plotting function in Scanpy. After the final tissue annotations were established, the analysis was repeated using the final annotated tissues, and selected tissue-enriched metabolites were summarized in a final dot plot.

### Spatial mapping and evaluation of cluster distributions

At each clustering and subclustering stage, cluster assignments were mapped to their original spatial coordinates using the *spatial* plotting function in Scanpy. Each fly section was visualized separately, and cluster distributions were compared across all 68 sections. Individual clusters were highlighted as needed to examine their location, shape, and consistency. After the final tissue annotations were established, the annotated tissue distributions were mapped across all sections using a consistent set of tissue labels and colors.

### Spatial neighborhood enrichment analysis

Spatial associations among annotated tissues were evaluated using the neighborhood enrichment analysis in Squidpy (*34*). For each pair of annotated tissues i and j, the observed number of spatial graph connections between pixels assigned to the two tissues was counted and denoted c_ij_. Tissue labels were then randomly permuted 1,000 times while the spatial connectivity graph was kept. The mean, µ_ij_, and standard deviation, σ_ij_, of the permuted connection counts were used to calculate an enrichment z score:

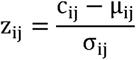

Positive z scores indicate that two tissues were spatial neighbors more frequently than expected by random labeling, whereas negative z scores indicate that they were neighbors less frequently than expected. The resulting z score matrix was visualized as a heatmap, with tissues hierarchically ordered using single linkage.

### Spatial autocorrelation analysis

Spatial autocorrelation of normalized metabolite intensities was evaluated by calculating Moran’s I in Squidpy (*34*). For each metabolite, Moran’s I was calculated as follows:

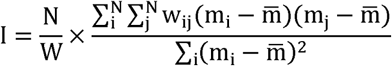

where m_i_ and m_j_ are the normalized intensities at pixels i and j, m is the mean intensity across all pixels, w_ij_ is spatial weights between pixels, N is the total number of pixels, and W=∑_i_∑_j_ w_ij_ is the sum of all spatial weights. Positive Moran’s I values indicate that pixels with similar metabolite intensities tend to occur near one another. Statistical significance was assessed using 100 permutations, so the smallest attainable *P* value is 0.01 before correction, and *P* values were adjusted for multiple testing using the Benjamini–Hochberg method. Metabolites were ranked by their Moran’s I values, with higher values indicating more strongly spatially organized distributions. Pairwise Spearman correlations were calculated using their normalized intensities, and the resulting correlation matrix was hierarchically clustered using average linkage and correlation distance.

### Comparison with dissected region-specific metabolomics

Metabolite profiles from a representative Fly-Iso-Imaging (FISO) sample were compared with region-specific metabolite profiles (RegMet) from dissected head, thorax, gut, and abdomen measured by LC-MS (*27*). For both datasets, z-scores were calculated for each metabolite across clusters or dissected regions. KEGG identifiers were used to identify metabolites detected in both datasets. The top 20 enriched metabolites from each FISO cluster and each region were then selected based on their z-scores, and similarity between the enriched metabolite sets was quantified using the Jaccard similarity index:

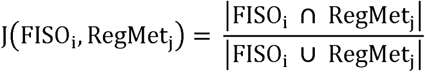

where FISO_i_ and RegMet_j_ represent the KEGG-mapped enriched metabolite sets from FISO cluster i and RegMet region j, respectively.

### Comparison with dissected tissue-specific metabolomics

Tissue-enriched metabolites identified by FISO across 68 flies were compared with dissected tissue metabolomics from FlyMet (*36*). For FISO data, tissue-specific enriched metabolites were ranked based on the scores obtained from differential metabolite abundance analysis. For FlyMet data, metabolite abundances were log_2_-transformed, and the relative abundance of each metabolite in each tissue compared with its median across all dissected tissues was calculated and ranked by magnitude. KEGG identifiers were used to identify metabolites detected in both datasets. The top 19 metabolites from each FISO tissue and each FlyMet tissue were then compared, and similarity between the enriched metabolite sets was quantified using the Jaccard similarity index:

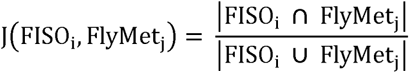

where FISO_i_ and FlyMet_j_ represent the KEGG-mapped enriched metabolite sets from FISO tissue i and FlyMet tissue j, respectively.

For the radial plots, relative Jaccard similarity was calculated for each FISO tissue by normalizing the Jaccard similarity index values to the maximum value across all FlyMet tissues, with the largest FlyMet tissue match set to 1.

### 13C and ^15^N labeling analysis

For each metabolite, isotopologue intensities from M+0 through M+n were extracted using IsoScope. Fractional labeling of metabolite was defined as:

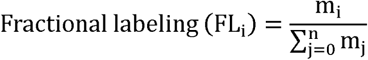

where m_i_ is the M+i isotopologue abundance and 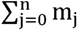 is the summed abundances across all isotopologues from M+0 to M+n. For [U-^13^C_6_]glucose isotope tracing experiments, each isotopologue fractional labeling was compared across the annotated tissues using one-way ANOVA, followed by Tukey’s HSD pairwise comparisons; tissues sampled from the same fly were treated as independent, and *P* values were not adjusted across metabolites and tissues.

The weighted fractional labeling was also defined as:

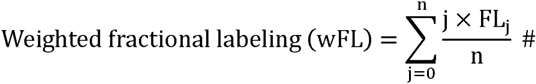

where n is the number of atoms of the tracer element in the metabolite: carbon for ^13^C tracers and nitrogen for ^15^N tracers. In the ^15^N experiments, n=1 for glutamate and N-acetyl-aspartate, and n=3 for glutathione.

Weighted fractional labeling was calculated for each pixel in each nutrient tracing experiments. Tissues containing at least five pixels with detectable wFL values and pixel coverage greater than 5% were kept. For each tissue, mean wFL was first calculated for each fly and averaged across biological replicates

(wFL_tissue_). To visualize relative labeling patterns within each tracing experiment, wFL was normalized to the median across tissues. For each tracer and metabolite, relative weighted fractional labeling was calculated as:

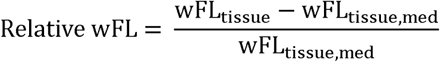

where wFL_tissue_, is the median of the wFL_tissue_ across tissues. wFL was compared across the annotated tissues using one-way ANOVA.

### 13C-metabolic flux analysis

^13^C-metabolic flux analysis (^13^C-MFA) was performed to estimate intracellular metabolic fluxes (*70*). For each pixel, experimentally measured mass isotopomer distributions (MIDs) of target metabolites were obtained from [U-^13^C_6_]glucose tracing experiments in wild-type OreR female flies. The metabolic network was then defined to include the TCA cycle and associated reactions (**Data S2**). Glycolytic and pentose phosphate pathway reactions were simplified given limited measurements of MIDs for intermediate metabolites in those pathways, and glucose carbon stored as trehalose or glycogen was not represented in the network. Isotopic labeling patterns were simulated using the elementary metabolite unit (EMU) framework based on the atom transitions within the defined reaction network (*71*). For metabolites present in subcellular compartments, mixing parameters were used to combine predicted MIDs from the respective pools. Assuming a pseudo-steady state for both metabolic fluxes and isotopic labeling patterns, flux vectors for each pixel were estimated by minimizing the sum of squared residuals between the simulated and experimentally measured MIDs:

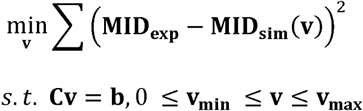

where **Cv**=**b** indicates the steady-state mass balance requirement and other equality constraints. **v_min_** and **v_max_** represent the lower and upper bounds of the flux vectors, respectively. The optimization solver was initialized from 5,000 random starting points per pixel, and the computations were parallelized across a high-performance computing cluster. The resulting solution, **v_opt_**, represents the set of metabolic fluxes in the network model that best fit the experimental MID data for each pixel. The method used to solve this optimization problem has been described previously (*72*). All estimated fluxes are expressed relative to the glucose input flux of the same pixel, so they describe how carbon is partitioned within a tissue rather than absolute rates and are not comparable between tissues on an absolute basis. Pixel-level best-fit fluxes were averaged within each annotated tissue of each individual fly. Tissue-specific mean fluxes were then averaged across flies. For each reaction ratio or fraction, values were compared across tissues using one-way ANOVA, followed by Tukey’s HSD pairwise comparisons. For network visualization, mean flux values were mapped onto a curated metabolic network using the *NetworkX* package in Python.

### Direct contribution analysis

Direct contributions of circulating nutrients to tissue metabolites or other circulating nutrients were estimated from normalized weighted fractional labeling, as previously described (*44*, *47*). Details are provided in the supplementary text. In brief, assuming isotopic pseudo-steady state, the measured labeling of each target metabolite was modeled as:

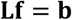

where **L** is an m × n matrix containing the normalized labeling of n circulating nutrients across m tracer experiments, **f** is an n × 1 vector of their direct fractional contributions to the target metabolite, and **b** is an m × 1 vector containing the measured normalized labeling of the target metabolite across the m tracer experiments. Fractional contributions were estimated by solving the following constrained least-squares optimization problem:

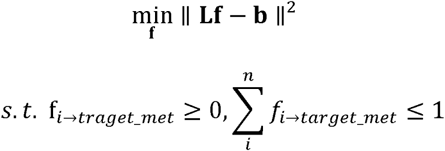

where *f_i_*→*_traget_met_* is the estimated direct fractional contribution of circulating nutrient i to the target metabolite. The nonnegativity constraint was applied to prevent negative contribution estimates; estimates of zero may therefore lie at this bound rather than indicate the absence of a contribution. The sum constraint ensured that the total estimated fractional contribution did not exceed one. Because this constraint is an inequality, 1 *∑_i_f_i_* is the fraction of the target metabolite’s labeling that the modelled nutrient set does not account for; it varies between tissues and should be read alongside the estimated contributions. Uncertainty was estimated by Monte Carlo sampling (5,000 iterations) of the measured means and their SEM; this propagates measurement uncertainty only, not uncertainty in the set of nutrients included in the model.

### Lifespan analysis

For survival analysis, flies were collected within 24 h post-eclosion (20 females and 3 males per vial) and maintained on standard laboratory fly food at 18 °C, with 60% humidity, and a 12 h light/dark cycle, as previously described (*51*). After two days, mature and mated females were transferred to 29 °C to induce gene expression. To maintain vial cleanliness, flies were transferred to fresh vials every two days, and the number of dead flies was recorded daily.

## ACKNOWLEDGEMENTS

We thank all members of the Perrimon laboratory for useful discussions and experimental advice. We also thank Dr. Connie Cepko and Dr. Ryan Delgado for training and access to cryostat. We also thank the Research Computing Group at Harvard Medical School for access to the O2 High Performance Compute Cluster.

## AUTHOR CONTRIBUTIONS

Conceptualization: SJM, SMD, NP Methodology: SJM, KH, JLEB, CN, JMA, SMD

Investigation: SJM, KH, JLEB, CN, YP, YC, HN, VP, MAN, JMA, SMD

Visualization: SJM, KH, JLEB, CN, YP

Funding acquisition: SMD, NP

Project administration: SMD, NP

Supervision: SMD, NP

Writing – original draft: SJM, SMD, NP

Writing – review & editing: SJM, KH, JLEB, CN, YP, YC, HN, VP, MAN, YH, JA, EF, JMA, SMD, NP

