## Supplemental Materials for "Spatial isotope tracing maps compartmentalized nutrient use across the adult fruit fly"

**The PDF file includes:**

Supplementary Text
Figs. S1 to S31
Tables S1 to S18

### Supplementary Text

**Allura Red detection and quantification**
Allura Red dye was included in the tracer diet as a visible indicator of food ingestion. Flies showing red color in the abdomen were selected to reduce variation associated with incomplete feeding. The Allura Red signal was measured by MALDI-MSI to assess food ingestion. The detected ion was assigned as a fragment derived from Allura Red based on the expected *m/z* and ESI-MS/MS fragmentation pattern. The ion was reproducibly detected across sections and used to assess relative signal abundance and spatial coverage.

**Normalization methods during data processing**
We applied total ion count (TIC)-based normalization, a method commonly used in mass spectrometry imaging to reduce variation caused by uneven matrix deposition and differences in overall signal intensity (*73*). Each metabolite intensity was divided by the total intensity of all evaluated metabolites within the corresponding pixel. Because a whole-fly sagittal section contains multiple tissues, this normalization may not fully correct for tissue-dependent differences in ionization. We therefore used the normalized data to evaluate relative distribution of metabolite intensities and to perform clustering, rather than to quantify absolute metabolite concentrations. During the initial single-fly analysis shown in **Fig. S3**, we used median normalization. Because the median- and TIC-based normalized data produced similar major cluster patterns, we selected TIC-based normalization for the final analysis of all 68 flies. The resulting clusters were mapped back to the fly sections and compared with known *Drosophila* anatomy to assess their correspondence with anatomical structures.

**Details of tissue annotation**
After obtaining the initial 37 clusters, we performed tissue annotation and refinement according to the annotation strategy described in the **Methods** and summarized in **Fig. S9**. The analysis consisted of 14 rounds of annotation, subclustering, and re-evaluation, summarized in **Table S4**.

In round 1, we assigned initial tissue annotations based on the criteria described in Methods (**Table S5**). The correspondence between the original Leiden clusters and the initial tissue annotations is shown for representative sections in **Fig. S10**. The initial annotations were then mapped across all fly sections to evaluate their spatial distribution and consistency (**Fig. S11**).

In round 2, we refined annotations of eye- and Malpighian tubule-associated regions. These regions showed overlapping enriched-metabolite profiles, including enrichment of uric acid and kynurenic acid, making them difficult to distinguish based on metabolic profiles alone. We therefore used anatomical region masks to separate eye-associated regions in the head from MT-associated regions in the abdomen. In representative sections (**Fig. S12**), eye/Malpighian tubules (MT)-associated annotations were distributed across both the head and abdomen, and portions of the initially annotated MT and MT2 regions extended outside the expected thoracic and abdominal regions. Using the anatomical masks, we annotated these regions as Eye, Eye2, MT, MT2, or unannotated MT-associated regions (**Table S6**).

In rounds 3 to 9, we performed subclustering analysis on specific target regions (e.g., MT regions in round 3; ovary and fat body in round 4). The strategy is summarized in **Fig. S9**. For instance, in round 3, we performed subclustering analysis on MT, MT_2, and annotated_MT tissue types defined from round 2. We then generated a UMAP representation to qualitatively visualize the separation of these clusters (**Fig. S13**). These clusters were mapped to the spatial coordinates of individual flies (**Fig. S14**). This information served as spatial and consistency evidence. For metabolic evidence, we performed differential metabolite abundance analysis on these subclusters (**Fig. S15**). Using this evidence, we evaluated each subcluster and updated the annotation (**Table S7**). This iterative subclustering and refinement of tissue annotations was repeated to evaluate the major regions of the fly (**Table S8-S13**).

In rounds 10 to 13, we re-evaluated provisional and unresolved annotations. These rounds included thoracic edge and cuticle-associated regions, MT and adjacent gut regions, hemolymph and neighboring regions, and gut lumen and fat body-associated regions. Based on this re-evaluation, we consolidated or reassigned annotations where appropriate (**Tables S14-S17**). In round 14, we performed a final review of all working annotations, including consolidation, reassignment, evaluation of remaining uncertain regions, and standardization of tissue nomenclature to generate the final tissue annotations (**Table S18**).

**Details of direct contribution analysis**

The aim of this analysis is to estimate the contribution of circulating nutrients in the hemolymph (the fly equivalent of blood) to tissue TCA cycle metabolites. The calculation was adapted from a previous study by Hui *et al.* (*47*). In brief, the fractional labeling was defined as:

$$\text{Fractional labeling } (\mathrm{FL_i}) = \frac{\mathrm{m_i}}{\sum_{\mathrm{j=0}}^{\mathrm{n}} \mathrm{m_j}}$$

, where $\mathrm{m_i}$ is the abundance (e.g., peak intensity) of the $\mathrm{M+i}$ isotopologue, and $\sum_{\mathrm{j=0}}^{\mathrm{n}} \mathrm{m_j}$ is the summed abundances across all isotopologues. The weighted fractional labeling was then defined as (*37*):

$$\text{Weighted fractional labeling (wFL)} = \sum_{\mathrm{j=0}}^{\mathrm{n}} \frac{\mathrm{j \times FL_j}}{\mathrm{n}}$$

, where $n$ is the number of carbons in the metabolite.

Next, we define the normalized labeling, $\mathrm{L_{A\to X}}$, from the $^{13}$C-labeled hemolymph nutrient A into tissue metabolite X as:

$$\mathrm{L_{A\to X}} = \frac{\mathrm{wFL_{tissue}^{X}}}{\mathrm{wFL_{hemo}^{A}}}$$

, where $wFL_{tissue}^{X}$ is the weighted fractional labeling of tissue metabolite X in the tracer A experiment, and $wFL_{hemo}^{A}$ is the weighted fractional labeling of nutrient A in the hemolymph from the same tracer experiment. Given that TCA cycle metabolites are at pseudo isotopic steady state, we establish a mass balance equation around a TCA metabolite (e.g., malate):

$$\sum_{i}^{n} J_{i\rightarrow TCA} \times wFL_{i,hemo} = J_{TCA\_out} \times wFL_{TCA}$$

, where $J_{i\rightarrow TCA}$ is the input flux generating TCA metabolite from nutrient i in the hemolymph, $wFL_{i,hemo}$ is the weighted fractional labeling of the nutrient i in the hemolymph, and $J_{TCA\_out}$ is the output flux consuming TCA metabolite.

Dividing the equation by $J_{TCA\_out}$, it gives:

$$\sum_{i}^{n} f_{i\rightarrow TCA} \times wFL_{i,hemo} = wFL_{TCA}$$

, where $f_{i\rightarrow TCA} = \frac{J_{i\rightarrow TCA}}{J_{TCA\_out}}$. $f_{i\rightarrow TCA}$ represents the fractional contribution of nutrient i to the TCA cycle metabolite. By normalizing the equation to $wFL_{hemo}^{input}$, the weighted fractional labeling of the input nutrient in the hemolymph, the equation becomes:

$$\sum_{i}^{n} L_{input\rightarrow i} \times f_{i\rightarrow TCA} = L_{input\rightarrow TCA}$$

For the input nutrient itself, $L_{input\rightarrow input} = 1$. $L_{input\rightarrow i}$ represents the normalized labeling of $^{13}$C-labeled input nutrient to other hemolymph nutrient i. For multiple tracing experiments, the equations can be written in matrix form as:

$$\mathbf{Lf} = \mathbf{b}$$

,where

$$\mathbf{L} = \begin{bmatrix} L_{1\rightarrow 1} & L_{1\rightarrow 2} & \cdots & L_{1\rightarrow n} \\ L_{2\rightarrow 1} & L_{2\rightarrow 2} & \cdots & L_{2\rightarrow n} \\ \vdots & \vdots & \ddots & \vdots \\ L_{m\rightarrow 1} & L_{m\rightarrow 2} & \cdots & L_{m\rightarrow n} \end{bmatrix}$$

$$\mathbf{f} = \begin{bmatrix} F_{1\rightarrow TCA} \\ F_{2\rightarrow TCA} \\ \vdots \\ F_{n\rightarrow TCA} \end{bmatrix}$$

and

$$\mathbf{b} = \begin{bmatrix} L_{1\rightarrow TCA} \\ L_{2\rightarrow TCA} \\ \vdots \\ L_{m\rightarrow TCA} \end{bmatrix}$$

Here, each row represents one tracing experiment, and each column represents one circulating hemolymph nutrient included in the model. The vector **f** contains the estimated fractional contributions of each nutrient to the TCA cycle metabolite. We then set up an optimization problem to solve for **f**, fractional contribution to TCA cycle metabolites:

$$\min_{\mathbf{f}} \| \mathbf{Lf} - \mathbf{b} \|^2$$

subject to

$$f_{i \to TCA} \geq 0$$
$$\sum_{i}^{n} f_{i \to TCA} \leq 1$$

The non-negativity constraints were applied to prevent negative contribution values, and the sum constraint was applied to ensure that the total did not exceed 1.

To evaluate uncertainty in the measured labeling values, Monte Carlo sampling was performed. For each iteration, the hemolymph wFL of each nutrient in each tracer experiment was sampled as:

$$wFL_{i,hemo}^{k} \sim N(\overline{wFL}_{i,}, SEM_{i,hemo}^{2})$$

Negative sampled values were set to zero. The normalized hemolymph labeling matrix, $\mathbf{L^k}$, was then reconstructed by normalizing the sampled hemolymph wFL values to the sampled wFL of the input nutrient:

$$L_{input \to i}^{k} = \frac{wFL_{i,hemo}^{k}}{wFL_{input,hemo}^{k}}$$

Similarly, the normalized labeling of the target metabolite was sampled from its mean and SEM to construct the target vector $\mathbf{b^k}$. The fractional contribution vector was then estimated by solving the original constrained objective function for each iteration. This procedure was repeated for 5,000 iterations.

Direct contributions among circulating hemolymph metabolites were estimated using the same method described above for nutrient contributions to TCA cycle metabolites. For each circulating metabolite, its labeling across the tracing experiments was used as the target vector, and contributions from the other hemolymph nutrients were estimated using the same constrained optimization procedure.

**MALDI-MSI data analysis for *yki* flies**

Quality control, data integration, and unsupervised clustering

Three *yki* and control flies were processed using the same per-fly quality-control and TIC-based normalization procedures described in the **Materials and Methods**. Metabolites passing the coverage and intensity criteria were retained, resulting in 74 metabolites. The individual fly objects were then concatenated using an outer join, and values for metabolites not retained in an individual fly were filled with zeros, yielding an integrated dataset of 30,253 pixels. PCA was performed on the normalized metabolite intensities, and seven principal components explaining at least 90% of the cumulative variance were retained. Harmony was applied to the resulting principal components, with a maximum of 20 Harmony iterations. Using the Harmony-corrected principal components, we generated nearest-neighbor graphs with neighborhood sizes of 15, 30, and 50, and used 50 nearest neighbors for the final integrated graph. For visualization, a UMAP representation was generated from the selected neighborhood graph using a minimum distance of 0.3. Leiden clustering was performed on the selected neighborhood graph at resolutions of 0.2, 0.3, 0.4, and 0.5, which identified 6, 8, 11, and 12 clusters, respectively. Resolution 0.2 was selected for subsequent spatial analysis. Clusters were mapped to the original spatial coordinates of individual sections, and cluster-associated metabolites were ranked using the Wilcoxon test implemented with the *rank_genes_groups* function in Scanpy.

**Tissue annotation**

Initial tissue annotation

The overall tissue annotation strategy followed the method described above. In brief, the six clusters were initially assigned to tissue classes based on the three criteria, resulting in annotations corresponding to Eye, CNS, Muscle, Muscle_2, Fat body/oenocyte, and Hemolymph. These assignments were mapped to the original spatial coordinates of each control and *yki* section and inspected across individual flies. Gut-associated regions were initially identified using Allura Red, an exogenous feeding marker of gut contents. For each pixel, the maximum Allura Red intensity was calculated, and pixels with values at or above the 60$^{th}$ percentile of the maximum intensities across the integrated dataset were assigned as Gut.

<u>Tissue annotation refinement</u>
To further refine the tissue assignments, previously generated body-region annotations for the same control and *yki* flies were transferred to the integrated dataset. These annotations contained manually defined regions corresponding to head, thorax, abdomen, gut, and reproductive system based on the post-H&E staining analysis. Pixels corresponding to the reproductive system were assigned as reproductive_system. For gut annotation, pixels identified either by Allura Red intensities or by the manually defined gut region were assigned as Gut, while the remaining metabolite-based tissue annotations were retained. The refined tissue annotations were evaluated by mapping them to a UMAP and to the original spatial coordinates of individual sections. Metabolite enrichment across the annotated tissues was subsequently evaluated using the Wilcoxon test implemented with the *rank_genes_groups* function in Scanpy.

<u>Tissue annotation refinement - Fat body and hemolymph</u>
Pixels initially annotated as Fat body/oenocyte or Hemolymph were further evaluated by subclustering. Using the Harmony-corrected principal components from the integrated analysis, we constructed a nearest-neighbor graph with 15 neighbors and generated a UMAP representation. Leiden clustering was performed at a resolution of 0.2, resulting in six subclusters. The resulting subclusters were mapped to the original spatial coordinates of individual sections and evaluated across flies based on their spatial distributions, their relationships to the initial tissue annotations, and their enriched metabolite profiles. Metabolites enriched in each subcluster were ranked using the Wilcoxon test implemented with the *rank_genes_groups* function in Scanpy using TIC-based normalized metabolite intensities. Based on the spatial and metabolic evaluation, subclusters 0, 1, and 3 were assigned as Hemolymph, subcluster 2 was assigned as a second fat-body region, and subclusters 4 and 5 were assigned as the primary fat-body region. These refined assignments were mapped back to the full integrated dataset, while tissue annotations outside the subclustered fat body or hemolymph pixels were retained. The two fat-body annotations were renamed as Fat_body and Fat_body_2.

**Differential metabolite abundance analysis**
To evaluate metabolite abundance changes between control and *yki* flies, TIC-based normalized metabolite intensities were averaged across all pixels within each fly to obtain one whole-section pseudobulk value for each metabolite. Mean abundances were compared using Welch's two-sample *t*-test, and *P* values were adjusted across metabolites using the Benjamini–Hochberg procedure. Metabolites with an adjusted *P* value below 0.05 and an absolute $\log_2$ fold change of at least 1 were considered significantly altered. For tissue-specific analysis, TIC-based normalized metabolite intensities were similarly averaged across pixels within each tissue for each fly and metabolite. Control and *yki* flies were compared using Welch's two-sample *t*-test, followed by Benjamini–Hochberg adjustment across metabolites separately within each tissue. Metabolites with an FDR below 0.05 in at least one tissue were selected for visualization. For visualization, metabolite abundances were standardized using z-scores and metabolites were hierarchically clustered.

**$^{13}$C labeling analysis**
As described above, weighted fractional labeling (wFL) was calculated across sample. For each metabolite, mean wFL was calculated for each body region in each fly. Mean wFL values were compared among body regions using one-way ANOVA, followed by Tukey's HSD for pairwise comparisons among body regions for each metabolite.

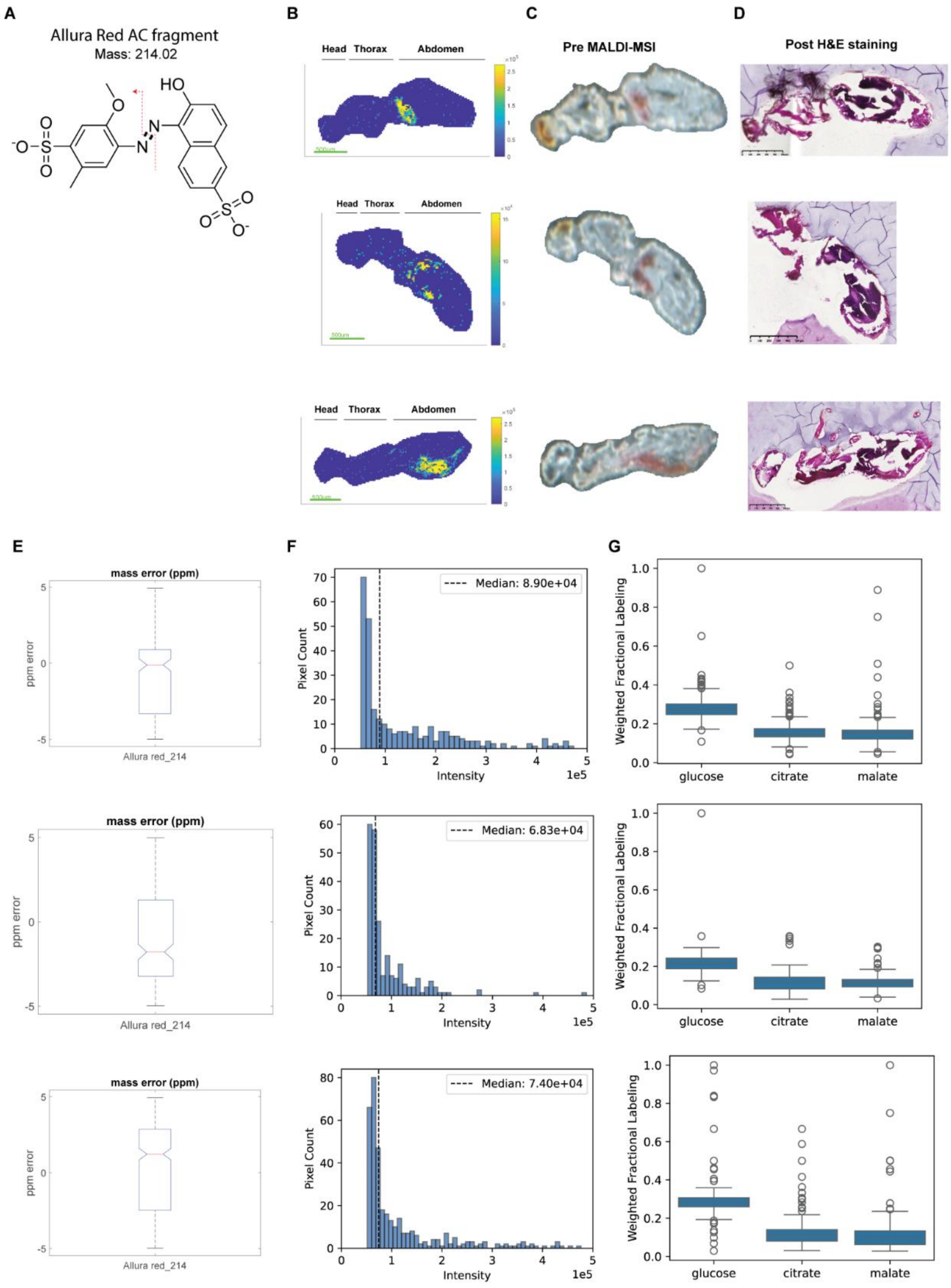

**Fig. S1. Validation of Fly-Iso-Imaging workflow.**

**(A)** Chemical structure of the Allura Red dye (Sigma, 458848) fragment. **(B)** Spatial distribution of the Allura Red dye fragment in sagittal sections from flies fed [U-$^{13}C_6$]glucose after MALDI-MSI. **(C)** Scanned images of the corresponding sections before MALDI-MSI. **(D)** H&E staining of the corresponding sections after MALDI-MSI. **(E)** Mass error of the Allura Red AC fragment across the corresponding sections. **(F)** Pixel intensity distributions of the Allura Red AC fragment. Dashed lines indicate median intensity. **(G)** Weighted fractional labeling of glucose, citrate, and malate in [U-$^{13}C_6$]glucose-fed flies.

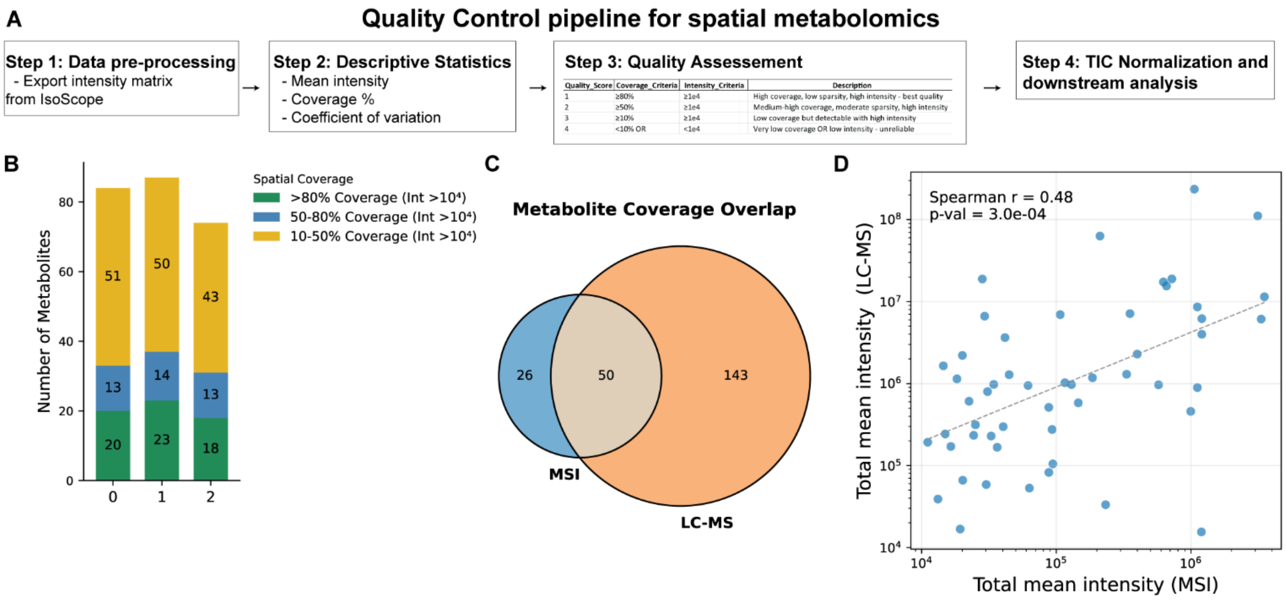

**Fig. S2. Quality control and comparison between MALDI-MSI and LC–MS/MS metabolomics.**

**(A)** Quality-control pipeline for spatial metabolomics, including data preprocessing, descriptive statistics, quality assessment, TIC-based normalization, and downstream analysis. **(B)** Number of metabolites passing spatial coverage and intensity thresholds in each sectioned sample. **(C)** Overlap of KEGG-mapped metabolites detected by MALDI-MSI and LC–MS/MS. **(D)** Correlation between mean metabolite intensities measured by MALDI-MSI and LC–MS/MS. Each point represents one shared metabolite; the dashed line indicates the linear fit.

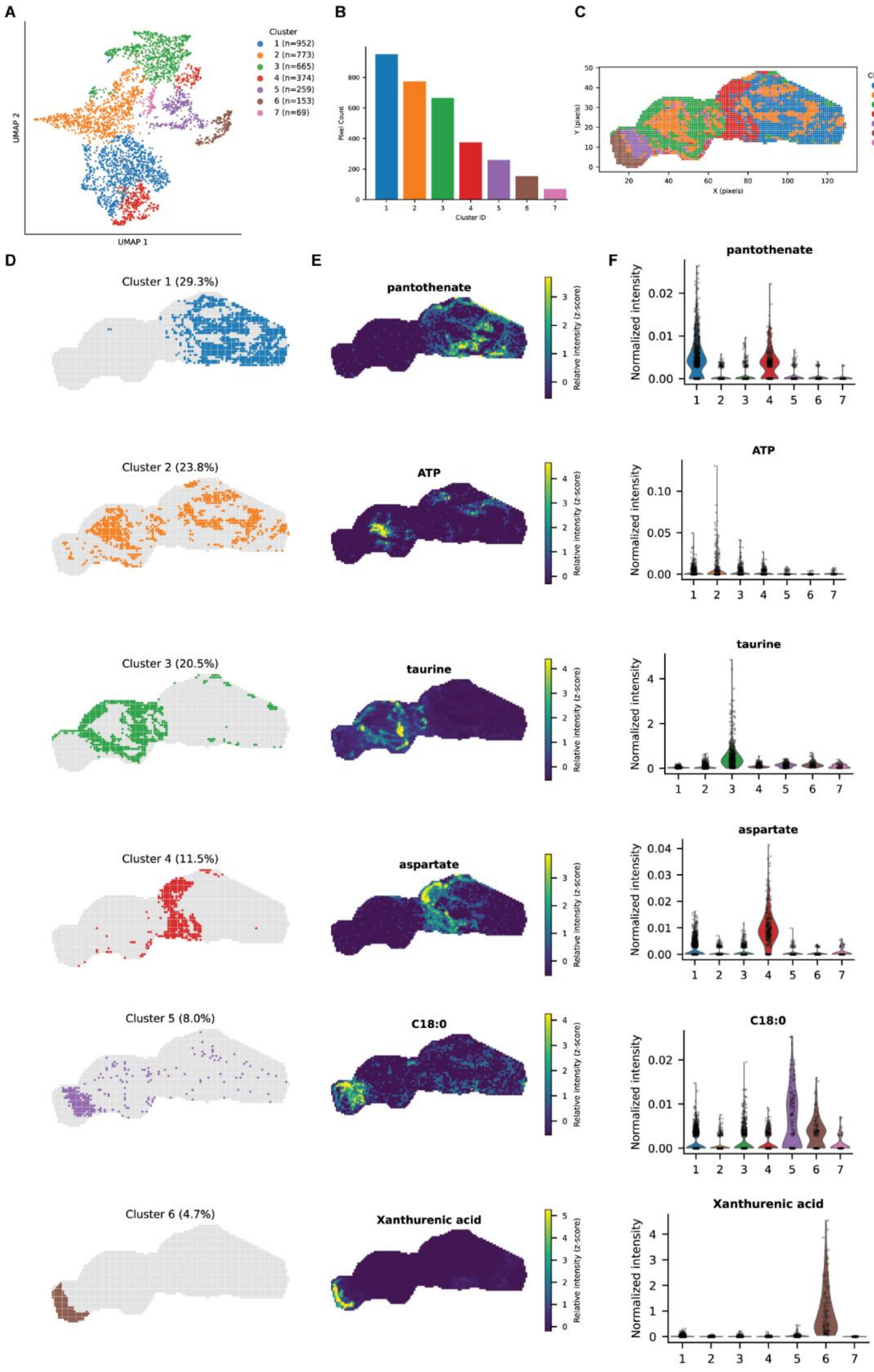

**Fig. S3. Spatial clustering of metabolites in a representative sagittal section.**

**(A)** UMAP of pixel-level metabolite profiles colored by cluster. **(B)** Number of pixels in each cluster. **(C)** Spatial distribution of clusters in the sagittal section. **(D)** Spatial maps of individual clusters. Percentages indicate the fraction of total pixels. **(E)** Representative spatial maps of metabolites enriched in corresponding clusters. TIC-based normalized intensities were scaled per metabolite. **(F)** Quantification of TIC-based normalized metabolite intensities across clusters. Each point represents one pixel.

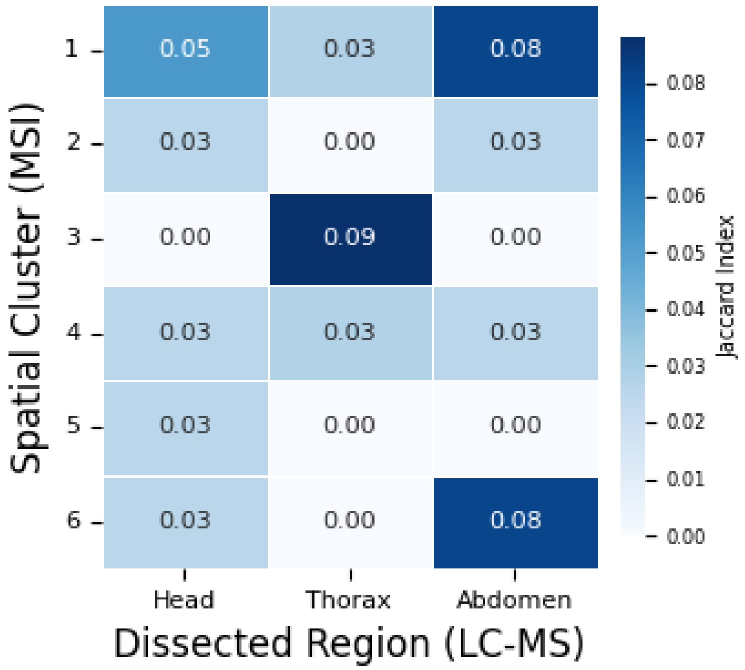

**Fig. S4. Comparison of enriched metabolites between FISO spatial clusters and dissected regions.**

Jaccard similarity between the top 20 enriched metabolites from each spatial cluster in a representative Fly-Iso-Imaging (FISO) sample and region-specific metabolite profiles (RegMet) from dissected head, thorax, and abdomen measured by LC-MS.

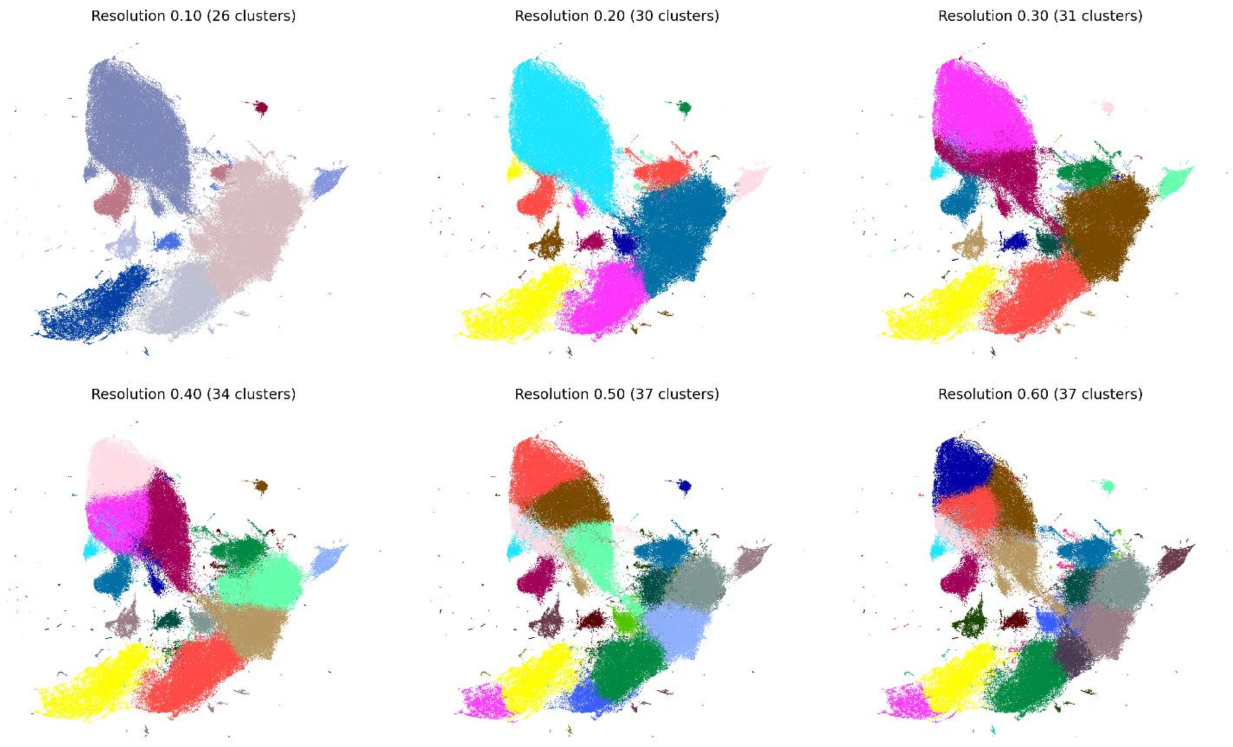

**Fig. S5. Comparison of Leiden clustering resolutions.**

UMAP representations of 187,709 MSI pixels colored by Leiden cluster assignments at resolutions from 0.1 to 0.6. The respective resolutions identified 26, 30, 31, 34, 37, and 37 clusters. The same UMAP embedding, generated from the Harmony-corrected principal-component representation, was used across all panels; only the Leiden cluster assignments differed. Colors denote cluster identities within each resolution and are not intended to correspond across resolutions.

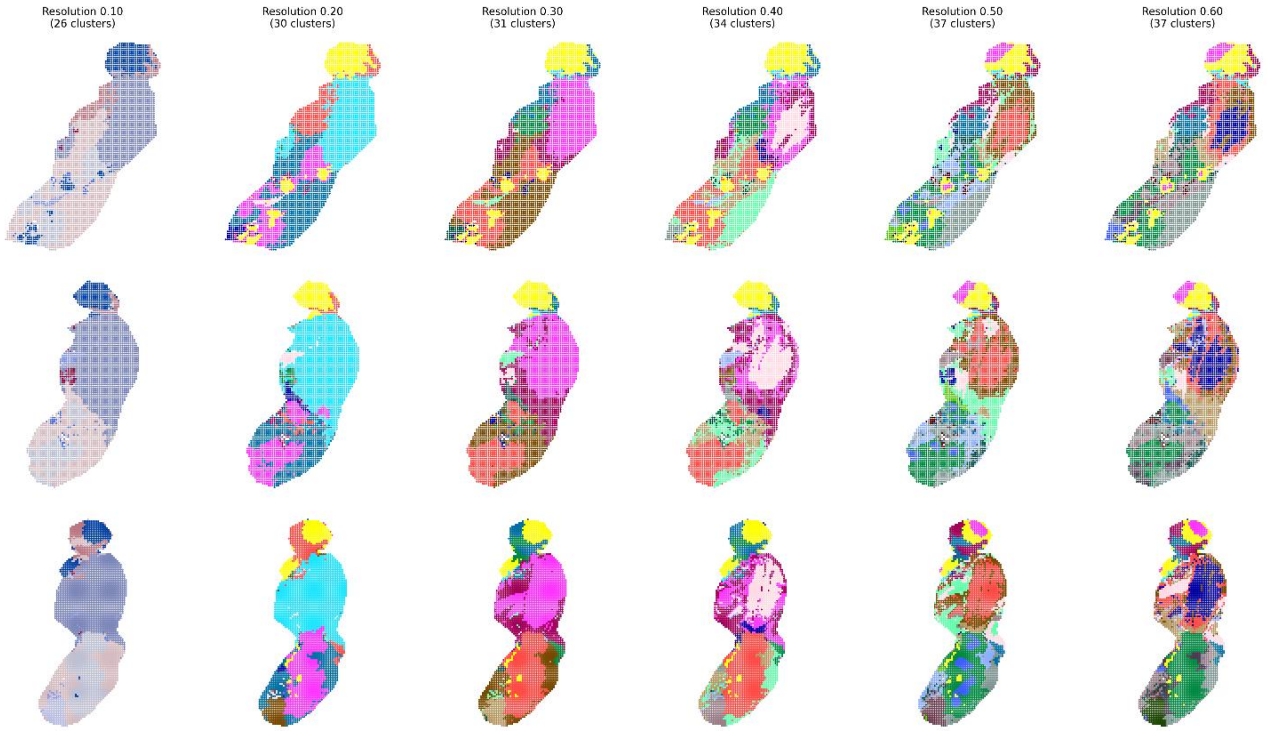

**Fig. S6. Spatial evaluation of Leiden clustering resolutions.**

UMAP representations of 187,709 MSI pixels colored by Leiden cluster assignments at resolutions from 0.1 to 0.6. The respective resolutions identified 26, 30, 31, 34, 37, and 37 clusters. The same UMAP embedding, generated from the Harmony-corrected principal-component representation, was used across all panels; only the Leiden cluster assignments differed. Rows represent different fly samples. Colors denote cluster identities within each resolution and are not intended to correspond across resolutions.

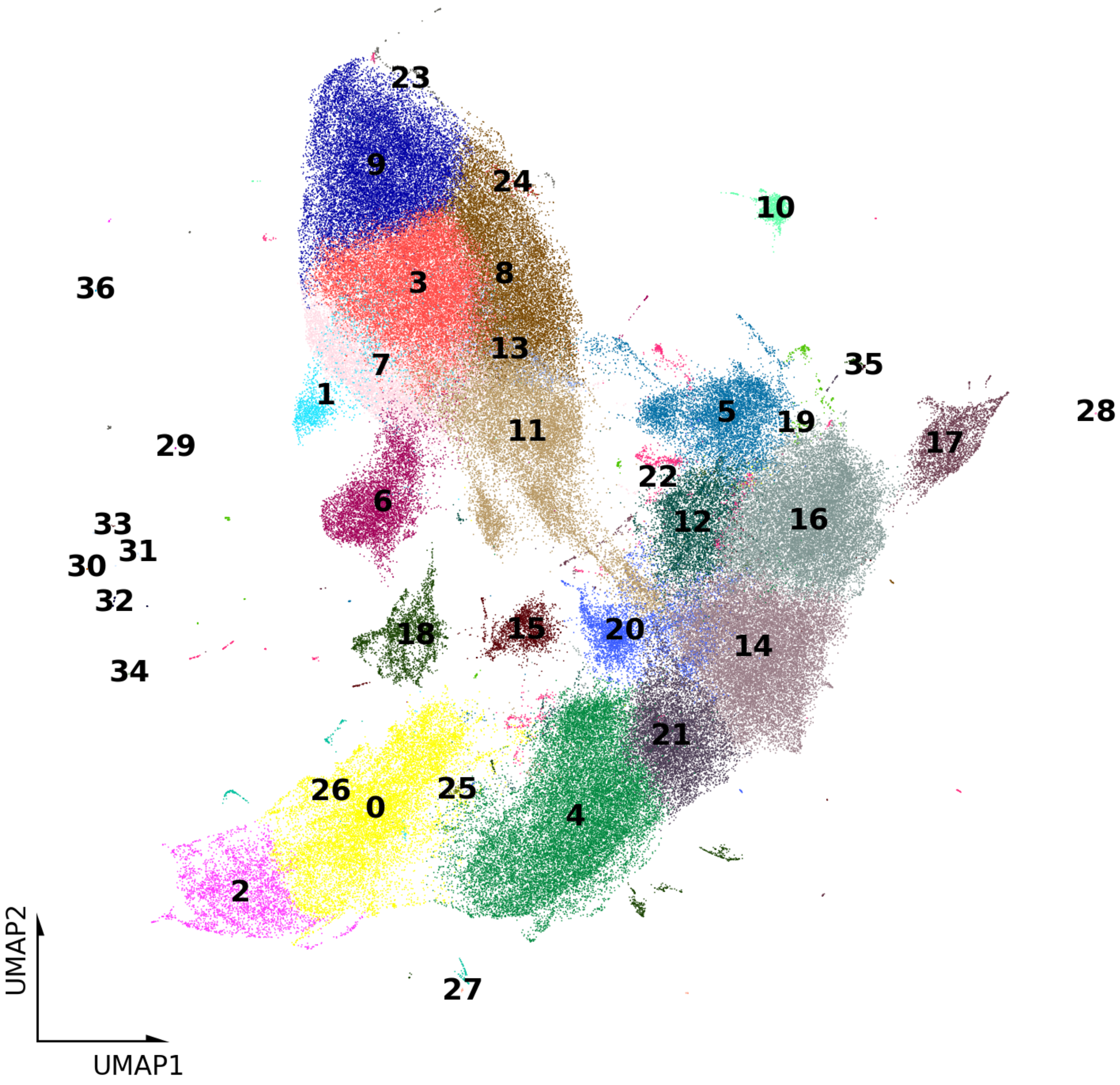

**Fig. S7. UMAP representation of the selected Leiden clustering solution.**

UMAP representation of 187,709 MSI pixels using the Harmony-corrected principal-component representation and the selected nearest-neighbor graph. Pixels are colored by their Leiden cluster assignments at resolution 0.6, which identified 37 clusters. Cluster numbers are shown near the corresponding populations. The UMAP was used to visualize relationships among the clusters and was not used directly to assign tissues. The same UMAP embedding is shown as the resolution-0.6 panel in Fig. S5.

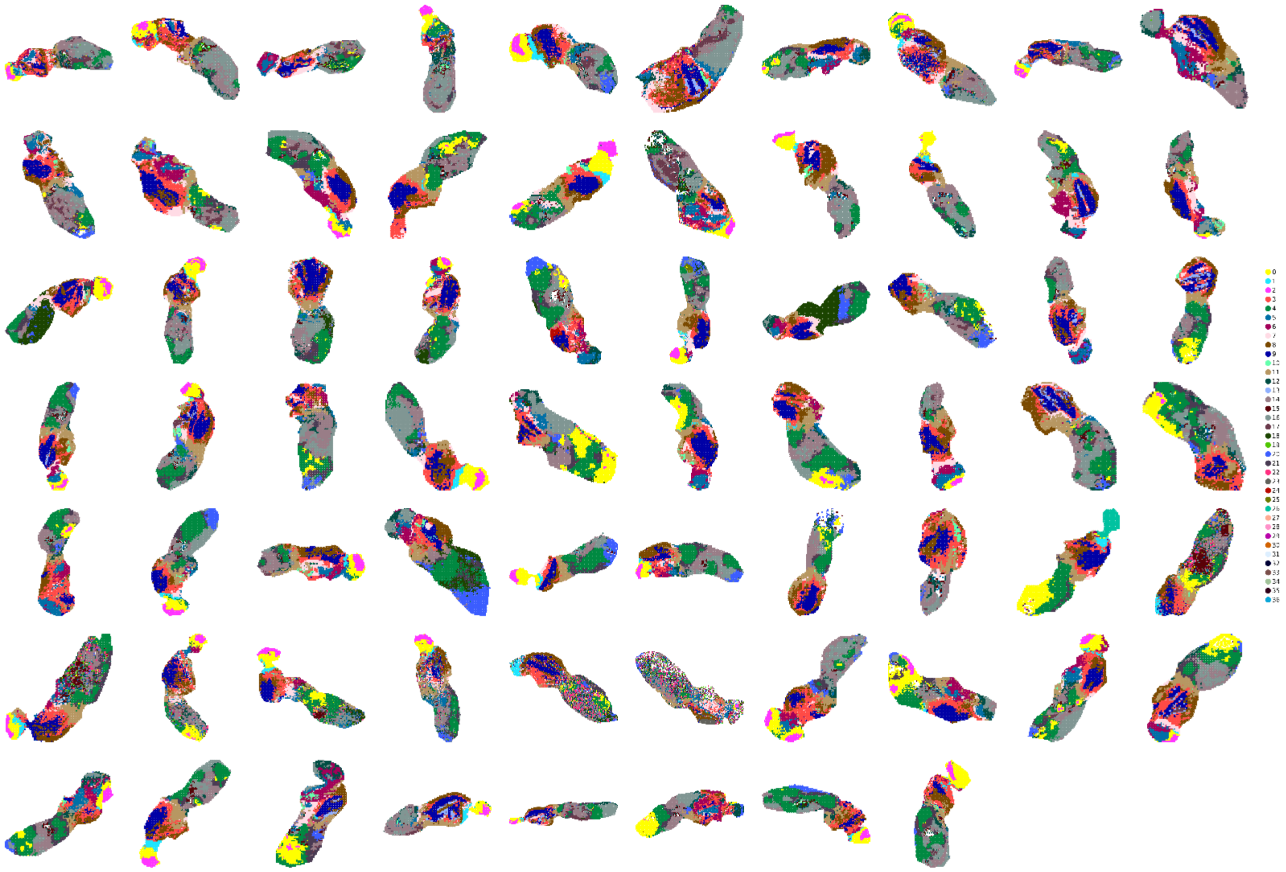

**Fig. S8. Spatial mapping of the selected Leiden clusters across all 68 fly sections.**

Leiden cluster assignments were mapped to the original spatial coordinates of all 68 fly sections using the selected Leiden resolution of 0.6. Colors denote the 37 Leiden cluster identities. These maps were used to assess the anatomical location, shape, continuity, and consistency of each cluster across sections during the initial round of annotation.

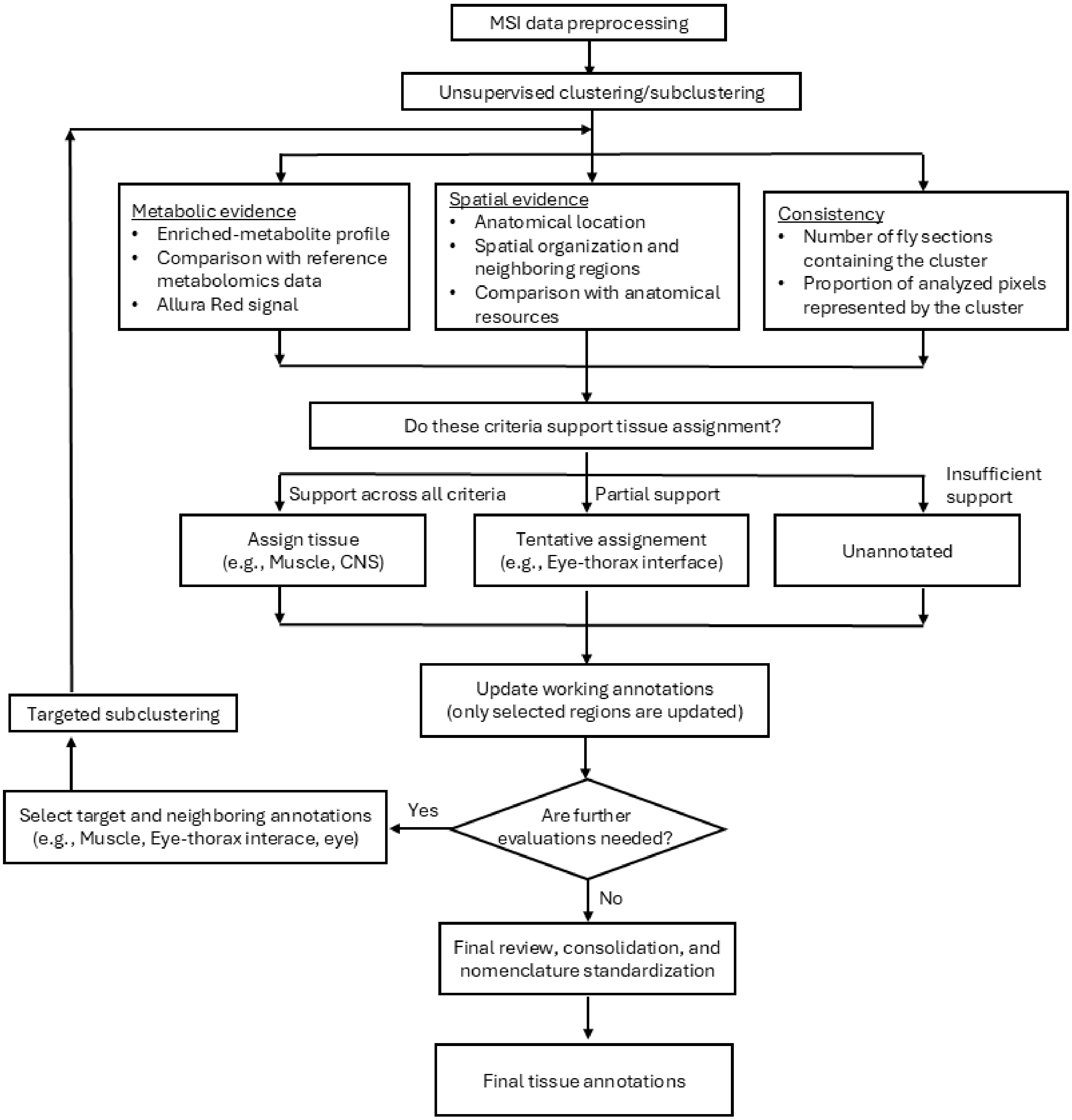

**Fig. S9. Overview of the iterative tissue annotation workflow.**

Clusters and subclusters were evaluated and refined across 14 sequential rounds. In each round, only the selected annotations were updated, while all other annotations were carried forward to the next round. The workflow concluded with final review, consolidation, and nomenclature standardization.

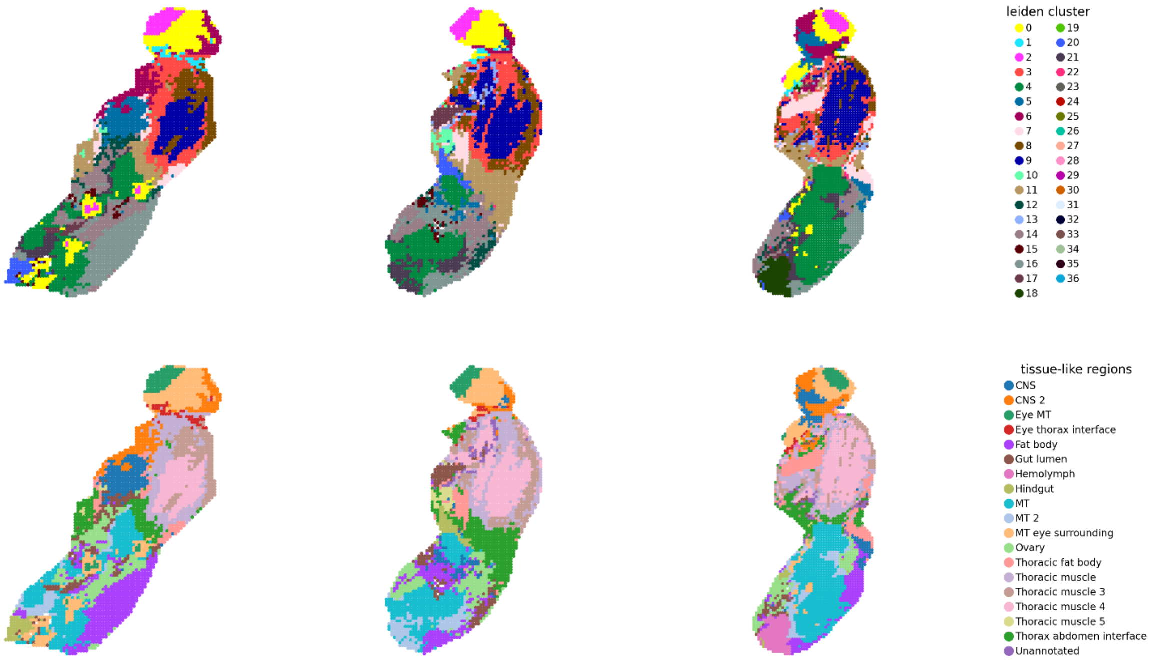

**Fig. S10. Comparison of Leiden clusters and initial tissue annotations in representative fly sections.**

Spatial maps of three representative fly sections showing the 37 Leiden clusters identified at resolution 0.6 (top) and the corresponding initial annotations (bottom).

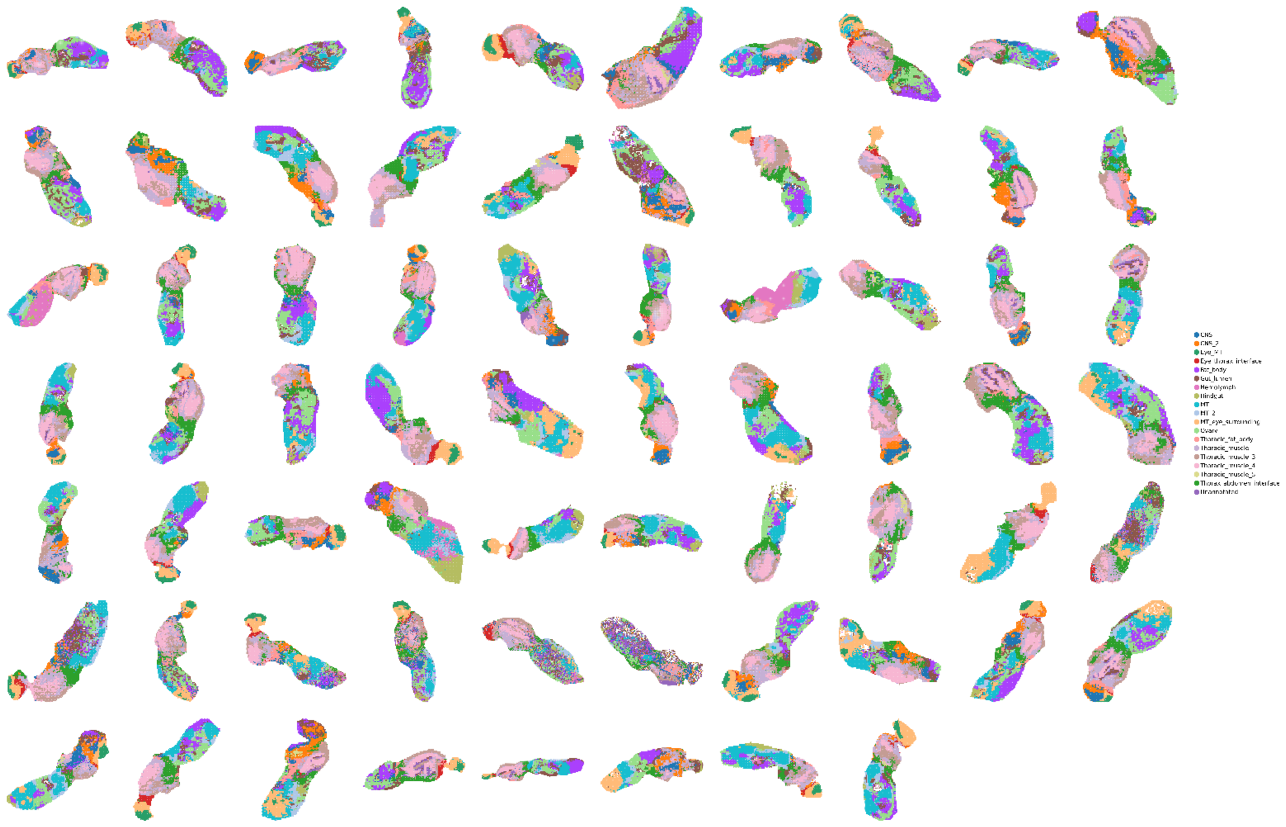

**Fig. S11. Spatial mapping of the initial tissue annotations across all 68 fly sections.**

Initial tissue annotations were mapped to the original spatial coordinates of all 68 fly sections to evaluate their spatial distribution and consistency across independent sections. Colors denote the tissue annotations assigned during the initial round of annotation.

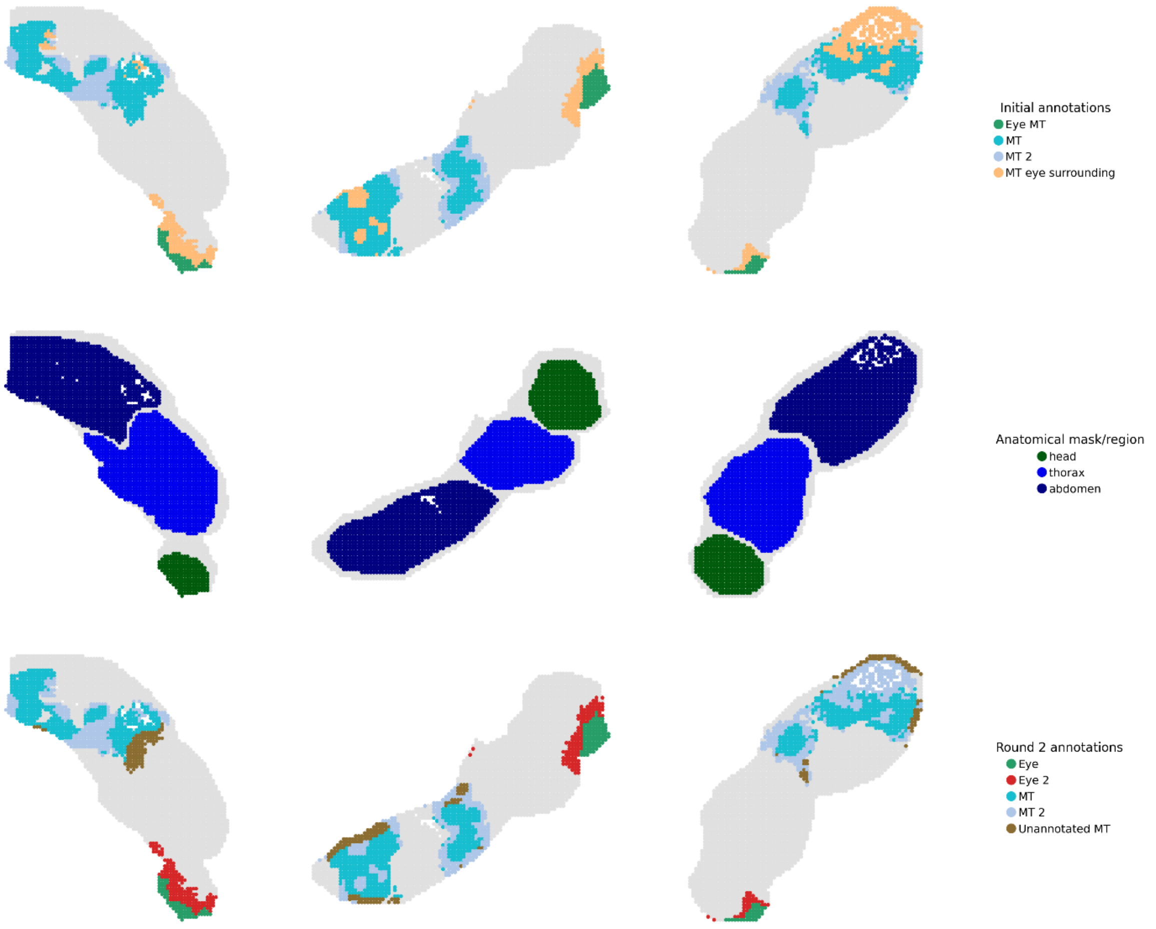

**Fig. S12. Refinement of eye- and Malpighian tubule-associated annotations using anatomical region masks.**

Spatial maps of three representative fly sections showing the initial eye- and Malpighian tubule-associated annotations (top), the corresponding head, thorax, and abdomen masks used for anatomical separation (middle), and the resulting Round 2 annotations separating eye and Malpighian tubule regions (bottom). Candidate regions within the abdomen mask were assigned to Malpighian tubule-associated regions. Remaining candidate regions were assigned to eye-associated regions when located in the head or closer to the center of the head than to the center of the abdomen. Regions not meeting these criteria and outside the anatomical masks were retained as unannotated MT-associated regions. Gray indicates all other regions.

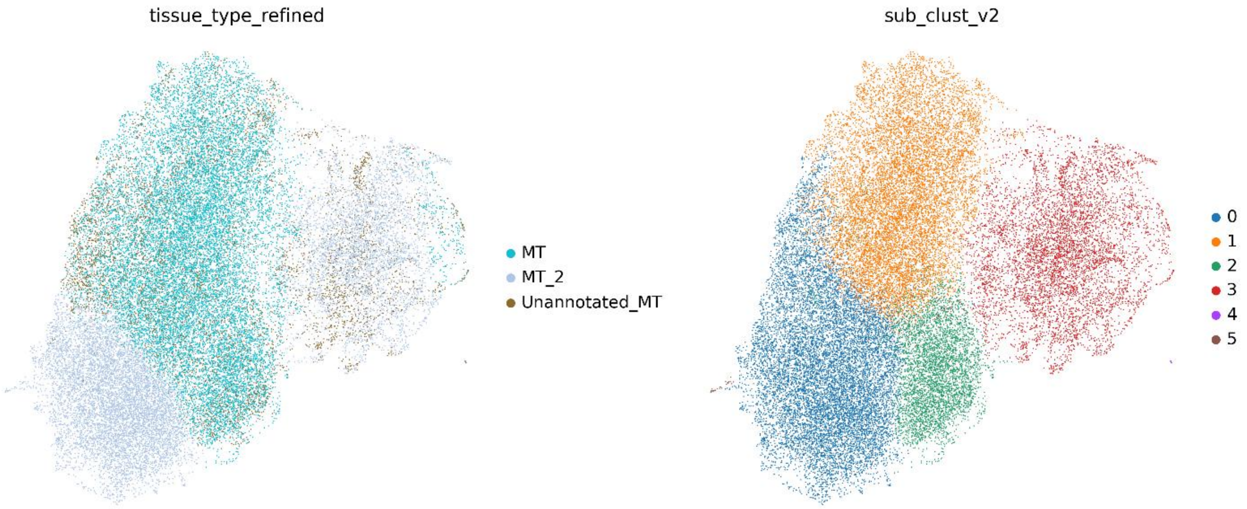

**Fig. S13. UMAP representation of subclustering of MT-associated region annotations.**

UMAP representation of 35,356 MSI pixels previously annotated as MT, MT_2, or Unannotated_MT in tissue_type_refined. A 15-nearest-neighbor graph was constructed using the Harmony-corrected principal-component representation, and UMAP was recalculated for the selected pixels. Pixels are colored by their Round 2 annotations in tissue_type_refined (left) and by their Leiden subcluster assignments in sub_clust_v2 (right). Leiden clustering was performed at resolution 0.2 using the igraph implementation and identified six computational subclusters. The same UMAP embedding is shown in both panels.

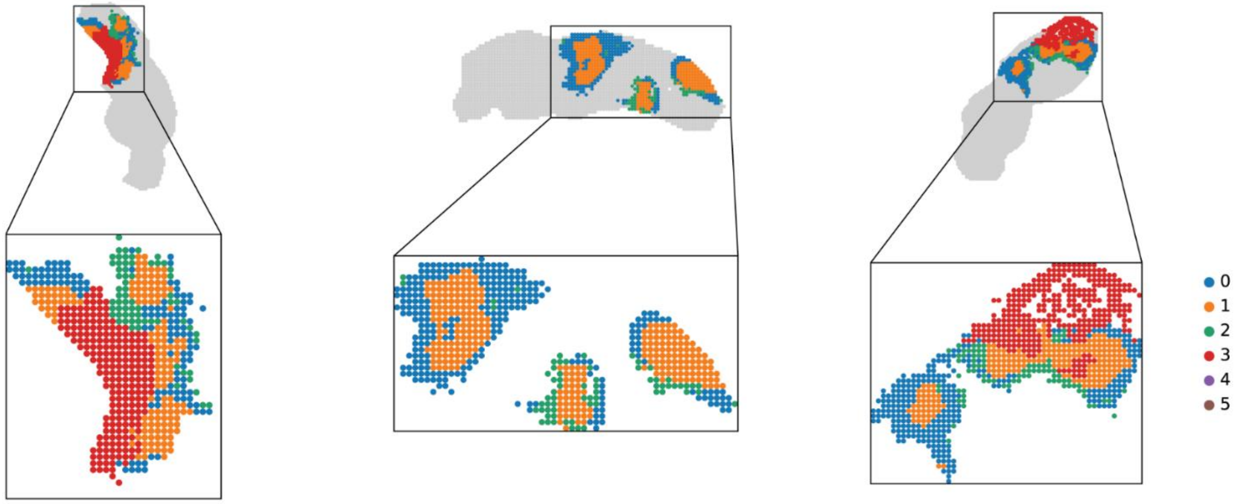

**Fig. S14. Spatial evaluation of subclusters from the MT, MT_2, and Unannotated_MT annotations.**

Spatial maps of three representative fly sections showing subclusters identified among pixels annotated as MT, MT_2, or Unannotated_MT after Round 2. Colors denote the sub_clust_v2 identities. Enlarged maps show only pixels included in the Round 3 subclustering. Insets show the corresponding whole sections, with other pixels in grey and boxes indicating the enlarged regions. The mapping of these subclusters to the updated tissue region annotations is provided in Table S4.

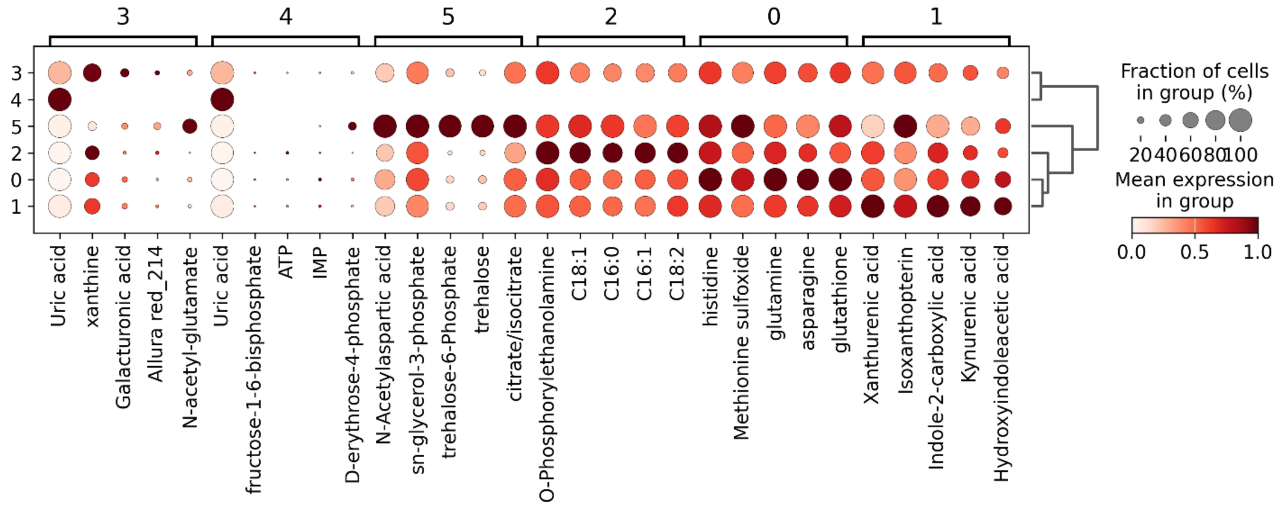

**Fig. S15. Enriched metabolite profiles of subclusters from the MT, MT_2, and Unannotated_MT annotations.**

Dot plot showing the five top-ranked enriched metabolites for each of the six subclusters identified among pixels annotated as MT, MT_2, or Unannotated_MT after Round 2. Each subcluster was compared with all other subclusters using a Wilcoxon rank-sum test. Columns are grouped according to the subcluster for which each metabolite was selected, and rows represent the six sub_clust_v2 identities. Dot size indicates the percentage of pixels with detected signal, and color indicates the mean metabolite abundance scaled from 0 to 1 across subclusters for each metabolite.

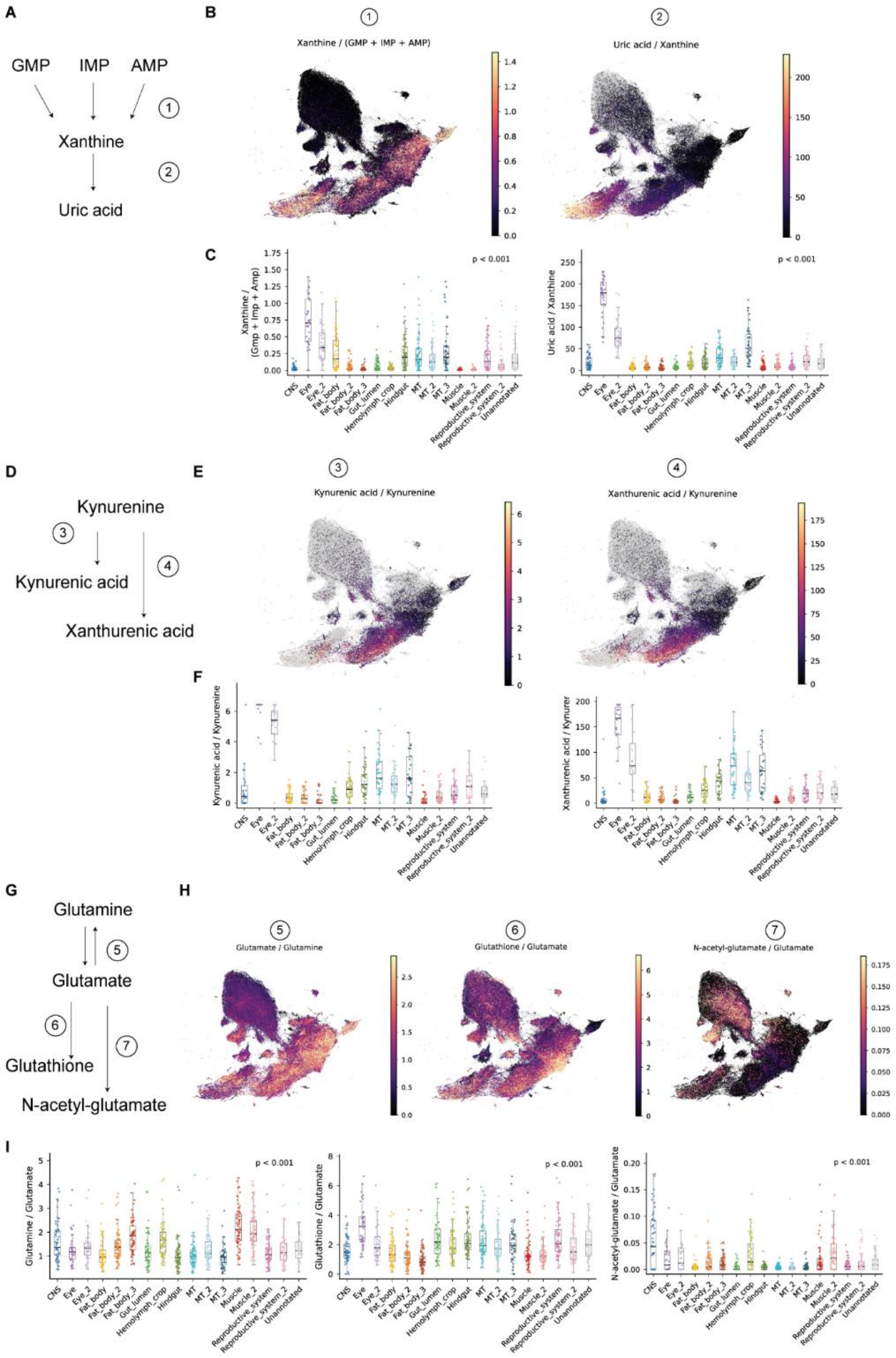

**Fig. S16. Metabolite ratios across annotated tissue regions.**

**(A)** Schematic of purine degradation from guanosine monophosphate (GMP), inosine monophosphate (IMP), and adenosine monophosphate (AMP) to xanthine and uric acid. **(B)** UMAP projections of xanthine/(GMP + IMP + AMP) and uric acid/xanthine ratios. **(C)** Quantification of purine-degradation ratios across annotated tissue regions. **(D)** Schematic of kynurenine conversions to kynurenic acid and xanthurenic acid. **(E)** UMAP projections of kynurenic acid/kynurenine and xanthurenic acid/kynurenine ratios. **(F)** Quantification of kynurenine-associated ratios across annotated tissue regions. **(G)** Schematic of glutamine, glutamate, glutathione, and N-acetyl-glutamate reactions. **(H)** UMAP projections of glutamine/glutamate, glutathione/glutamate, and N-acetyl-glutamate/glutamate ratios. **(I)** Quantification of glutamate-associated ratios across annotated tissue regions.

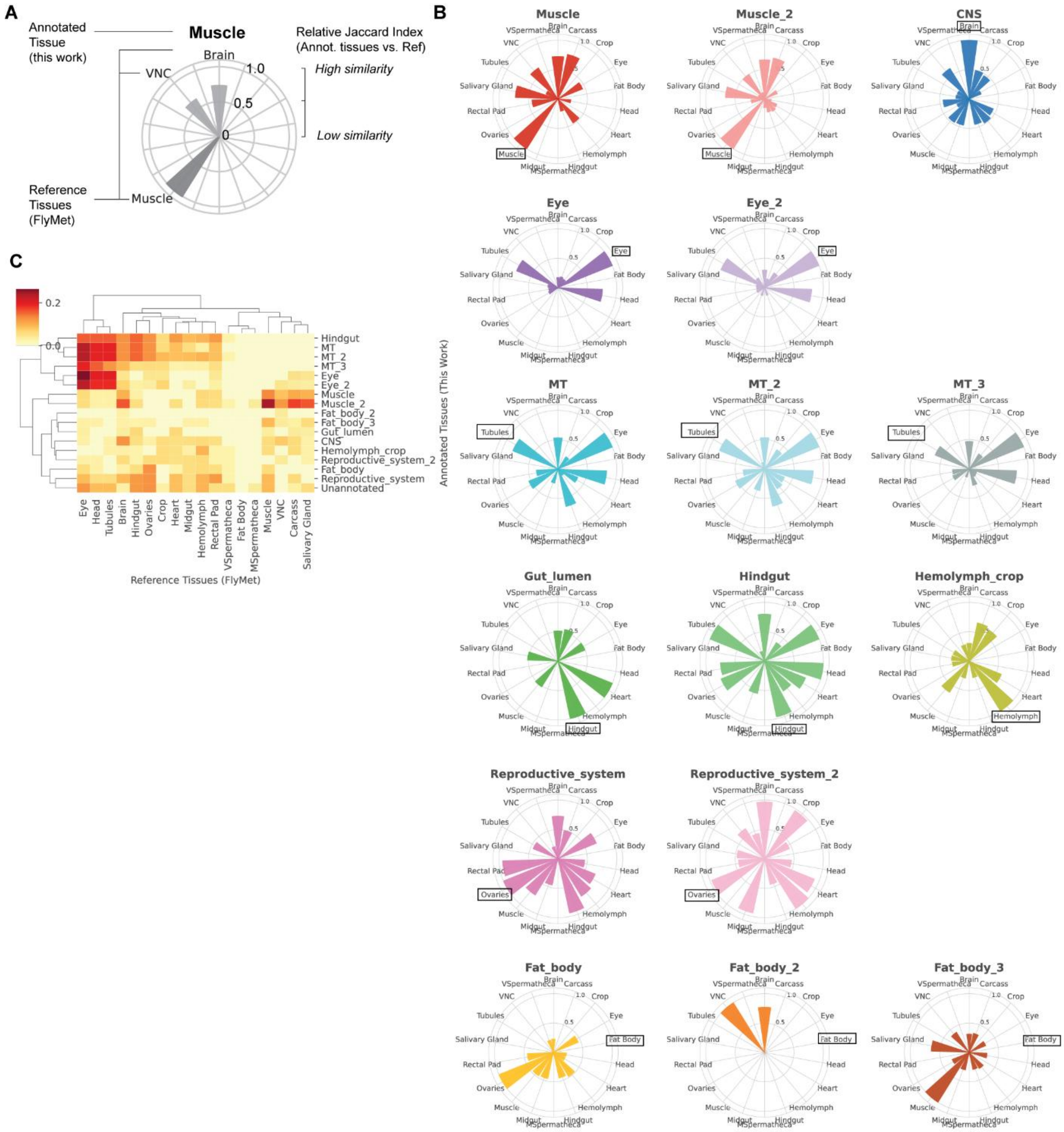

**Fig. S17. Comparison of tissue-enriched metabolites from FISO with dissected tissue-enriched metabolites from FlyMet.**

**(A)** Schematic of the similarity analysis comparing tissue-enriched metabolite sets from each FISO tissue annotation with all FlyMet reference tissues. Each radial axis represents a FlyMet tissue, and bar length indicates the relative Jaccard similarity, with higher values indicating greater metabolite-set similarity. The FISO muscle annotation is shown as an example, with the highest similarity to dissected muscle. **(B)** Radial plots showing the relative Jaccard similarity between each FISO tissue and all FlyMet reference tissues. Values were normalized within each FISO tissue to the maximum Jaccard similarity across FlyMet tissues. Boxed labels indicate the expected corresponding dissected tissue. **(C)** Hierarchically clustered heatmap showing the raw Jaccard similarity index between FISO tissue annotations and FlyMet reference tissues.

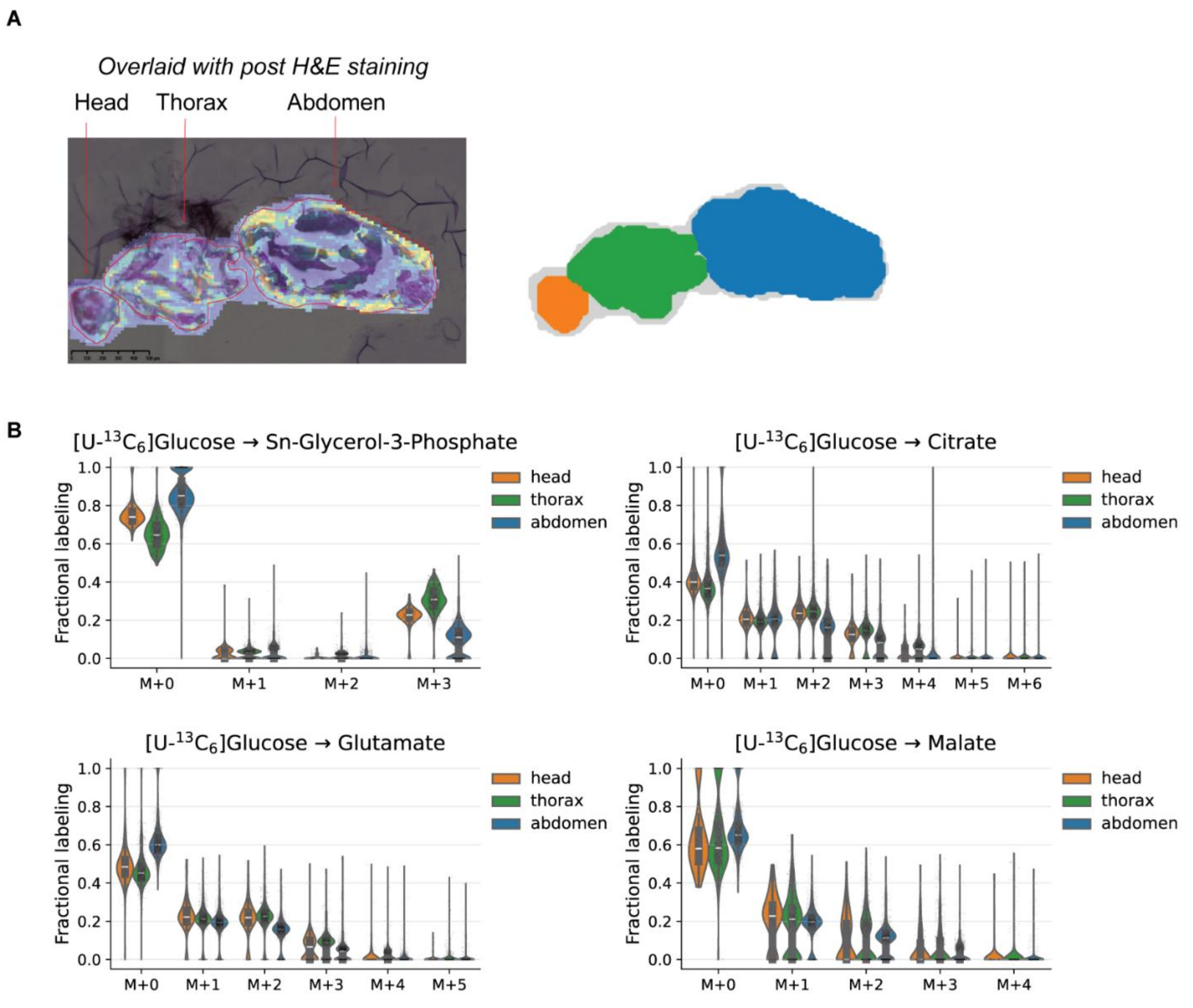

**Fig. S18. Labeling patterns from [U-$^{13}C_6$]glucose tracing across different anatomical regions.**

**(A)** Representative sagittal section overlaid with post-MSI H&E staining, showing head, thorax, and abdomen regions. The right panels are the masks used for respective regions in a representative fly. **(B)** Mass isotopomer distributions of G3P, citrate, glutamate, and malate across head, thorax, and abdomen regions in [U-$^{13}C_6$]glucose-fed flies.

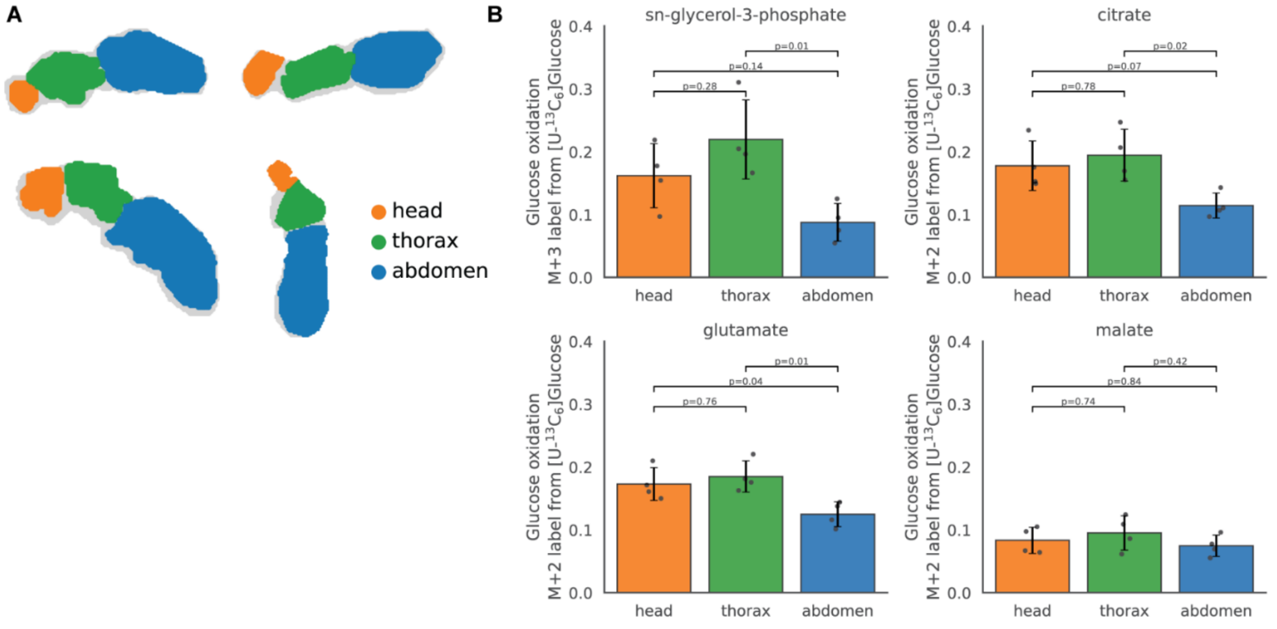

**Fig. S19. Labeling patterns from [U-$^{13}C_6$]glucose tracing across fly samples.**

**(A)** Region masks used to quantify head, thorax, and abdomen labeling across sections. **(B)** Quantification of selected glucose-derived isotopologue labeling for G3P, citrate, glutamate, and malate across anatomical regions. Bars indicate mean ± SEM; points indicate biological replicates.

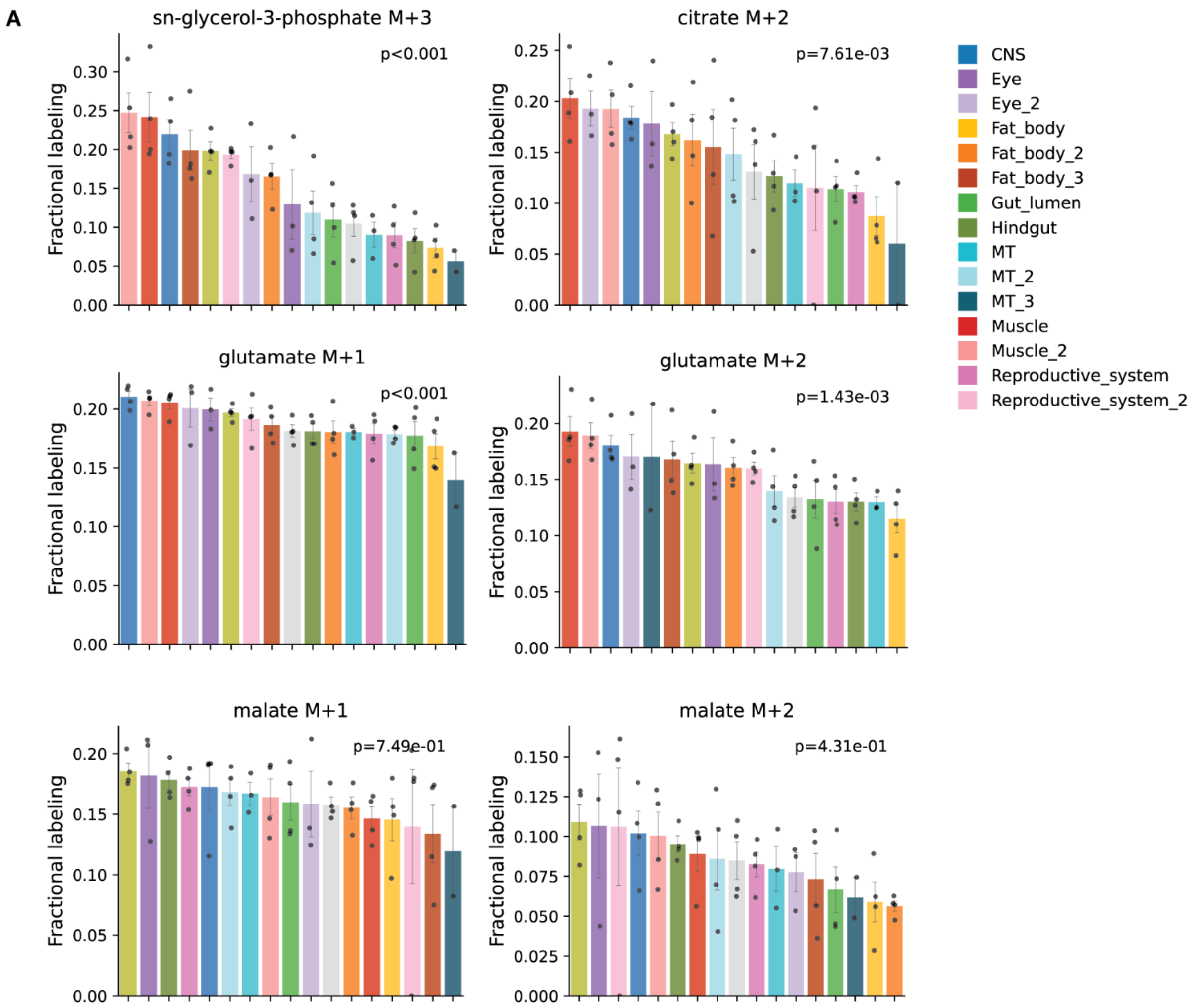

**Fig. S20. Glucose-derived fractional labeling of metabolites across annotated tissue regions.**

Tissue-level fractional labeling of isotopologues that varied significantly across annotated regions, including G3P M+3, citrate M+2, and glutamate M+1/M+2, malate M+1/M+2 across annotated regions. Bars indicate mean ± SEM; points indicate biological replicates. *P* values were calculated by one-way ANOVA.

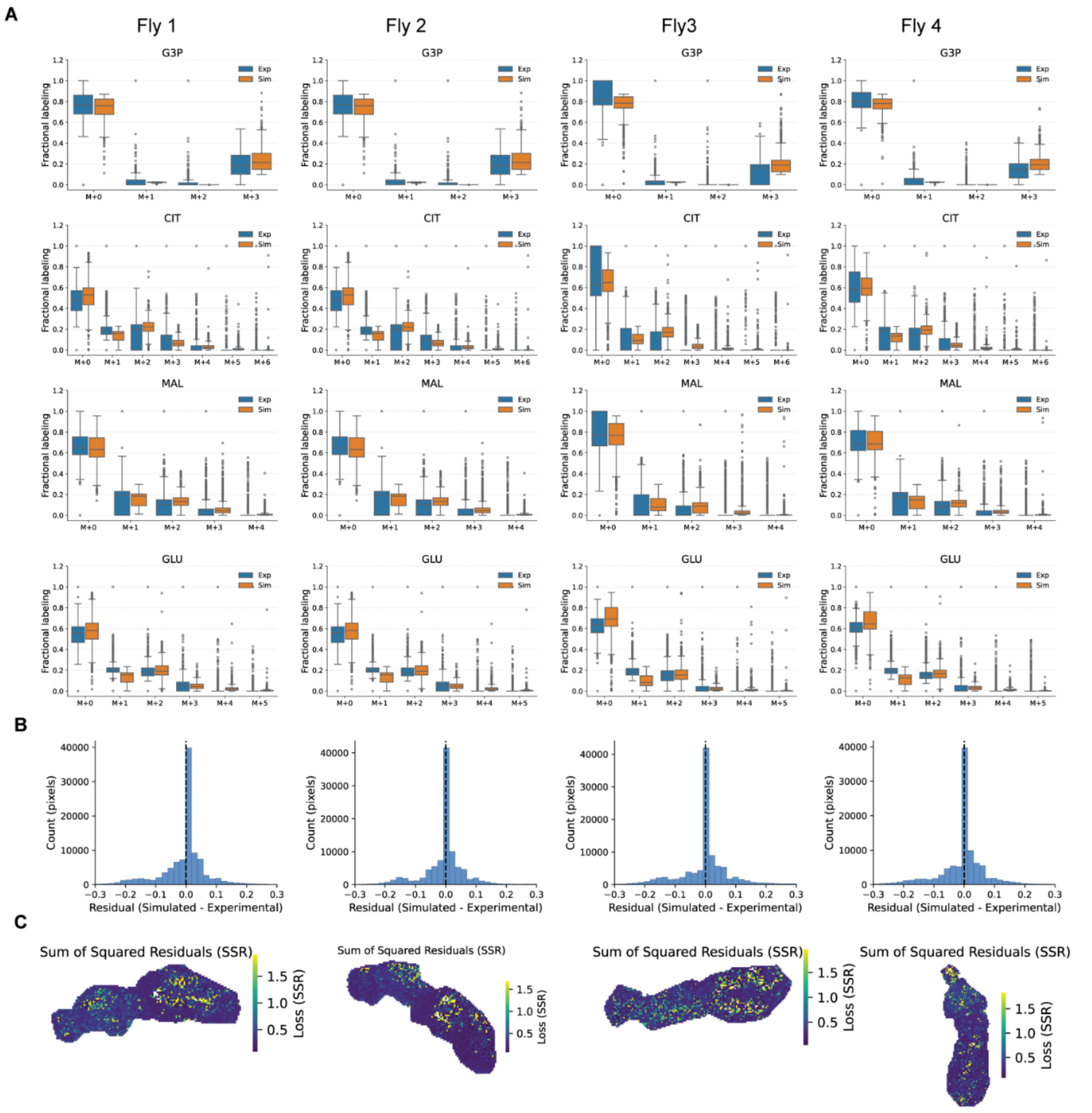

**Fig. S21. Validation of tissue-level $^{13}C$-metabolic flux analysis.**

**(A)** Comparison of experimentally measured and simulated mass isotopomer distributions for G3P, citrate, malate, and glutamate in individual flies fed [U-$^{13}C_6$]glucose. **(B)** Distributions of residuals between simulated and experimentally measured labeling values for each fly. **(C)** Spatial maps of summed squared residuals showing model fit across pixels in each fly.

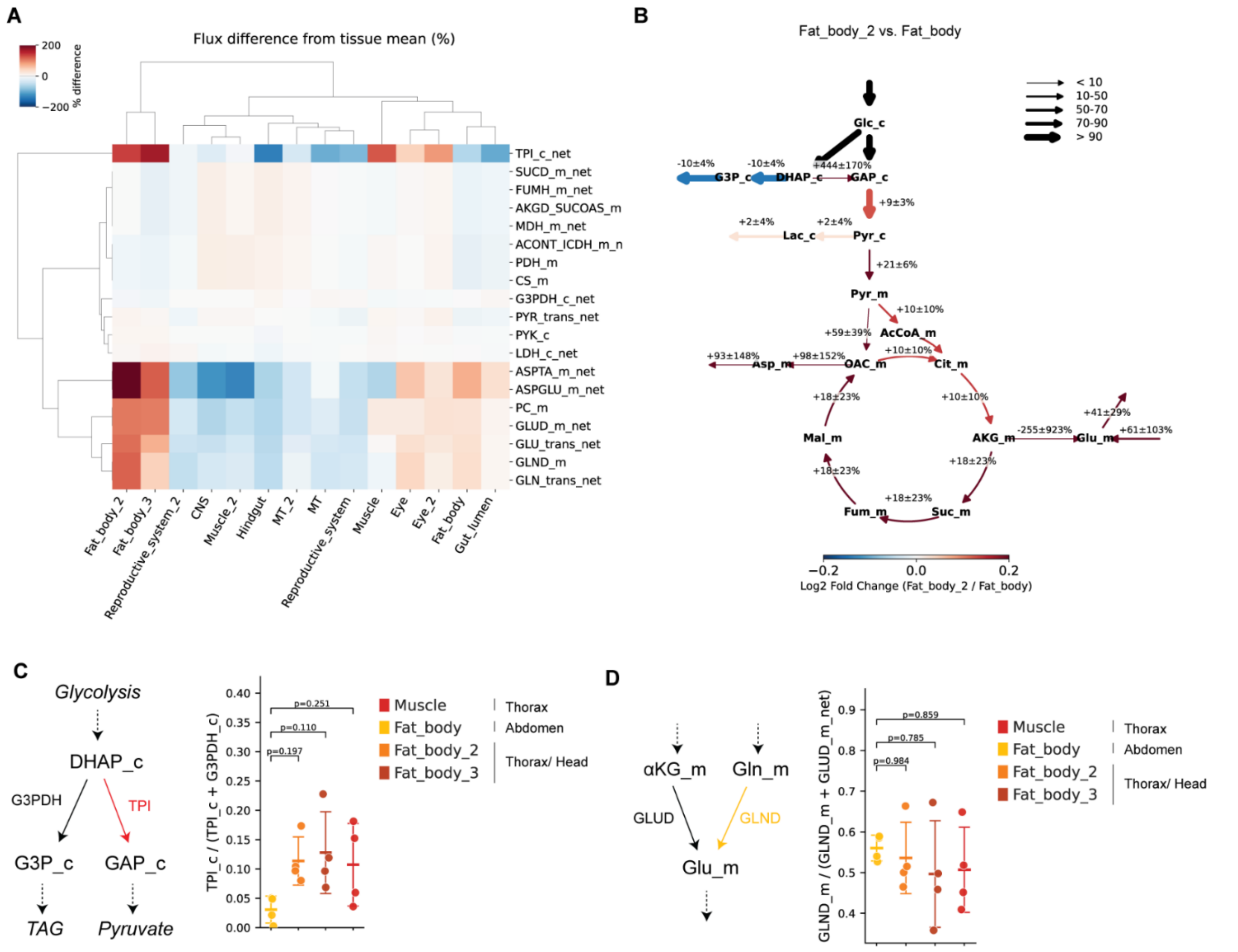

**Fig. S22. Differential flux estimates at tissue-level.**

**(A)** Flux differences across tissues. Values are percentage differences from the mean across tissues for each reaction; reactions with a difference greater than 5% in at least one tissue are shown. Colors are limited to ±200%. **(B)** Differential flux map comparing fat body_2 with abdominal fat body. Color indicates $\log_2$ fold change relative to Fat_body; edge width indicates net flux magnitude in Fat_body relative to glucose input. **(C)** Relative triose-phosphate isomerase (TPI) flux compared with glycerol-3-phosphate dehydrogenase (G3PDH) flux across fat body and muscle. **(D)** Relative glutaminase (GLND) flux compared with glutamate dehydrogenase (GLUD) flux across tissues.

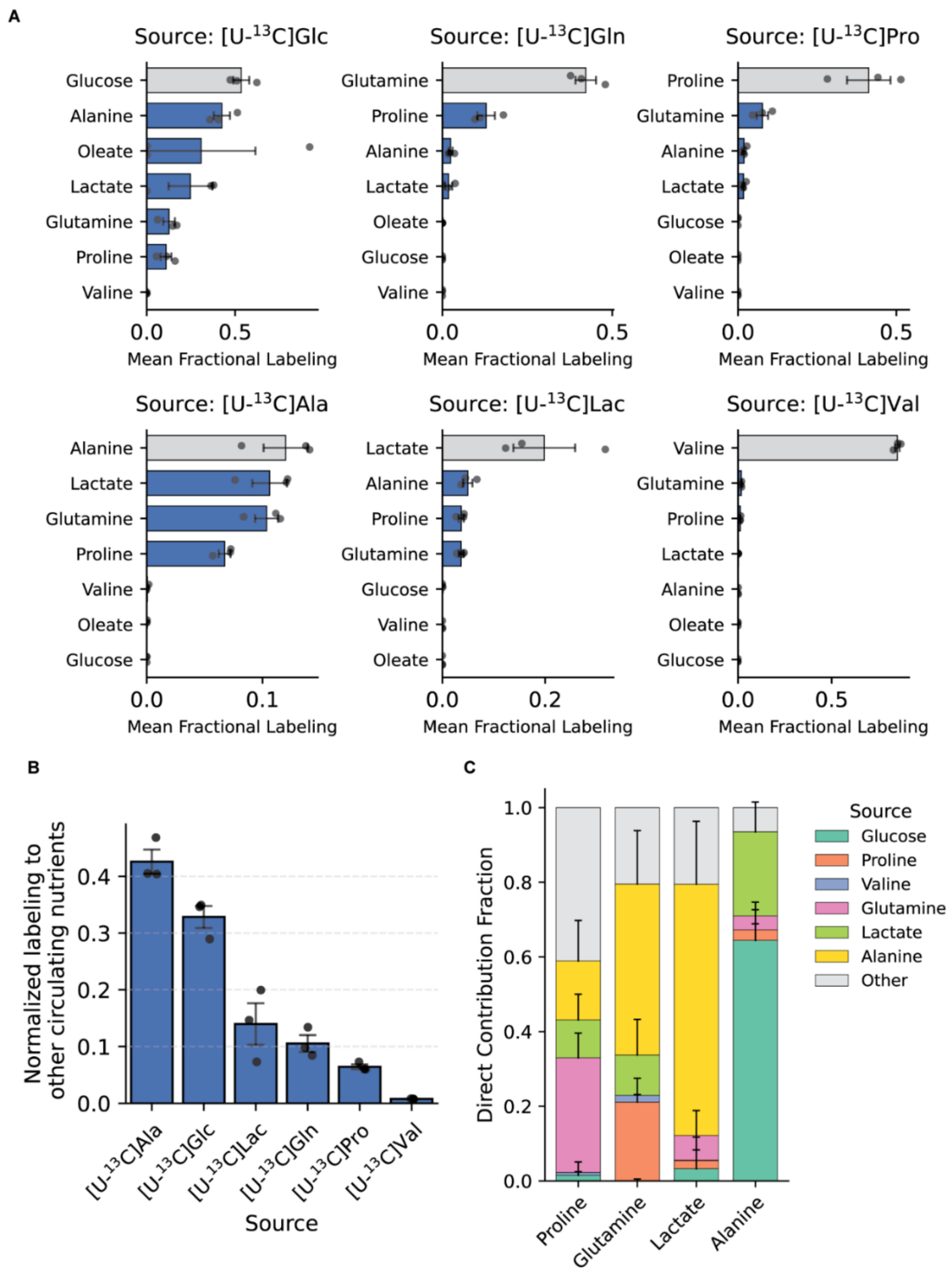

**Fig. S23. Nutrient interconversion in hemolymph.**

**(A)** Mean fractional labeling of circulating nutrients in hemolymph after feeding flies with the indicated $^{13}C$-labeled nutrient tracers. **(B)** Labeling of non-input circulating nutrients normalized to input nutrient labeling. **(C)** Direct contribution analysis estimating fractional contributions of circulating nutrients to proline, glutamine, lactate, and alanine in hemolymph. Bars indicate mean ± SEM; points indicate biological replicates.

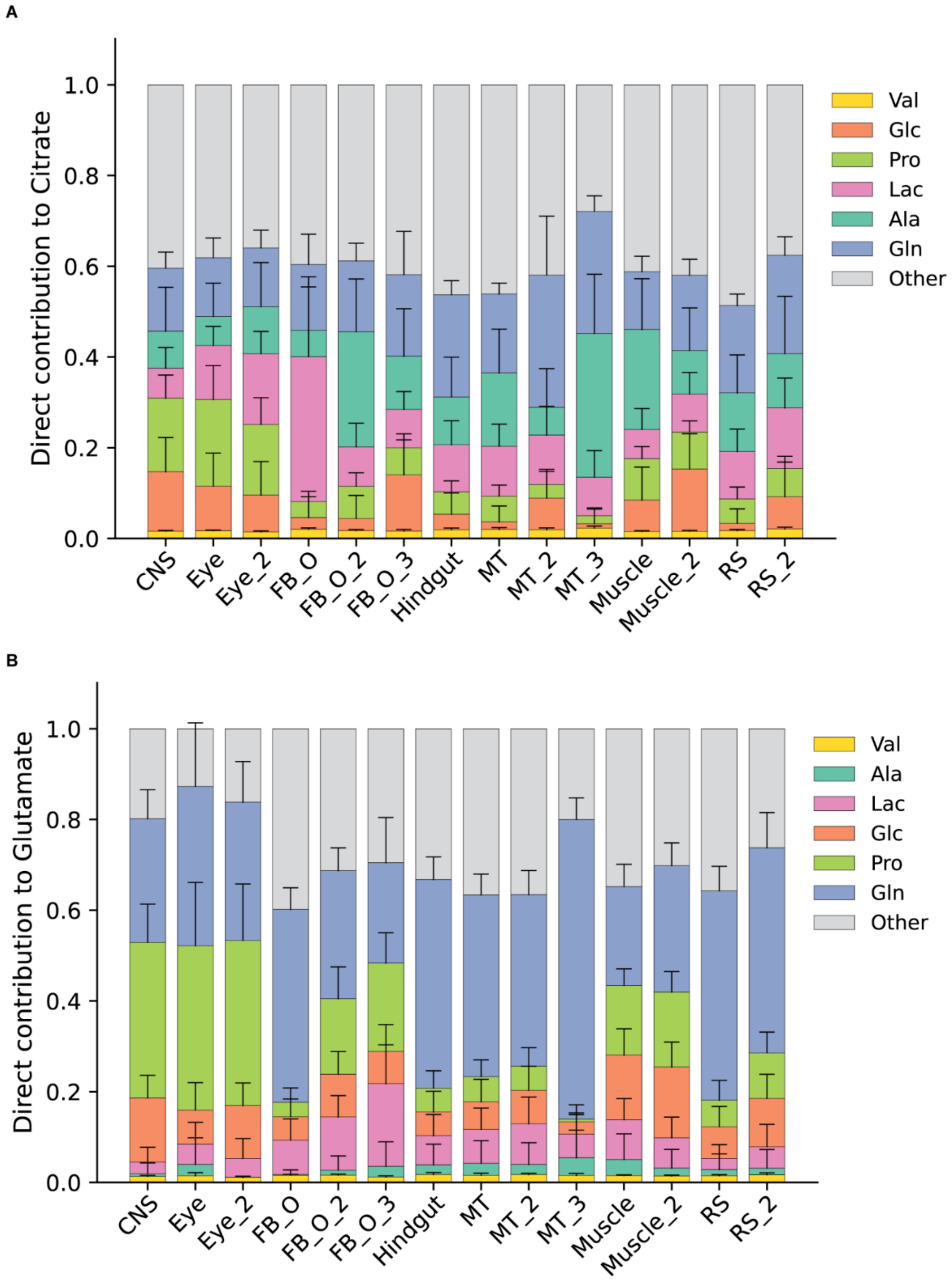

**Fig. S24. Direct nutrient contributions to citrate and glutamate across annotated tissue regions.**

**(A)** Estimated direct nutrient contributions to citrate across annotated tissue regions. **(B)** Estimated direct nutrient contributions to glutamate across annotated tissue regions. Bars indicate mean ± SEM.

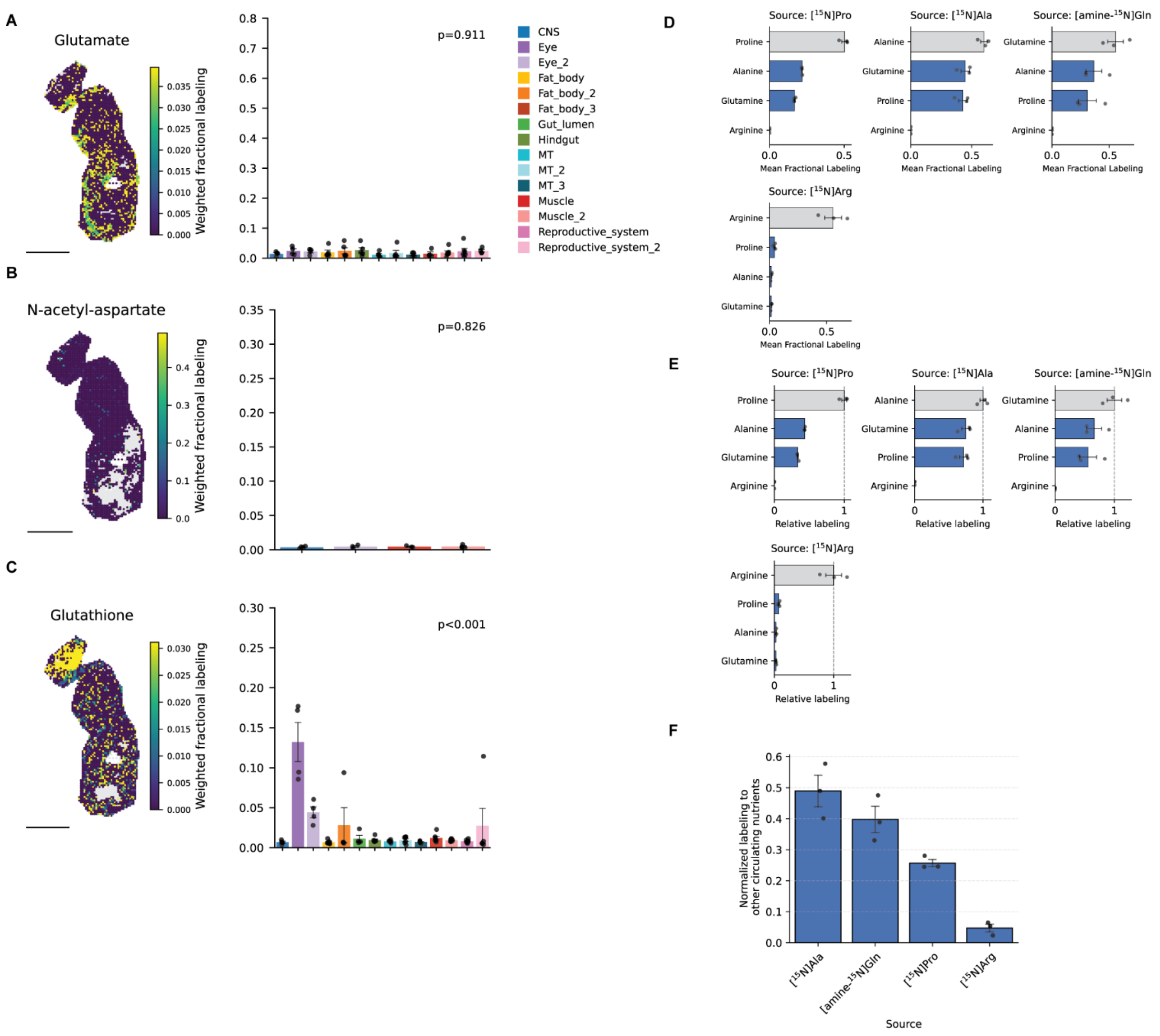

**Fig. S25. Arginine-derived tissue labeling and nitrogen interconversion in hemolymph.**

**(A)** Spatial map and tissue-level quantification of $^{15}$N labeling in glutamate from [$^{15}$N]arginine. **(B)** Spatial map and tissue-level quantification of $^{15}$N labeling in N-acetyl-aspartate from [$^{15}$N]arginine. **(C)** Spatial map and tissue-level quantification of $^{15}$N labeling in glutathione from [$^{15}$N]arginine. **(D)** Mean fractional labeling of circulating nitrogen-containing nutrients in hemolymph after feeding flies with the indicated $^{15}$N-labeled amino acid tracers. **(E)** Hemolymph labeling normalized to the input nutrient labeling. **(F)** Labeling of non-input circulating nutrients normalized to input nutrient labeling. Bars indicate mean ± SEM; points indicate biological replicates.

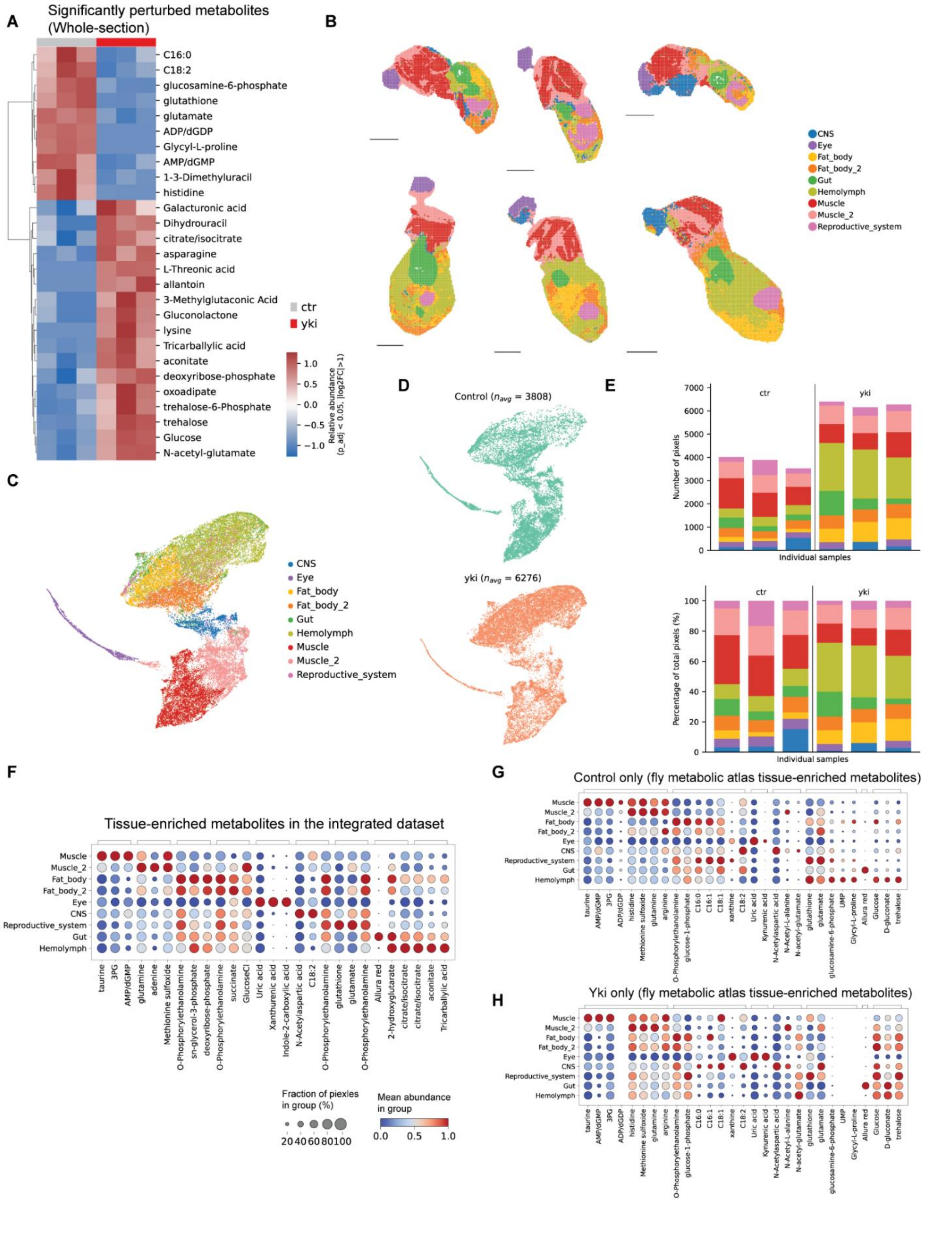
A
Significantly perturbed metabolites (Whole-section)
C16:0
C18:2
glucosamine-6-phosphate
glutathione
glutamate
ADP/dGDP
Glycyl-L-proline
AMP/dGMP
1-3-Dimethyluracil
histidine
Galacturonic acid
Dihydrouracil
citrate/isocitrate
asparagine
L-Threonic acid
allantoin
3-Methylglutaconic Acid
Gluconolactone
lysine
Tricarballylic acid
aconitate
deoxyribose-phosphate
oxoadipate
trehalose-6-Phosphate
trehalose
Glucose
N-acetyl-glutamate
ctr
yki
Relative abundance (p_adj < 0.05, |log2FC|>1)
B
CNS
Eye
Fat_body
Fat_body_2
Gut
Hemolymph
Muscle
Muscle_2
Reproductive_system
C
D
Control ($n_{avg}$ = 3808)
yki ($n_{avg}$ = 6276)
E
Number of pixels
Percentage of total pixels (%)
Individual samples
F
Tissue-enriched metabolites in the integrated dataset
Fraction of piexles in group (%)
Mean abundance in group
G
Control only (fly metabolic atlas tissue-enriched metabolites)
H
Yki only (fly metabolic atlas tissue-enriched metabolites)

**Fig. S26. Spatial metabolite profiling analysis in *yki*$^{3SA}$-tumor-bearing flies.**

**(A)** Differential metabolite abundance between control and *yki* flies at the whole-section level. TIC-based normalized metabolite intensities were averaged across all pixels within each fly to obtain one pseudobulk value per metabolite and biological replicate. Metabolites were compared between control and *yki* flies using Welch's two-sample *t*-test followed by Benjamini–Hochberg correction. Metabolites with adjusted $P < 0.05$ and absolute $\log_2$ fold change ≥ 1 are shown. Metabolite abundances were z-score normalized across biological replicates, and metabolites were hierarchically clustered. **(B)** Representative sagittal sections showing tissue annotations in control and *yki* flies. Colors indicate the annotated tissues. Scale bar, 500 µm. **(C)** UMAP of integrated spatial metabolite profiles from control and *yki* flies. Each point represents one pixel and is colored by tissue annotation. PCA was performed using seven principal components, followed by Harmony batch correction. A 50-nearest-neighbor graph was used for a UMAP generation with a minimum distance of 0.3, and Leiden clustering was performed at a resolution of 0.2. **(D)** Distribution of control and *yki* pixels within the same integrated UMAP embedding, shown separately by condition. $n_{avg}$ indicates the mean number of pixels per fly. **(E)** Number and proportion of annotated tissue pixels across individual control and *yki* sections. **(F)** Tissue-enriched metabolite patterns across annotated tissues in the integrated dataset. Metabolites were ranked across tissues using the Wilcoxon test, and the top three positively enriched metabolites with adjusted $P < 0.05$ for each tissue are shown. Dot size indicates the fraction of pixels with detectable metabolites within each tissue, and color indicates mean metabolite abundance scaled across tissues for each metabolite. **(G, H)** Tissue-enriched metabolites previously identified in the fly metabolic atlas, visualized separately in control (**G**) and *yki* (**H**) flies. Dot size indicates the fraction of pixels with detectable metabolites within each tissue, and color indicates relative mean metabolite abundance within each condition.

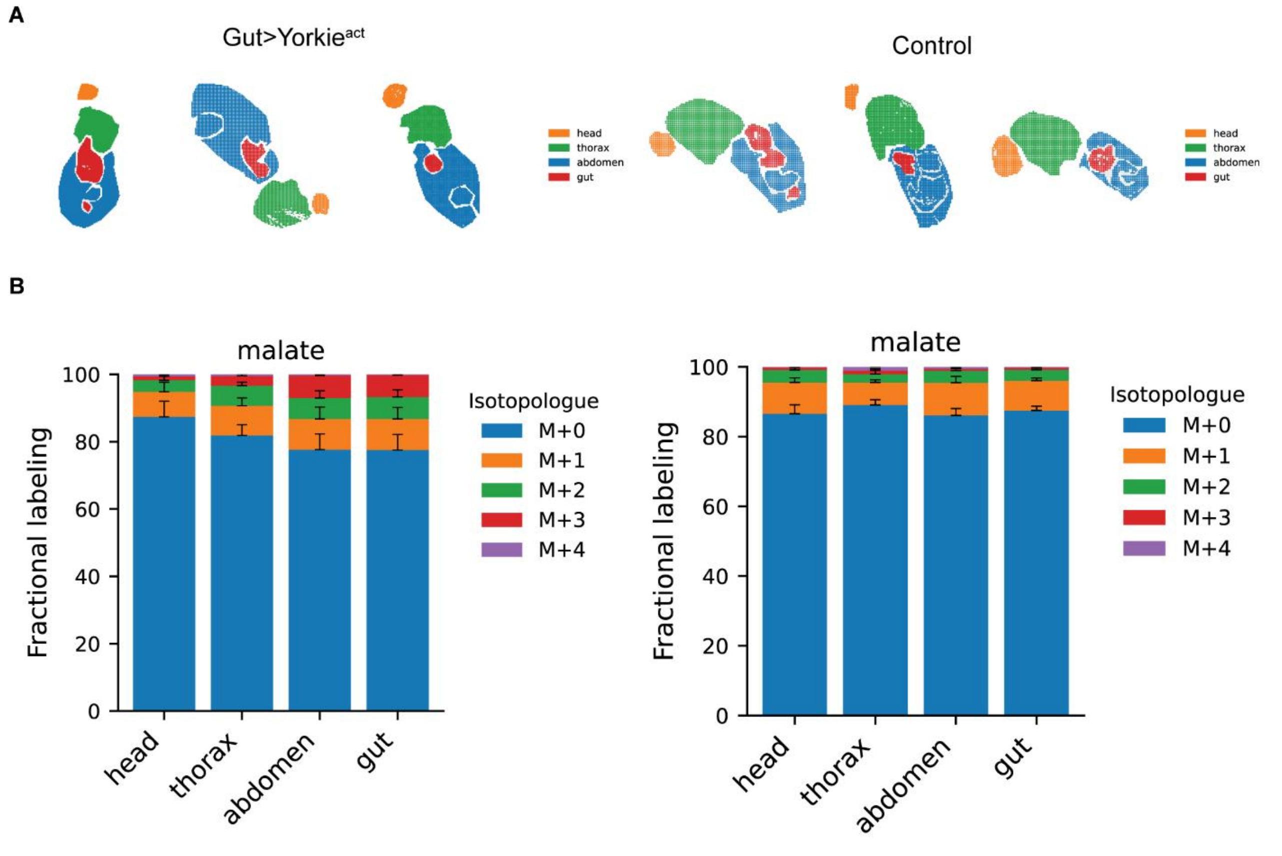

**Fig. S27. Malate fractional labeling from [U-$^{13}C_3$]alanine across anatomical regions in *yki*$^{3SA}$-tumor-bearing flies.**

**(A)** Anatomical region masks used to quantify head, thorax, abdomen, and gut labeling in *yki* and control flies. **(B)** Malate isotopologue distributions across anatomical regions in *yki* and control flies fed [U-$^{13}C_3$]alanine. Bars indicate mean ± SEM.

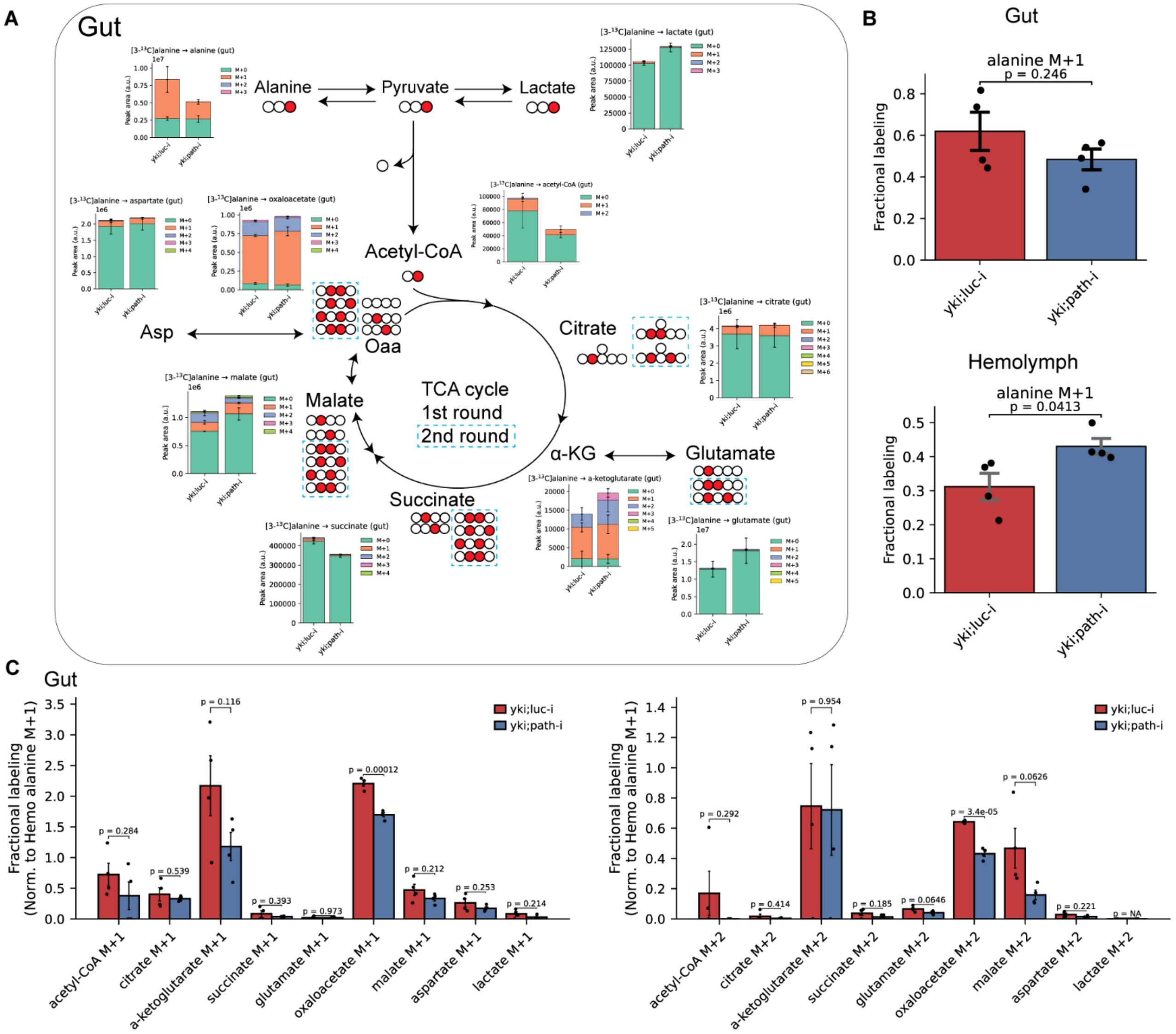

**Fig. S28. Evaluation of alanine contribution to TCA cycle metabolites in the gut of *yki;luc-i* and *yki;path-i* flies.**

**(A)** Schematic of the expected $^{13}$C-labeling patterns generated from [3-$^{13}$C]alanine around the TCA cycle. Peak areas of isotopologues of indicated metabolites in dissected guts following [3-$^{13}$C]alanine tracing in *yki;luc-i* and *yki;path-i* flies. Eight-day-old female *yki;luc-i* and *yki;path-i* flies were starved for 4 h, fed 0.5 % (w/v) [3-$^{13}$C]alanine for 5 h, and guts were dissected for metabolite extraction and LC-MS/MS analysis. Stacked bars show the mean peak area of each mass isotopologue and error bars indicate SEM. **(B)** Fractional labeling of the alanine M+1 isotopologue in gut or hemolymph from *yki;luc-i* and *yki;path-i* flies. Bars show mean ± SEM and dots represent biological replicates. Two-sided unpaired Student's *t*-tests were performed, with the corresponding *P* values shown. **(C)** Fractional labeling of indicated gut metabolite isotopologues in *yki;luc-i* and *yki;path-i* flies, normalized to the corresponding hemolymph alanine M+1 fractional labeling. Bars show mean ± SEM and dots represent biological replicates. Two-sided unpaired Student's *t*-tests were performed, with the corresponding *P* values shown.

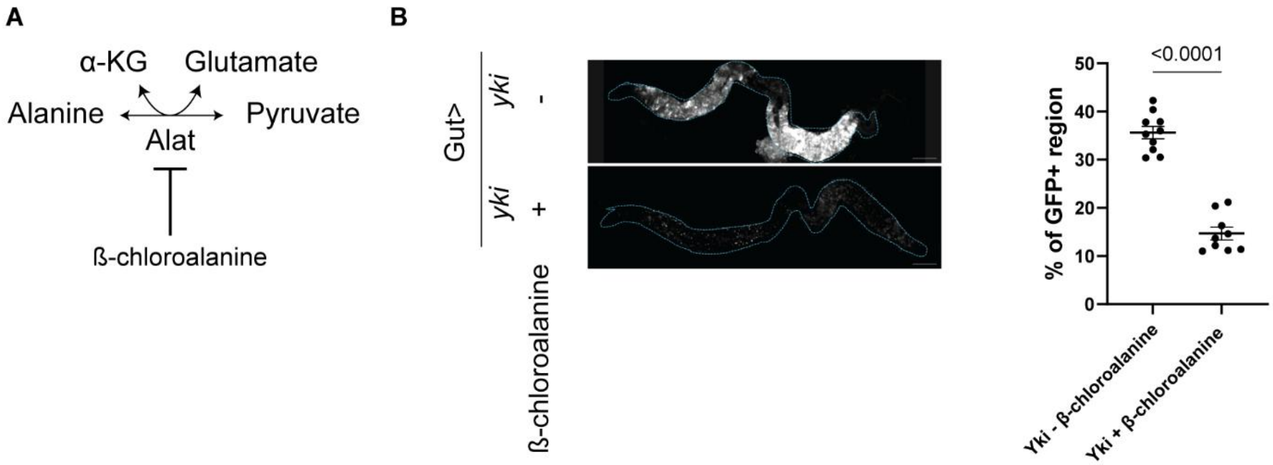

**Fig. S29. Pharmacological inhibition of alanine aminotransferase in *yki*$^{3SA}$-tumor-bearing flies.**

**(A)** Schematic of the alanine aminotransferase (*Alat*) reaction and its inhibition by β-chloroalanine. **(B)** Representative gut images and quantification of tumor growth in *esg*$^{ts}$>*yki* flies treated with or without β-chloroalanine. For inhibitor treatment, flies were transferred onto food containing 250 μM L-β-chloroalanine hydrochloride at day 0, simultaneously with tumor induction. Food was replaced every two days and gut GFP signals were quantified at day 8.

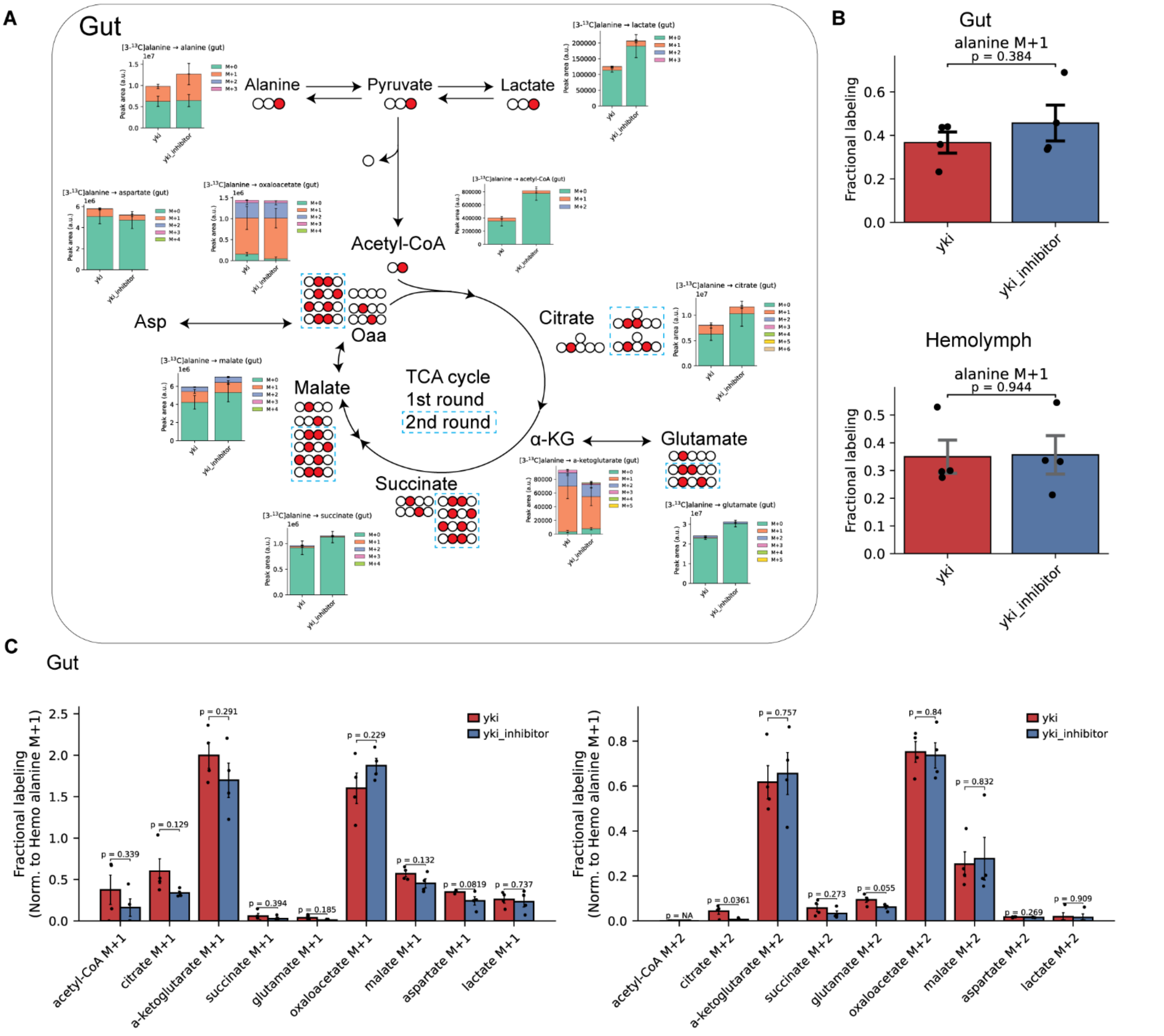

**Fig. S30. Evaluation of alanine contribution to TCA cycle metabolites following alanine aminotransferase inhibition in *yki*$^{3SA}$-tumor-bearing flies.**

**(A)** Schematic of the expected $^{13}$C-labeling patterns generated from [3-$^{13}$C]alanine around the TCA cycle, followed by peak areas of isotopologues of indicated metabolites in dissected guts from *yki*$^{3SA}$-tumor-bearing flies treated with or without β-chloroalanine. Stacked bars show the mean peak area of each mass isotopologue and error bars indicate SEM. **(B)** Fractional labeling of the alanine M+1 isotopologue in gut or hemolymph from *yki* flies treated with or without β-chloroalanine. Bars show mean ± SEM and dots represent biological replicates. Two-sided unpaired Student's *t*-tests were performed, with the corresponding *P* values shown. **(C)** Fractional labeling of indicated gut metabolite isotopologues in *yki* flies treated with or without β-chloroalanine, normalized to the corresponding hemolymph alanine M+1 fractional labeling. Bars show mean ± SEM and dots represent biological replicates. Two-sided unpaired Student's *t*-tests were performed, with the corresponding *P* values shown.

A

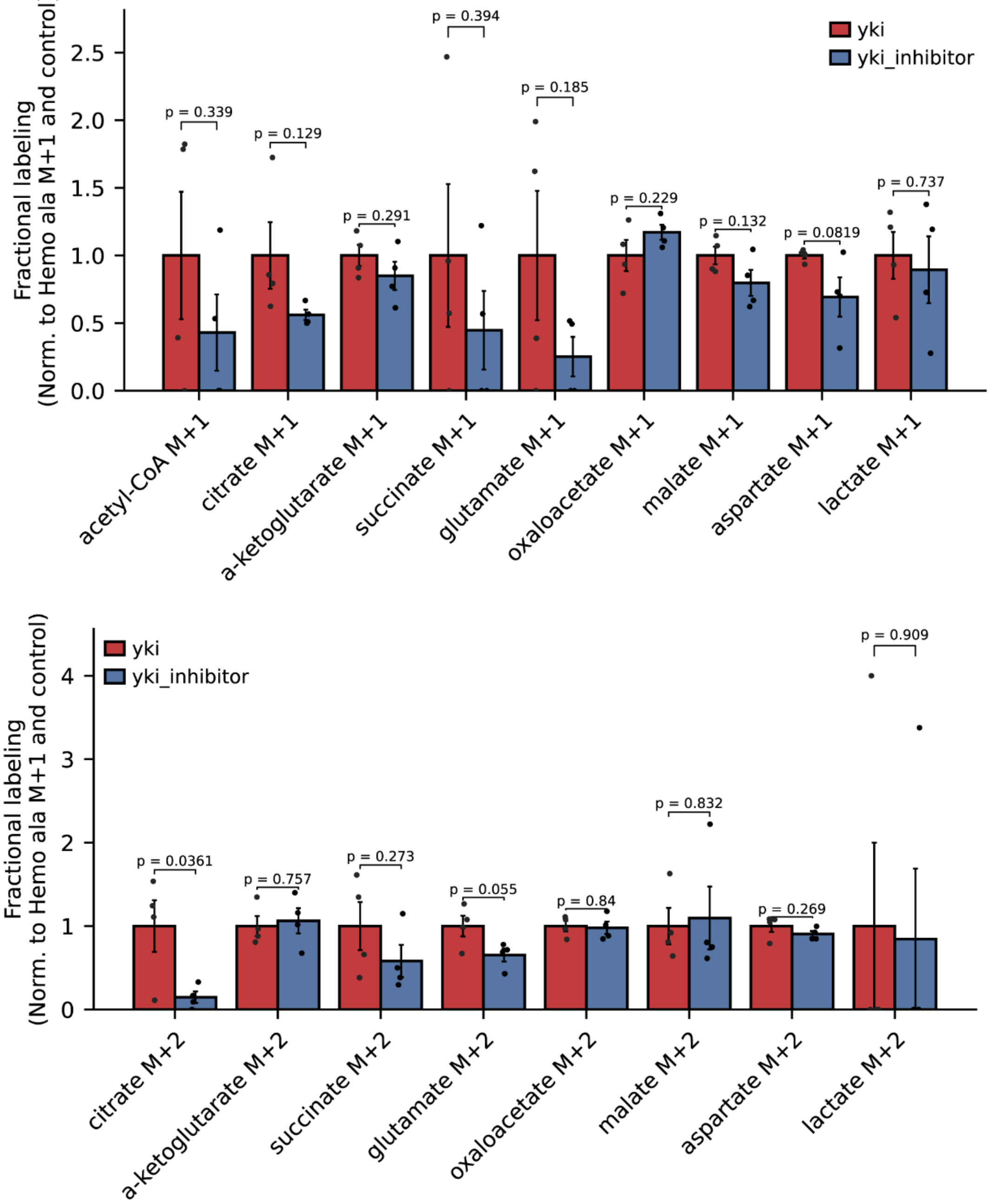

**Fig. S31. Normalized fractional labeling in response to alanine aminotransferase inhibition in *yki*$^{3SA}$-tumor-bearing flies.**

**(A–B)** Fractional labeling of M+1 **(A)** and M+2 **(B)** isotopologues of indicated gut metabolites following [3-$^{13}$C]alanine tracing in *yki* flies treated with or without β-chloroalanine, normalized to the corresponding hemolymph alanine M+1 fractional labeling and to the untreated *yki* control. Bars show mean ± SEM and dots represent biological replicates. Two-sided unpaired Student's *t*-tests were performed, with the corresponding P values shown.

**Table S1.** Characterization of Allura Red.

| Compound ID | *m/z* | ID method | *m/z* matching fragments (by MS2) |
|---|---|---|---|
| Allura Red | 452.035356 | $MS^2$ | 215.025778 |

**Table S2.** Concentrations of $^{13}C$- and $^{15}N$-labeled nutrients used for adult *Drosophila* feeding.

| Tracer used | Tracer concentration (mM) |
|---|---|
| [U-13C]glucose | 110 |
| [U-13C]glutamine | 100 |
| [U-13C]fructose | 110 |
| [U-13C]leucine | 110 |
| [U-13C]citrate | 50 |
| [U-13C]alanine | 110 |
| [U-13C]oleate | 10 |
| [U-13C]valine | 120 |
| [U-13C]lactate | 90 |
| [U-13C]proline | 130 |
| [amine-15N]glutamine | 100 |
| [15N]alanine | 110 |
| [15N]proline | 80 |
| [15N]arginine | 90 |

**Table S3.** Reference enriched metabolites from head, thorax, abdomen, and gut*.

| Region | Metabolite | Log2FC (region / average of other regions) | Adjusted p-value |
|---|---|---|---|
| Thorax | Adenylosuccinate | 3.2 | 5.8E-07 |
| | NADH | 3.0 | 5.4E-08 |
| | ADP-D-glucose | 2.6 | 1.9E-07 |
| | adenosine | 2.6 | 1.1E-04 |
| | AMP | 2.4 | 2.7E-08 |
| | homocysteic acid | 2.2 | 3.8E-02 |
| | Geranyl-PP | 2.2 | 4.6E-03 |
| | acetoacetyl-CoA-posi | 2.2 | 3.3E-05 |
| | Thiamine pyrophosphate | 2.2 | 3.8E-07 |
| | guanosine 5-diphosphate,3-diphosphate | 2.1 | 1.0E-02 |
| | thiamine-phosphate | 1.96 | 2.1E-03 |
| | adenine | 1.96 | 1.9E-04 |
| Head | 4-aminobutyrate | 2.8 | 3.8E-07 |
| | N-acetyl-L-aspartic acid | 2.7 | 6.8E-06 |
| | GMP | 2.6 | 8.9E-07 |
| | Ascorbic acid | 2.6 | 4.6E-06 |
| | 2-Isopropylmalic acid | 2.5 | 1.2E-05 |
| | N6-Acetyl-L-lysine | 2.4 | 0.0346 |
| | pantothenate | 2.4 | 4.7E-05 |
| | Phosphorylcholine | 2.2 | 0.00029 |
| | sarcosine | 2.2 | 6.4E-06 |
| | alanine | 2.1 | 4.8E-06 |
| | cytidine | 2.1 | 2.8E-05 |
| | glyoxylate | 2.1 | 0.00335 |
| | Indole-3-carboxylic acid | 2.1 | 0.02383 |
| Gut | methylsuccinate | 4.2 | 2.1E-03 |
| | thymidine | 4.1 | 5.4E-08 |
| | betaine | 3.6 | 6.7E-04 |
| | pyridoxine | 3.5 | 3.1E-06 |
| | quinolinate | 3.3 | 3.2E-03 |
| | Uric acid | 2.9 | 3.7E-05 |
| | N-acetyl-glutamine | 2.8 | 7.5E-04 |
| | homocysteine | 2.5 | 8.6E-03 |
| | anthranilate | 2.5 | 9.7E-05 |
| | p-aminobenzoate | 2.4 | 7.2E-05 |
| | 2,3-Diphosphoglyceric acid | 2.4 | 1.3E-03 |
| | D-glucarate | 2.4 | 1.3E-03 |
| | phosphocreatine | 2.4 | 6.7E-03 |
| Abdomen without gut | S-adenosyl-L-methioninamine | 2.9 | 1.3E-03 |
| | 5-methyl-THF | 1.1 | 2.1E-04 |
| | xanthosine | 1.0 | 3.9E-03 |
| | hypoxanthine | 1.0 | 2.4E-03 |
| | homocysteic acid | 0.9 | 3.8E-02 |
| | nicotinamide | 0.7 | 4.4E-03 |

| | | | |
|---|---|---|---|
| | cysteine | 0.5 | 5.2E-03 |
| | UDP-nega | 0.4 | 1.6E-02 |
| | methylnicotinamide | 0.4 | 1.3E-02 |
| | xanthine | 0.4 | 4.8E-03 |
| | 1-Methyladenosine | 0.3 | 2.6E-02 |
| | ethanolamine | 0.2 | 2.3E-03 |
| | UMP | 0.1 | 1.0E-02 |

*Full list of enriched metabolite sets are provided in the **Data_S1J**.

**Table S4.** Overview of the iterative annotation workflow.

| **Round** | **Primary focus** | **Goal** | **Related tables** |
|---|---|---|---|
| 1 | All 37 computational clusters | Generate initial tissue assignments using enriched metabolite profiles and spatial locations of clusters. | S5 |
| 2 | Eye- and MT-associated annotations | Separate anatomically distinct eye- and MT-associated pixels using anatomical region masks. | S6 |
| 3 | MT-associated regions | Resolve MT-related and previously uncertain abdominal regions. | S7 |
| 4 | Ovary and fat body | Resolve these abdominal regions. | S8 |
| 5 | Interstitial and unassigned abdominal regions | Resolve whether unresolved pixels were associated with neighboring MT, fat body, or hindgut regions. | S9 |
| 6 | Reproductive and surrounding abdominal regions | Resolve reproductive-system, ovary, and neighboring abdominal annotations. | S10 |
| 7 | Gut lumen annotations | Resolve gut lumen labels generated in different rounds and determine whether they should be retained, combined, or refined. | S11 |
| 8 | Thoracic and adjacent muscle regions | Resolve accumulated muscle-, thoracic fat body-, and interface-associated annotations. | S12 |
| 9 | CNS and fat body regions | Resolve related CNS and fat body labels and additional head- and thorax-associated patterns. | S13 |
| 10 | Thoracic edge, cuticle, and neighboring regions | Re-evaluate provisional thoracic edge and cuticle annotations relative to muscle, gut lumen, and fat body. | S14 |
| 11 | MT and adjacent gut regions | Re-evaluate accumulated MT-related labels and determine which could be consolidated or reassigned. | S15 |

| | | | |
|---|---|---|---|
| 12 | Hemolymph and unresolved neighboring regions | Re-evaluate the provisional hemolymph annotation because its enriched metabolite profile showed limited hemolymph support and partial MT features. | S16 |
| 13 | Gut lumen, fat body, and neighboring regions | Re-evaluate remaining related or uncertain working annotations before finalization. | S17 |
| 14 | All working annotations | Perform the final consolidation, reassignment, uncertainty review, and nomenclature standardization. | S18 |

**Table S5.** Round 1 - initial cluster annotations.

| Cluster(s) | Initial annotation | Anatomical region | Spatial pattern | Top enriched metabolites (log2 fold change versus other clusters)* | Description |
|---|---|---|---|---|---|
| 0, 26 | MT_eye_surrounding | Head and abdomen | Located near the eye and the eye–thorax interface in the head. In the abdomen, these clusters appeared as small localized regions, often near the posterior end. | Cluster 0: uric acid (3.0), kynurenic acid (2.8), xanthurenic acid (1.6), indole-2-carboxylic acid (1.4).<br>Cluster 26: uric acid (3.9), xanthurenic acid (2.3), indole-2-carboxylic acid (2.0), kynurenic acid (1.4). | Uric acid and related purine metabolites have known roles in *Drosophila* eye pigmentation (*74*). Uric acid, xanthurenic acid, and related metabolites were also enriched in the gut reference profile from our previous study (*27*). The combined head and abdominal distribution limited clear separation of eye- and MT-associated regions at this stage. |
| 1 | Eye_thorax_interface | Head and thorax | Located near the boundary between the posterior head and anterior thorax. | Uric acid (1.1), kynurenic acid (1.2), taurine (0.8). | Uric acid and kynurenic acid supported an eye-associated profile, whereas taurine may reflect contribution from adjacent thoracic tissue. Because this cluster occupied an anatomical boundary, its tissue composition may be mixed. |
| 2 | Eye_MT | Head and abdomen | Located mainly in the head in a pattern consistent with the eye region, but also detected in the | Uric acid (4.2), kynurenic acid (3.0), xanthurenic acid (2.2), indole-2-carboxylic acid (1.6), xanthine (0.5). | This cluster showed the strongest uric acid enrichment and a head location consistent with the eye. However, its presence in the abdomen suggested a |

| | | | | | |
|---|---|---|---|---|---|
| | | | abdomen in some sections. | | possible MT-associated component. The eye and MT assignments were not separated at this stage. |
| 3 | Thoracic_muscle | Thorax | Located mainly in the thorax and formed a region near the outer edge of the thoracic section. | Taurine (1.0), 3PG (1.0), AMP/dGMP (0.8), glycerol-3-phosphate (0.5). | Enrichment of taurine, AMP, and glycolytic metabolites supported a muscle-associated metabolic profile. AMP enrichment was also consistent with the thorax-enriched reference profile from our previous study (*27*). |
| 7 | Thoracic_fat_body | Thorax | Located within the thorax outside the major thoracic muscle regions. | C18:0 (0.9), taurine (0.8), hydroxyproline (0.8). | Fatty acid enrichment, together with its thoracic location, supported an initial thoracic fat body annotation. The metabolite evidence was limited, and the possible contribution of other lipid-rich cell types required further evaluation. |
| 8 | Thoracic_muscle_3 | Thorax | Located within a distinct thoracic spatial region. | Taurine (1.1), methionine sulfoxide (0.9), histidine (0.8), glutamine (0.7). | This cluster was initially assigned to a thoracic muscle region mainly based on its spatial location. Its metabolite evidence was less specific than that of other thoracic muscle clusters, and it was retained separately. |

| | | | | | |
|---|---|---|---|---|---|
| 9 | Thoracic_muscle_4 | Thorax | Located mainly within the interior of the thorax. | Taurine (1.83), 3PG (1.7), ADP/dGDP (1.7), AMP/dGMP (1.3), C18:2 (0.6). | Enrichment of taurine, adenylates, and 3PG supported a muscle-associated profile. AMP enrichment was consistent with the thorax reference profile from our previous study (*27*). The relative enrichment of C18:2 suggested some metabolic difference from the other thoracic muscle clusters. |
| 10 | Thoracic_muscle_5 | Thorax | Located mainly in the thorax and represented a small spatial population. | AMP/dGMP (2.6), ADP/dGDP (1.5), C18:2 (0.7). | Strong enrichment of adenylates supported a muscle-associated annotation. This cluster represented only 0.52% of all pixels and was retained separately because of its distinct spatial distribution. |
| 11 | Thorax_abdomen_interface | Thorax and abdomen | Located near the boundary between the posterior thorax and anterior abdomen. | Methionine sulfoxide (0.7), histidine (0.6), asparagine (0.6). | This cluster occupied an anatomical boundary and may contain contributions from several adjacent tissues. Its enriched metabolites did not support a more specific tissue assignment. |
| 4 | MT | Abdomen | Located in localized abdominal regions that could correspond to MT or adjacent | Isoxanthopterin (3.0), xanthurenic acid (2.4), indole-2-carboxylic acid (2.1), uric acid (0.8). | Enrichment of uric acid-related, pteridine, and kynurenine-pathway metabolites supported an excretory or gut-associated region. |

| | | | | | |
|---|---|---|---|---|---|
| | | | gut structures. | | Xanthurenic acid and indole-associated-carboxylic acid were enriched in the gut reference profile from our previous study (*27*). The MT assignment remained uncertain because nearby gut regions showed related metabolite profiles. |
| 21 | MT_2 | Abdomen | Located in localized abdominal regions that could correspond to MT or adjacent gut structures. | Isoxanthopterin (1.1), xanthurenic acid (1.0), glutathione (0.86). | The abdominal location and enrichment of isoxanthopterin and xanthurenic acid supported an MT- or gut-associated interpretation. |
| 5 | CNS | Mainly head and thorax | Located mainly in the head and along central nervous system-associated structures. Thoracic regions may represent the ventral nerve cord. | C18:0 (2.8), phosphoserine (1.8), N-acetylaspartic acid (1.8). | N-acetylaspartic acid enrichment was consistent with the head-enriched reference profile from our previous study (*27*) and with its known enrichment in the nervous system(*75*). The spatial pattern further supported a CNS annotation. |
| 6 | CNS_2 | Head and thorax | Located in spatial regions related to the CNS cluster but retained as a separate subregion. | N-acetylaspartic acid (3.0), N-acetyl-L-alanine (2.0), N-acetyl-glutamate (2.0), hydroxyproline (2.0). | Strong N-acetylaspartic acid enrichment supported a CNS annotation. This cluster was kept separate from cluster 5 because it showed different enriched metabolite profile. |

| | | | | | |
|---|---|---|---|---|---|
| 12, 15, 17, 19 | Gut_lumen | Abdomen | Located within gut- and lumen-associated regions of the abdomen across sections. | Cluster 12: Allura Red_214 (1.8), succinate (0.9), galacturonic acid (0.8).<br>Cluster 15: Allura Red (5.9), Allura Red_214 (2.2), O-phosphorylethanolamine (0.9).<br>Cluster 17: adenine (4.2), Allura Red_214 (4.1), hypoxanthine (3.3).<br>Cluster 19: O-phosphorylethanolamine (2.3), Allura Red_214 (1.8), glutamine (1.7). | Enrichment of Allura Red provided direct supporting evidence for gut lumen-associated regions. The four clusters were grouped under the same initial annotation but represented different spatial and metabolic patterns within the gut/lumen region. |
| 14 | Ovary | Abdomen | Located in a distinct abdominal region | GMP (2.2), glucosamine-6-phosphate (1.8), UMP (1.8), uridine (1.7), D-erythrose-4-phosphate (1.7), glutathione (1.6). | Enrichment of nucleotide-related metabolites, sugar phosphates, and glutathione, together with the abdominal location, supported an ovary- or reproductive system annotation. UMP enrichment was also consistent with the abdomen-enriched reference profile from our previous study (*27*). |
| 16 | Fat body | Abdomen | Broadly distributed in abdominal regions surrounding abdomen. | Xanthine (1.5), C14:0 (1.5), C16:0 (1.4), C18:1 (1.2), C16:1 (1.2), O-phosphorylethanolamine (1.0), glucose-1-phosphate (1.0). | Enrichment of several fatty acids, together with the broad abdominal distribution, supported a fat body annotation. Xanthine enrichment was consistent with the abdomen reference profile from our previous study (*27*). A contribution from |

| | | | | | |
|---|---|---|---|---|---|
| | | | | | oenocytes or other lipid-rich cell types could not be excluded. |
| 18 | Hemolymph | Abdomen | Sparsely distributed across the abdomen. Two sections contained a relatively large region that may instead represent the crop. | Glucose (4.2), GlucoseCl (3.9), trehalose (3.3), gluconolactone (3.2), deoxyribose-phosphate (3.0). | Trehalose is one of the major circulating sugar in insect hemolymph (*30*), and the enrichment of glucose and trehalose supported a hemolymph interpretation. However, the larger regions observed in two flies may represent crop contents. |
| 20 | Hindgut | Abdomen | Located in the posterior abdominal region. | Citrate/isocitrate (2.5), aconitate (2.4), galacturonic acid (1.9), glucose (1.8), succinate (1.1). | The posterior abdominal location and enrichment of galacturonic acid, which was also associated with gut/lumen clusters, supported a hindgut annotation. |
| 13, 22–25, 27–36 | Unannotated | Head, thorax, or abdomen | These clusters did not show a sufficiently clear or consistent anatomical pattern across sections to support a specific annotation. | No sufficiently specific enriched metabolite profile supported assignment to one tissue at this stage. | These clusters were retained as unannotated for further inspection. Most individual clusters each accounted for less than 0.1% of the 187,709 pixels, corresponding to fewer than approximately 188 pixels per cluster across the integrated dataset. In a typical section, these clusters were absent or represented by only a few pixels, with median counts ranging |

| | | | | | |
|---|---|---|---|---|---|
| | | | | | from 0 to 2 pixels per section. |

*All listed metabolites were significantly enriched by the Wilcoxon rank-sum test (adjusted $P < 1 \times 10^{-6}$).

**Table S6.** Round 2 - separation of eye and Malpighian tubule annotations.

| Round 1 annotation | Round 2 annotation | Anatomical criterion | Pixels, n (%)* | Description |
|---|---|---|---|---|
| Eye_MT | Eye | Outside the abdomen mask and closer to the head center than the abdomen center | 2,883 (89.7%) | Spatial location supported an eye annotation |
| Eye_MT | MT | Within the abdomen mask/region | 226 (7.0%) | Abdominal location supported an MT annotation |
| Eye_MT | Unannotated_MT | Outside the abdomen mask and closer to the abdomen center than the head center | 106 (3.3%) | Inconsistent with an Eye location but outside the abdomen mask and therefore not assigned as MT |
| MT | MT | Within the abdomen mask | 17,198 (79.5%) | Retained as MT based on abdominal location |
| MT | Eye_2 | Outside the abdomen mask and closer to the head center than the abdomen center | 365 (1.7%) | Spatial location was more consistent with an eye-associated region |
| MT | Unannotated_MT | Outside the abdomen mask and closer to the abdomen center than the head center | 4,076 (18.8%) | Outside the abdomen mask and therefore not retained as MT |
| MT_eye_surrounding | Eye_2 | Within the head mask, or outside both the head and abdomen masks and closer to the head center | 6,389 (53.4%) | Head-associated portion of the initial mixed annotation |
| MT_eye_surrounding | MT_2 | Within the abdomen mask/region | 4,266 (35.7%) | Abdominal portion of the initial annotation |
| MT_eye_surrounding | Unannotated_MT | Outside both the head and abdomen masks and closer to the abdomen center | 1,306 (10.9%) | Outside the abdomen mask and therefore not assigned as MT |

*Percentages were calculated relative to the total number of pixels within each Round 1 annotation.

**Table S7.** Round 3 - evaluation of the subclusters of MT, MT_2, and unannotated_MT annotations

| Subcluster | Round 2 annotation composition | Round 3 annotation | Pixels, n (%) | Sections containing pixels, n | Top enriched metabolites | Description |
|---|---|---|---|---|---|---|
| 0 | MT, 26%; MT_2, 65%; Unannotated_MT, 10% | Abdominal_unassigned_tissue | 12,588 (35.6%) | 68 | Histidine; methionine sulfoxide; glutamine; asparagine; glutathione | MT regions identified in Round 1 showed enrichment of purine-related metabolites, whereas this subcluster did not show the same profile. Because it was spatially located within the abdomen but lacked sufficient evidence for assignment to a specific tissue region, it was retained as an abdominal unassigned region for further evaluation. |
| 1 | MT, 84%; MT_2, 0%; Unannotated_MT, 16% | MT | 11,561 (32.7%) | 65 | Xanthurenic acid; isoxanthopterin; indole-2-carboxylic acid; kynurenic acid; hydroxyindoleacetic acid | Assigned as an MT region based on its spatial distribution and strong enrichment of MT-associated metabolites. |
| 2 | MT, 79%; MT_2, 1%; Unannotated_MT, 21% | oenocyte_fat_body | 4,668 (13.2%) | 65 | O-phosphorylethanolamine; C18:1; C16:0; C16:1; C18:2 | Assigned as an oenocyte/fat body region based on its spatial distribution, surrounding MT, and enrichment of O-phosphorylethanolamine and |

| | | | | | | |
|---|---|---|---|---|---|---|
| | | | | | | long-chain fatty acids. |
| 3 | MT, 12%; MT_2, 65%; Unannotated_MT, 23% | Interstitial_space | 6,501 (18.4%) | 46 | Uric acid; xanthine; galacturonic acid; Allura red_214; N-acetyl-glutamate | Assigned as an interstitial-space region because it occupied broad, continuous abdominal areas rather than the more discrete MT structures, and its enriched metabolite profile differed from that of the MT-associated subcluster. |
| 4 | MT, 10%; MT_2, 0%; Unannotated_MT, 90% | Unannotated_MT | 10 (0.03%) | 2 | Uric acid; fructose-1,6-bisphosphate; ATP; IMP; D-erythrose-4-phosphate | Retained as Unannotated_MT because the subcluster contained only 10 pixels distributed across two sections, providing insufficient evidence for a more specific tissue annotation. |
| 5 | MT, 0%; MT_2, 100%; Unannotated_MT, 0% | Unannotated_MT | 28 (0.08%) | 1 | N-acetylaspartic acid; sn-glycerol-3-phosphate; trehalose-6-phosphate; trehalose; citrate/isocitrate | Retained as Unannotated_MT because the subcluster contained only 28 pixels from a single section, providing insufficient evidence for a more specific tissue annotation. |

**Table S8.** Round 4 - evaluation of subclusters of Ovary and Fat_body annotations

| Subcluster | Round 3 annotation composition | Round 4 annotation | Pixels, n (%) | Sections containing pixels, n | Top enriched metabolites | Description |
|---|---|---|---|---|---|---|
| 0 | Ovary, 100% | Fat_body_outer_2 | 9,637 (27.0%) | 68 | Glutathione; histidine; citrate/isocitrate; methionine sulfoxide; aconitate | Assigned as Fat_body_outer_2 based on its peripheral spatial distribution across sections. Because all pixels were previously annotated as Ovary, this reassignment was considered tentative. |
| 1 | Fat_body, 7%; Ovary, 93% | Ovary | 8,474 (23.7%) | 67 | Glutathione; glucosamine-6-phosphate; UMP; glutamate; glycyl-L-proline | Retained as an ovary region based on its spatial distribution, strong correspondence with the previous Ovary annotation, and metabolite profile, including UMP and glutathione. |
| 2 | Fat_body, 96%; Ovary, 4% | Fat_body_outer | 4,877 (13.6%) | 67 | 1,3-Dimethyluracil; O-phosphorylethanolamine; succinate; GlucoseCl; malate | Assigned as Fat_body_outer based on its spatial distribution as a peripheral subregion of the broader Fat_body annotation. |
| 3 | Fat_body, 100% | Fat_body | 12,437 (34.8%) | 67 | O-phosphorylethanolamine; C18:2; C18:1; C16:0; C16:1 | Retained as a fat body region based on its spatial distribution and strong enrichment of O-phosphorylethanolamine and long-chain fatty acids. |
| 4 | Ovary, 100% | Fat_body | 58 (0.16%) | 4 | Glucose-1-phosphate; AMP/dGMP; D-erythrose-4-phosphate; UMP; GMP | Tentatively reassigned as Fat_body based on its spatial distribution and enrichment of glucose-1-phosphate, which was considered compatible with glycogen-associated metabolism (*76*). Because this subcluster contained less than 1% of the |

| | | | | | | |
|---|---|---|---|---|---|---|
| | | | | | | evaluated pixels and remained difficult to distinguish from reproductive-system regions, the assignment was considered low confidence and requires further evaluation. |
| 5 | Fat_body, 100% | Fat_body | 42 (0.12%) | 9 | Isoxanthopterin; O-phosphorylethanolamine; ornithine; fructose-1,6-bisphosphate; gluconolactone | Retained as Fat_body based on its previous annotation and enrichment of O-phosphorylethanolamine. Because this subcluster contained less than 1% of the evaluated pixels, the assignment was considered low confidence. |
| 6 | Fat_body, 100% | Fat_body | 64 (0.18%) | 1 | C16:1; C18:1; C18:2; C16:0; sn-glycerol-3-phosphate | Retained as Fat_body based on its previous annotation and enrichment of long-chain fatty acids. Because the subcluster was detected in only one section, the assignment was considered low confidence. |
| 7 | Fat_body, 100% | Fat_body_outer | 58 (0.16%) | 5 | Citrate/isocitrate; arginine; indole-2-carboxylic acid; Allura red; malate | Assigned as Fat_body_outer based on its spatial distribution. Because this subcluster contained less than 1% of the evaluated pixels, the assignment was considered low confidence. |
| 8 | Ovary, 100% | Fat_body_outer_2 | 29 (0.08%) | 1 | Citrate/isocitrate; GlucoseCl; trehalose; deoxyribose-phosphate; malate | Combined with subcluster 0 as Fat_body_outer_2 based on its similar spatial distribution. Because it contained |

| | | | | | | |
|---|---|---|---|---|---|---|
| | | | | | | only 29 pixels from one section, the assignment was considered low confidence. |
| 9 | Fat_body, 100% | Ovary | 45 (0.13%) | 2 | ADP/dGDP; isoxanthopterin; xanthurenic acid; indole-2-carboxylic acid; C14:0 | Reassigned as Ovary based on its spatial distribution. Because it contained less than 1% of the evaluated pixels and was previously annotated as Fat_body, the assignment was considered low confidence. |
| 10 | Fat_body, 100% | Fat_body | 14 (0.04%) | 2 | O-phosphorylethanolamine; ornithine; fructose-1,6-bisphosphate; gluconolactone; Allura red_214 | Retained as Fat_body based on its previous annotation and enrichment of O-phosphorylethanolamine. Because it contained only 14 pixels across two sections, the assignment was considered low confidence. |

**Table S9.** Round 5 - evaluation of abdominal subclusters from previously assigned and unassigned abdominal regions

| Subcluster | Round 4 annotation composition | Round 5 annotation | Pixels, n (%) | Sections containing pixels, n | Top enriched metabolites | Description |
|---|---|---|---|---|---|---|
| 0 | Hindgut, 99%; Abdominal_unassigned_tissue, 1% | Reproductive_system | 3,823 (9.6%) | 64 | Citrate/isocitrate; malate; aconitate; 2-hydroxyglutarate; arginine | Reassigned as Reproductive_system based on its spatial localization and enrichment of several TCA-cycle intermediates. In contrast, subcluster 6 showed the posterior distribution and uric acid-, xanthine-, and Allura Red-associated profile expected for hindgut, supporting reconsideration of the previous Hindgut annotation of subcluster 0. |
| 1 | Abdominal_unassigned_tissue, 100% | hemolymph | 8,183 (20.6 %) | 68 | Histidine; glutamate; deoxyribose-phosphate; methionine sulfoxide; glutamine | Provisionally assigned as hemolymph based on its broadly distributed abdominal pattern rather than a clearly bounded tissue morphology. Because its metabolite profile was not sufficiently tissue-specific, this assignment was retained |

| | | | | | | |
|---|---|---|---|---|---|---|
| | | | | | | as tentative and was evaluated again in subsequent rounds. |
| 2 | Hindgut, 99%; other, <1% | Reproductive_system | 559 (1.4%) | 21 | Citrate/isocitrate; deoxyribose-phosphate; glucose; GlucoseCl; trehalose | Reassigned as Reproductive_system based primarily on its spatial localization. Its limited section coverage and less distinctive metabolite profile made this assignment more tentative than that of subcluster 0. |
| 3 | MT, 64%; Abdominal_unassigned_tissue, 31%; oenocyte_fat_body, 4%; other, <1% | MT_outer | 6,289 (15.8 %) | 64 | Xanthurenic acid; indole-2-carboxylic acid; guanosine; kynurenic acid; hydroxyindole acetic acid | Assigned as MT_outer based on its spatial distribution around the periphery of the more core MT subcluster and enrichment of metabolites associated with Malpighian tubule regions, including xanthurenic acid and kynurenic acid. |
| 4 | MT, 100% | MT | 6,078 (15.3 %) | 65 | Isoxanthopterin; xanthurenic acid; indole-2-carboxylic acid; kynurenic acid; uric acid | Retained as MT based on its consistent spatial pattern, correspondence with the previous MT annotation, and strong enrichment of uric acid, |

| | | | | | | |
|---|---|---|---|---|---|---|
| | | | | | | isoxanthopterin, and kynurenine-pathway metabolites. |
| 5 | oenocyte_fat_body, 96%; Abdominal_unassigned_tissue, 3%; MT, 1% | Fat_body_2 | 4,230 (10.7%) | 64 | O-phosphorylethanolamine; C18:1; C16:0; C16:1; C18:2 | Assigned as Fat_body_2 based on its spatial distribution and strong enrichment of O-phosphorylethanolamine and long-chain fatty acids. The previous combined oenocyte_fat_body label was simplified, but the analysis did not definitively distinguish fat body from oenocyte contributions within this region. |
| 6 | Interstitial_space, 99%; other, <1% | Hindgut | 6,154 (15.5%) | 46 | Uric acid; xanthine; galacturonic acid; Allura Red_214; N-acetyl-glutamate | Reassigned as Hindgut based on its posterior rectal/hindgut spatial distribution and enrichment of uric acid, xanthine, galacturonic acid, and the ingested dye-related signal Allura Red_214. |
| 7 | Abdominal_unassigned_tissue, 71%; MT, 24%; oenocyte_fat_body, 5% | Abdominal_muscle | 2,848 (7.2%) | 62 | Taurine; AMP/dGMP; 3-phosphoglycerate; | Assigned as Abdominal_muscle based on its body-wall-associated spatial distribution and |

| | | | | | | |
|---|---|---|---|---|---|---|
| | | | | | isoxanthopterin; guanosine | enrichment of taurine, adenylates, and the glycolytic intermediate 3-phosphoglycerate. Its mixed Round 4 composition indicates that the region had not previously been cleanly separated from adjacent MT- and fat-body pixels. |
| 8 | Hindgut, 100% | Unannotated | 27 (0.1%) | 5 | Citrate/isocitrate; fructose-1,6-bisphosphate; ATP; IMP; D-erythrose-4-phosphate | Left unannotated because it contained very few pixels, occurred in only five sections, and did not form a sufficiently reproducible spatial pattern to support a tissue assignment. |
| 9 | MT, 55%; oenocyte_fat_body, 40%; Abdominal_unassigned_tissue, 5% | Crop | 371 (0.9%) | 16 | GlucoseCl; deoxyribose-phosphate; glucose; trehalose; citrate/isocitrate | Assigned as Crop based on its spatial localization and carbohydrate-rich metabolite profile, including glucose and trehalose. Because the subcluster was small, occurred in relatively few sections, and originated from several previous annotations, this |

| | | | | | | |
|---|---|---|---|---|---|---|
| | | | | | | assignment was considered tentative. |
| 10 | Interstitial_space, 100% | Abdominal_muscle | 87 (0.2%) | 7 | Taurine; inosine; guanosine; L-kynurenine; hypoxanthine | Assigned as Abdominal_muscle based primarily on its body-wall-associated spatial localization, with taurine enrichment providing additional support. The small number of pixels and limited section coverage made this assignment tentative. |
| 11 | Interstitial_space, 94%; Abdominal_unassigned_tissue, 6% | Fat_body_2 | 86 (0.2%) | 5 | Glucose-1-phosphate; ADP/dGDP; uric acid; trehalose-6-phosphate; UMP | Assigned as Fat_body_2 based primarily on its spatial correspondence with the broader Fat_body_2 region. The assignment remained tentative because the subcluster was small and detected in only five sections. |
| 12 | MT, 79%; Abdominal_unassigned_tissue, 17%; Interstitial_space, 3%; other, <1% | Fat_body_2 | 716 (1.8%) | 21 | Citrate/isocitrate; aconitate; 2-hydroxyglutarate; kynurenic acid; GlucoseCl | Assigned as Fat_body_2 based primarily on its spatial distribution relative to the larger fat-body region. Because the enriched metabolites |

| | | | | | | |
|---|---|---|---|---|---|---|
| | | | | | | were not uniquely diagnostic of fat body, this reassignment was considered tentative. |
| 13 | Abdominal_unassigned_tissue, 99%; oenocyte_fat_body, 1% | Unannotated | 90 (0.2%) | 8 | Malate; 2-hydroxyglutarate; glutamate; taurine; fumarate | Left unannotated because the subcluster was small, inconsistently detected across sections, and lacked a sufficiently distinctive spatial pattern or metabolite profile to support a tissue assignment. |
| 14 | Interstitial_space, 93%; oenocyte_fat_body, 7% | Hindgut | 161 (0.4%) | 6 | Uric acid; aspartate; succinate; citrate/isocitrate; glutamate | Assigned as Hindgut based on its posterior spatial localization and enrichment of uric acid. Because it contained relatively few pixels and occurred in only six sections, this assignment was considered tentative. |

**Table S10.** Round 6 - evaluation of subclusters from reproductive and surrounding abdominal regions.

| Subcluster | Round 5 annotation composition | Round 6 annotation | Pixels, n (%) | Sections containing pixels, n | Top enriched metabolites | Description |
|---|---|---|---|---|---|---|
| 0 | Thorax_abdomen_interface, 100% | Abdominal_muscle_2 | 8,667 (16.2%) | 68 | Taurine; histidine; 3PG; sn-glycerol-3-phosphate; N-acetylaspartic acid | Assigned as Abdominal_muscle_2 based on its edge-associated spatial distribution and enrichment of taurine and glycolytic metabolites. |
| 1 | Thorax_abdomen_interface, 100% | Abdominal_muscle_2 | 2,105 (3.9%) | 57 | Taurine; isoxanthopterin; xanthurenic acid; uric acid; AMP/dGMP | Assigned as Abdominal_muscle_2 based on its spatial adjacency to cluster 0 and enrichment of taurine. The mixed metabolite profile indicated possible contributions from adjacent tissues. |
| 2 | Thorax_abdomen_interface, 50%; Fat_body_outer_2, 37%; Crop, 11%; Reproductive_system, 1%; other, <1% | Abdominal_muscle_2 | 5,906 (11.1%) | 68 | Taurine; histidine; glutathione; glutamine; methionine sulfoxide | Assigned as Abdominal_muscle_2 based on its spatial adjacency to cluster 0 and 1, with taurine enrichment providing additional support. |
| 3 | Ovary, 66%; Fat_body_outer_2, 33%; other, <1% | Fat_body_2 | 12,225 (22.9%) | 67 | O-phosphorylethanolamine; glucose-1-phosphate; glutathione; C16:1; C18:1 | Assigned as Fat_body_2 based on its spatial distribution and enrichment of O-phosphorylethanolamine and long-chain fatty acids. |
| 4 | Fat_body_outer_2, 94%; Crop, 5%; Ovary, 1% | Oenocyte | 2,286 (4.3%) | 68 | Glutathione; UMP; glycyl-L-proline; glucosamine-6-phosphate; trehalose-6-phosphate | Provisionally assigned as oenocyte. Because its metabolite profile was not considered characteristic of oenocytes, this assignment was re-evaluated in later rounds. |
| 5 | Crop, 100% | Crop | 3,420 (6.4%) | 36 | GlucoseCl; deoxyribose-phosphate; | Retained as Crop based on its spatial distribution and enrichment of |

| | | | | | | |
|---|---|---|---|---|---|---|
| | | | | | glucose; trehalose; oxoadipate | glucose- and trehalose-associated signals. |
| 6 | Reproductive_system, 98%; Crop, 1%; Ovary, 1% | Reproductive_system | 3,686 (6.9%) | 64 | Citrate/isocitrate; aconitate; 2-hydroxyglutarate; malate; succinate | Retained as Reproductive_system based on its spatial distribution and enrichment of multiple TCA-cycle intermediates. |
| 7 | Crop, 83%; Fat_body_outer_2, 13%; Ovary, 3%; other, <1% | MT_2 | 8,829 (16.5%) | 68 | Xanthurenic acid; isoxanthopterin; uric acid; indole-2-carboxylic acid; guanosine | Reassigned as MT_2 based on its tubular spatial distribution and strong enrichment of uric acid, isoxanthopterin, and kynurenine-pathway metabolites. |
| 8 | Reproductive_system, 96%; Ovary, 3%; Fat_body_outer_2, 1% | Reproductive_system | 669 (1.3%) | 26 | Citrate/isocitrate; deoxyribose-phosphate; glucose; GlucoseCl; aconitate | Retained as Reproductive_system based on its enrichment of TCA cycle metabolites. Its limited section coverage and mixed metabolites made this assignment more tentative than that of subcluster 6. |
| 9 | Fat_body_2, 99%; Crop, 1% | Gut_lumen_2 | 4,273 (8.0%) | 64 | Xanthurenic acid; isoxanthopterin; O-phosphorylethanolamine; indole-2-carboxylic acid; uric acid | Reassigned as Gut_lumen_2 based primarily on its internal luminal spatial distribution. Because its previous annotation and metabolite profile were mixed, this was retained as an assignment for later re-evaluation. |
| 10 | Crop, 100% | Crop | 369 (0.7%) | 16 | Isoxanthopterin; GlucoseCl; deoxyribose-phosphate; glucose; xanthurenic acid | Retained within the Crop working annotation based on its spatial distribution and carbohydrate-associated signals. Its small size and mixed metabolite profile made the assignment tentative. |
| 11 | Crop, 100% | Crop | 30 (0.1%) | 2 | Isoxanthopterin; GlucoseCl; glucose; trehalose; | Retained as Crop based on its previous annotation, but considered tentative because it contained |

| | | | | | | |
|---|---|---|---|---|---|---|
| | | | | | deoxyribose-phosphate | very few pixels and occurred in only two sections. |
| 12 | Fat_body_2, 100% | Unannotated | 86 (0.2%) | 5 | Uric acid; isoxanthopterin; glucose-1-phosphate; ADP/dGDP; trehalose-6-phosphate | Left unannotated because it was small, occurred in only five sections, and lacked a reproducible spatial pattern supporting a tissue assignment. |
| 13 | Thorax_abdomen_interface, 100% | Unannotated | 52 (0.1%) | 2 | Hypoxanthine; 2-hydroxyglutarate; adenine; 1,3-dimethyluracil; fumarate | Left unannotated because it contained very few pixels, occurred in only two sections, and lacked sufficient spatial evidence for assignment. |
| 14 | Fat_body_2, 100% | Crop | 712 (1.3%) | 21 | Citrate/isocitrate; isoxanthopterin; xanthurenic acid; aconitate; kynurenic acid | Consolidated into the Crop working annotation based primarily on its adjacency to crop region. Because its metabolite profile was mixed and not diagnostic of crop, this assignment was considered provisional and re-evaluated later. |
| 15 | Ovary, 100% | Fat_body_2 | 46 (0.1%) | 2 | ADP/dGDP; C16:1; C18:1; C16:0; glucose-1-phosphate | Assigned as Fat_body_2 based on its spatial adjacency to fat body and enrichment of long-chain fatty acids. The assignment remained tentative because the subcluster contained very few pixels and occurred in only two sections. |

**Table S11.** Round 7 **-** evaluation of subclusters from the gut lumen annotations.

| Subcluster | Round 6 annotation composition | Round 7 annotation | Pixels, n (%) | Sections containing pixels, n | Top enriched metabolites | Description |
|---|---|---|---|---|---|---|
| 0 | Gut_lumen, 100% | CNS_3 | 5,281 (33.7%) | 67 | Histidine; sn-glycerol-3-phosphate; arginine; N-acetylaspartic acid; glutamine | Reassigned as CNS_3 based on its spatial localization and enrichment of N-acetylaspartic acid. |
| 1 | Gut_lumen, 100% | Gut_lumen | 2,192 (14.0%) | 61 | Allura Red; glucose-1-phosphate; glutathione; AMP/dGMP; ADP/dGDP | Retained as Gut_lumen based on its internal luminal distribution and strong enrichment of the ingested dye Allura Red. |
| 2 | Gut_lumen, 100% | Interstitial_space | 3,145 (20.1%) | 55 | Adenine; pyroglutamic acid; hypoxanthine; guanine; succinate | Reassigned as Interstitial_space based primarily on its diffuse spatial distribution between recognizable tissue structures rather than within a clearly defined gut lumen. This was retained as a tentative annotation for later re-evaluation. |
| 3 | Gut_lumen, 100% | Unannotated | 393 (2.5%) | 21 | O-phosphorylethanolamine; fructose-1,6-bisphosphate; ribose-phosphate; D-erythrose-4-phosphate; uridine | Left unannotated because it lacked a sufficiently consistent spatial pattern or distinctive metabolite profile to support a tissue assignment. |

| | | | | | | |
|---|---|---|---|---|---|---|
| 4 | Gut_lumen_2, 92%; Gut_lumen, 8% | Gut | 4,620 (29.5%) | 64 | Xanthurenic acid; isoxanthopterin; uric acid; indole-2-carboxylic acid; guanosine | Assigned as Gut based primarily on its spatial distribution along gut-associated tissue rather than within the dye-rich lumen. Because its metabolite profile was not uniquely diagnostic of gut, this was retained as a tentative annotation for later re-evaluation. |
| 5 | Gut_lumen, 100% | Unannotated | 21 (0.1%) | 3 | Malate; fructose-1,6-bisphosphate; ribose-phosphate; uridine; ornithine | Left unannotated because it contained very few pixels, occurred in only three sections, and lacked sufficient spatial evidence for assignment. |
| 6 | Gut_lumen, 100% | Unannotated | 23 (0.1%) | 1 | Pyroglutamic acid; fructose-1,6-bisphosphate; ribose-phosphate; uridine; ornithine | Left unannotated because it was detected in only one section and did not represent a reproducible tissue. |

**Table S12.** Round 8 - evaluation of subclusters from thoracic and adjacent muscle annotations.

| Subcluster | Round 7 annotation composition | Round 8 annotation | Pixels, n (%) | Sections containing pixels, n | Top enriched metabolites | Description |
|---|---|---|---|---|---|---|
| 0 | Eye_thorax_interface, 75%; Thoracic_muscle, 16%; Abdominal_muscle, 4%; Abdominal_muscle_2, 3%; Thoracic_muscle_3, 1% | Gut_lumen_3 | 2,256 (2.9%) | 53 | Uric acid; xanthurenic acid; kynurenic acid; indole-2-carboxylic acid; xanthine | Reassigned as Gut_lumen_3 based on its gut/MT associated metabolites. Because most pixels were previously assigned to Eye_thorax_interface and the enriched metabolites were not unique metabolites of gut lumen, this assignment was considered tentative. |
| 1 | Thoracic_muscle_4, 82%; Thoracic_muscle, 16%; Thoracic_muscle_3, 2% | Muscle | 19,532 (25.4%) | 68 | Taurine; AMP/dGMP; 3-phosphoglycerate; ADP/dGDP; C16:1 | Consolidated as muscle based on its broad thoracic distribution and enrichment of taurine, adenylates, and the glycolytic intermediate 3-phosphoglycerate. |
| 2 | Thoracic_fat_body, 48%; Thoracic_muscle, 39%; Abdominal_muscle_2, 11%; Thoracic_muscle_4, 1% | Fat_body_3 | 15,208 (19.8%) | 68 | O-phosphorylethanolamine; C16:0; C18:1; C18:2; 1,3-dimethyluracil | Assigned as Fat_body_3 based on its spatial distribution and enrichment of O-phosphorylethanolamine and long-chain fatty acids. |
| 3 | Thoracic_muscle_3, 55%; Thoracic_muscle, 31%; Abdominal_muscle_2, 13%; Thoracic_muscle_4, 1% | Thorax_edge | 24,153 (31.5%) | 68 | Histidine; methionine sulfoxide; asparagine; arginine; glutamine | Assigned as Thorax_edge based primarily on its peripheral spatial distribution along the thoracic boundary. The annotation was considered tentative. |
| 4 | Thoracic_muscle_5, 100% | Fat_body_3 | 978 (1.3%) | 66 | AMP/dGMP; C18:2; glucose-1-phosphate; ADP/dGDP; C16:0 | Reassigned as Fat_body_3 based on its spatial adjacency with the cluster 2 region and enrichment of several long-chain fatty acids. Because the metabolite profile also contained adenylates, this reassignment was considered tentative. |

| | | | | | | |
|---|---|---|---|---|---|---|
| 5 | Abdominal_muscle, 64%; Abdominal_muscle_2, 35% | Gut_lumen_2 | 4,382 (5.7%) | 64 | Xanthurenic acid; isoxanthopterin; uric acid; indole-2-carboxylic acid; guanosine | Reassigned as Gut_lumen_2 based primarily on its location between thorax and abdomen. This assignment was retained as provisional. |
| 6 | Abdominal_muscle_2, 99%; Thoracic_muscle_3, 1% | Gut_lumen_2 | 10,257 (13.4%) | 68 | Glutathione; malate; glutamate; tricarballylic acid; glycyl-L-proline | Reassigned as Gut_lumen_2 based on its spatial adjacency with subcluster 5. The metabolite profile was not unique to gut lumen. This reassignment was considered tentative |

**Table S13.** Round 9 - evaluation of subclusters from CNS and fat body annotations.

| Subcluster | Round 8 annotation composition | Round 9 annotation | Pixels, n (%) | Sections containing pixels, n | Top enriched metabolites | Description |
|---|---|---|---|---|---|---|
| 0 | CNS, 51%; CNS_2, 48%; Fat_body_3, 1% | CNS | 12,475 (23.5%) | 68 | N-acetylaspartic acid; hydroxyproline; N-acetyl-L-alanine; C18:0; sn-glycerol-3-phosphate | Consolidated as CNS based on its spatial localization in head, its predominant composition of the previous CNS and CNS_2 annotations, and enrichment of N-acetylaspartic acid. |
| 1 | Fat_body_3, 96%; CNS, 3%; CNS_2, 1% | Thoracic_cuticle | 15,647 (29.5%) | 68 | Taurine; 3-phosphoglycerate; AMP/dGMP; glutamine; methionine sulfoxide | Reassigned as Thoracic_cuticle based primarily on its peripheral spatial distribution around the thoracic region. Because most pixels were previously annotated as Fat_body_3 and the metabolite profile contained muscle features, this was retained as a tentative annotation for further evaluation. |
| 2 | Fat_body, 53%; CNS_3, 22%; Fat_body_outer, 21%; CNS, 5% | Fat_body | 23,846 (45.0%) | 67 | O-phosphorylethanolamine; glucose-1-phosphate; malate; glutamate; guanine | Consolidated as Fat_body based on its spatial distribution on the outer edge, continuity with the previous Fat_body and Fat_body_outer annotations, and enrichment of O-phosphorylethanolamine and glucose-1-phosphate. |
| 3 | Fat_body_3, 100% | Thoracic_cuticle | 977 (1.8%) | 66 | AMP/dGMP; 3-phosphoglycerate; ADP/dGDP; arginine; citrate/isocitrate | Reassigned as Thoracic_cuticle based primarily on its thin peripheral spatial distribution and adjacency to subcluster 1. Because its metabolite profile was not assigned as cuticle previously, this |

| | | | | | | |
|---|---|---|---|---|---|---|
| | | | | | | reassignment was considered provisional. |
| 4 | Fat_body, 100% | Fat_body | 58 (0.1%) | 4 | Glucose-1-phosphate; UMP; D-erythrose-4-phosphate; AMP/dGMP; GMP | Retained as Fat_body based on its previous annotation. The assignment remained tentative because the subcluster was small and detected in only four sections. |
| 5 | CNS_2, 100% | CNS | 29 (0.1%) | 3 | N-acetylaspartic acid; uridine; ribose-phosphate; D-erythrose-4-phosphate; N-acetyl-L-proline | Consolidated as CNS based on its previous CNS_2 annotation and enrichment of N-acetylaspartic acid. The assignment remained tentative because the subcluster contained very few pixels and occurred in only three sections. |

**Table S14**. Round 10 **-** re-evaluation of subclusters from thoracic edge, cuticle, and neighboring tissue annotations.

| Subcluster | Round 9 annotation composition | Round 10 annotation | Pixels, n (%) | Sections containing pixels, n | Top enriched metabolites | Description |
|---|---|---|---|---|---|---|
| 0 | Gut_lumen_3, 94%; Thoracic_cuticle, 5%; Muscle, 1% | Gut_lumen_3 | 2,026 (2.0%) | 54 | Uric acid; xanthurenic acid; kynurenic acid; indole-2-carboxylic acid; xanthine | Retained as Gut_lumen_3 based primarily on its spatial correspondence with the previous Gut_lumen_3 annotation. |
| 1 | Thorax_edge, 45%; Muscle, 37%; Thoracic_cuticle, 17%; Gut_lumen_3, 1% | Muscle | 52,138 (51.6%) | 68 | Taurine; AMP/dGMP; 3-phosphoglycerate; sn-glycerol-3-phosphate; ADP/dGDP | Consolidated as Muscle because the previously separate Thorax_edge, Muscle, and Thoracic_cuticle annotations formed a single coherent subcluster, without clear evidence supporting their continued separation. Its broad muscle-associated spatial distribution and enrichment of taurine, adenylates, and glycolytic metabolites further supported the Muscle annotation. |
| 2 | Thoracic_cuticle, 98%; Muscle, 2%; other, <1% | Thoracic_fat_body | 6,038 (6.0%) | 68 | O-phosphorylethanolamine; C18:2; C18:1; C16:0; taurine | Reassigned as Thoracic_fat_body based on its spatial distribution near the edge and enrichment of O-phosphorylethanolamine and long-chain fatty acids. |
| 3 | Fat_body, 100% | Fat_body | 23,646 (23.4%) | 67 | O-phosphorylethanolamine; glucose-1-phosphate; | Retained as Fat_body based on complete correspondence with the previous annotation, its spatial distribution, |

| | | | | | | |
|---|---|---|---|---|---|---|
| | | | | | C16:0; malate; guanine | and enrichment of O-phosphorylethanolamine and long-chain fatty acids. |
| 4 | Thoracic_cuticle, 100% | Unannotated | 977 (1.0%) | 66 | AMP/dGMP; ADP/dGDP; C18:2; N-acetylaspartic acid; arginine | Left unannotated because its spatial pattern and mixed metabolite profile did not provide sufficient evidence for a distinct cuticle-associated or other tissue assignment. |
| 5 | Gut_lumen_2, 98%; Thoracic_cuticle, 2% | Gut_lumen_2 | 4,689 (4.6%) | 64 | Xanthurenic acid; isoxanthopterin; uric acid; indole-2-carboxylic acid; guanosine | Retained as Gut_lumen_2 based on its correspondence with the previous annotation and its spatial distribution within the broader Gut_lumen_2-associated region. |
| 6 | Gut_lumen_2, 86%; Thoracic_cuticle, 7%; Thorax_edge, 4%; Fat_body, 2%; other, <1% | Gut_lumen_2 | 11,517 (11.4%) | 68 | Glutathione; histidine; citrate/isocitrate; malate; tricarballylic acid | Retained as Gut_lumen_2 based primarily on its spatial continuity with the broader Gut_lumen_2 region. Its mixed Round 9 composition and nonspecific metabolite profile indicated that this remained a working annotation. |
| 7 | Fat_body, 100% | Fat_body | 58 (0.1%) | 4 | Glucose-1-phosphate; UMP; D-erythrose-4-phosphate; GMP; C16:1 | Retained as Fat_body based on its previous annotation and spatial correspondence with the broader Fat_body region. The assignment remained tentative because the subcluster was small and detected |

| | | | | | | |
|---|---|---|---|---|---|---|
| | | | | | | in only four sections. |
| 8 | Gut_lumen_3, 100% | Unannotated | 19 (<0.1%) | 1 | N-acetylaspartic acid; hydroxyproline; N-acetyl-L-alanine; N-acetyl-glutamate; 3-hydroxybenzoic acid | Left unannotated because it contained very few pixels, was detected in only one section, and did not represent a reproducible tissue. |

**Table S15.** Round 11 **-** re-evaluation of subclusters from MT and adjacent gut annotations.

| Subcluster | Round 10 annotation composition | Round 11 annotation | Pixels, n (%) | Sections containing pixels, n | Top enriched metabolites | Description |
|---|---|---|---|---|---|---|
| 0 | MT_2, 86%; MT_outer, 14% | Hemolymph | 10,061 (38.9%) | 68 | Histidine; glutathione; methionine sulfoxide; glutamate; glutamine | Provisionally reassigned as Hemolymph based on its broadly distributed spatial pattern and lack of strong enrichment for the uric acid- and isoxanthopterin-associated MT profile. Because its metabolite profile did not show strong evidence of hemolymph, this was retained as a working annotation for later re-evaluation. |
| 1 | MT, 51%; MT_outer, 49% | MT | 7,259 (28.1%) | 62 | Xanthurenic acid; indole-2-carboxylic acid; isoxanthopterin; kynurenic acid; uric acid | Consolidated as MT based on its spatial distribution and strong enrichment of MT-associated metabolites. The combined composition indicated that the previous MT and MT_outer labels represented closely related regions. |
| 2 | MT, 83%; MT_outer, 14%; MT_2, 2% | MT | 2,813 (10.9%) | 64 | Isoxanthopterin; uric acid; xanthurenic acid; indole-2-carboxylic acid; guanosine | Consolidated as MT based on its predominant previous MT composition, spatial distribution, and enrichment of established MT-associated metabolites. |
| 3 | Gut, 89%; MT_outer, 9%; MT, 1% | Fat_body_3 | 5,123 (19.8%) | 66 | O-phosphorylethanolamine; C16:0; C16:1; C18:1; C18:2 | Reassigned as Fat_body_3 based on its spatial distribution and enrichment of O-phosphorylethanolamine and long-chain fatty acids. These features were more consistent with fat body than with |

| | | | | | | |
|---|---|---|---|---|---|---|
| | | | | | | the previous Gut or MT-associated annotations. |
| 4 | MT_outer, 100% | Unannotated | 250 (1.0%) | 16 | Taurine; xanthurenic acid; guanosine; glutathione; isoxanthopterin | Left unannotated because its spatial pattern and mixed muscle- and MT-associated metabolite profile did not provide sufficient evidence for a confident tissue assignment. |
| 5 | MT_outer, 100% | Unannotated | 9 (<0.1%) | 1 | Inosine; ribose-phosphate; guanosine; kynurenic acid; glucosamine-6-phosphate | Left unannotated because it contained very few pixels and was detected in only one section. |
| 6 | MT_outer, 100% | Unannotated | 135 (0.5%) | 7 | GlucoseCl; glucose; deoxyribose-phosphate; trehalose; citrate/isocitrate | Left unannotated because it was small, detected in relatively few sections. |
| 7 | MT_2, 100% | Unannotated | 11 (<0.1%) | 1 | Mandelic acid; histidine; 3-hydroxybenzoic acid; arginine; 1,3-dimethyluracil | Left unannotated because it contained very few pixels and was detected in only one section. |
| 8 | MT_2, 77%; MT_outer, 23% | Unannotated | 155 (0.6%) | 7 | GlucoseCl; deoxyribose-phosphate; glucose; succinate; 1,3-dimethyluracil | Left unannotated because it contained very few pixels and was detected in only one section. |
| 9 | Unannotated_MT, 100% | Unannotated | 10 (<0.1%) | 2 | Uric acid; fructose-1,6-bisphosphate; ATP; D-erythrose-4-phosphate; ornithine | Left unannotated because it contained very few pixels and was detected in only one section. |
| 10 | Unannotated_MT, 100% | Unannotated | 28 (0.1%) | 1 | N-acetylaspartic acid; sn-glycerol-3-phosphate; trehalose-6-phosphate; trehalose; citrate/isocitrate | Retained as Unannotated because it was detected in only one section and showed a mixed metabolite profile without a reproducible spatial pattern. |

**Table S16.** Round 12 – re-evaluation of subclusters from hemolymph and neighboring unresolved annotations.

| Subcluster | Round 11 annotation composition | Round 12 annotation | Pixels, n (%) | Sections containing pixels, n | Top enriched metabolites | Description |
|---|---|---|---|---|---|---|
| 0 | Unannotated, 100% | Interstitial_space | 976 (2.8%) | 66 | AMP/dGMP; C18:2; ADP/dGDP; taurine; 3-phosphoglycerate | Assigned as Interstitial_space based primarily on its location between recognizable tissue regions. Its mixed muscle- and lipid-associated metabolite profile did not support a more specific tissue assignment, so this remained a tentative annotation that will be re-evaluated later round. |
| 1 | Gut_lumen_2, 97%; Hemolymph, 3% | Gut_lumen_2 | 4,653 (13.5%) | 64 | Isoxanthopterin; xanthurenic acid; uric acid; guanosine; indole-2-carboxylic acid | Retained as Gut_lumen_2 based primarily on its spatial continuity with the broader Gut_lumen_2 region. |
| 2 | Gut_lumen_2, 100% | Interstitial_space | 7,253 (21.0%) | 68 | Taurine; histidine; sn-glycerol-3-phosphate; 3-phosphoglycerate; arginine | Reassigned as Interstitial_space because its spatial distribution did not form a clearly bounded gut-lumen region. Its muscle-associated metabolite profile was not considered sufficient for a more specific assignment, as well. This |

| | | | | | | |
|---|---|---|---|---|---|---|
| | | | | | | remained a tentative annotation that will be re-evaluated later round. |
| 3 | Gut_lumen_2, 97%; Hemolymph, 1%; Unannotated, 2% | Interstitial_space | 4,482 (13.0%) | 68 | Glutathione; histidine; glutamate; sn-glycerol-3-phosphate; glutamine | Reassigned as Interstitial_space because its spatial distribution did not form a clearly bounded gut-lumen region. This remained a working annotation. |
| 4 | Gut_lumen_2, 1%; Unannotated, 99% | Interstitial_space | 994 (2.9%) | 35 | Adenine; taurine; pyroglutamic acid; hypoxanthine; C18:2 | Reassigned as Interstitial_space because its spatial distribution did not form a clearly bounded gut-lumen region. This remained a tentative annotation that will be re-evaluated later round. |
| 5 | Interstitial_space, 100% | Interstitial_space | 3,038 (8.8%) | 54 | Adenine; pyroglutamic acid; guanine; succinate; hypoxanthine | Retained as Interstitial_space based on its previous annotation |
| 6 | Unannotated, 100% | Unannotated | 393 (1.1%) | 21 | O-phosphorylethanolamine; Allura Red_214; fructose-1,6-bisphosphate; ATP; D-erythrose-4-phosphate | Left unannotated because it contained very few pixels (~ 1 %). This remained a tentative annotation that will be re-evaluated later round. |

| | | | | | | |
|---|---|---|---|---|---|---|
| 7 | Gut_lumen_2, 1%; Hemolymph, 97%; Unannotated, 2% | MT_2 | 10,127 (29.3%) | 68 | Xanthurenic acid; uric acid; isoxanthopterin; indole-2-carboxylic acid; kynurenic acid | Reassigned as MT_2 based on its MT spatial distribution and enrichment of MT-associated metabolites. This indicated that the previous Hemolymph working annotation was more consistent with an MT-associated region. |
| 8 | Unannotated, 100% | Unannotated | 435 (1.3%) | 46 | Glutamine; O-phosphorylethanolamine; taurine; Allura Red_214; ATP | Left unannotated because its metabolite profile was mixed and its spatial distribution did not support a consistent tissue assignment. |
| 9 | Unannotated, 100% | Unannotated | 27 (0.1%) | 5 | Citrate/isocitrate; ATP; fructose-1,6-bisphosphate; Allura Red_214; D-erythrose-4-phosphate | Left unannotated because it lacked a sufficiently coherent spatial pattern and its mixed metabolite profile did not support a specific tissue assignment. |
| 10 | Unannotated, 100% | Unannotated | 688 (2.0%) | 18 | O-phosphorylethanolamine; histidine; Allura Red_214; ATP; fructose-1,6-bisphosphate | |
| 11 | Unannotated, 100% | Unannotated | 279 (0.8%) | 25 | Taurine; Allura Red_214; ATP; fructose-1,6-bisphosphate; D-erythrose-4-phosphate | |
| 12 | Gut_lumen_2, 1%; Interstitial_space, 28%; Unannotated, 72% | Unannotated | 383 (1.1%) | 25 | Xanthurenic acid; isoxanthopterin; glutamate; indole-2-carboxylic acid; guanosine | |

| | | | | | |
|---|---|---|---|---|---|
| 13 | Unannotated, 100% | Unannotated | 133 (0.4%) | 7 | GlucoseCl; glucose; deoxyribose-phosphate; trehalose; aconitate |
| 14 | Unannotated, 100% | Unannotated | 108 (0.3%) | 3 | 3-phosphoglycerate; sn-glycerol-3-phosphate; methionine sulfoxide; taurine; glutamine |
| 15 | Unannotated, 100% | Unannotated | 64 (0.2%) | 1 | Uric acid; malate; GlucoseCl; glucose-1-phosphate; citrate/isocitrate |
| 16 | Unannotated, 100% | Unannotated | 19 (0.1%) | 1 | N-acetylaspartic acid; N-acetyl-glutamate; hydroxyproline; N-acetyl-L-alanine; AMP/dGMP |
| 17 | Unannotated, 100% | Unannotated | 21 (0.1%) | 3 | Malate; ATP; fructose-1,6-bisphosphate; Allura Red_214; D-erythrose-4-phosphate |
| 18 | Unannotated, 100% | Unannotated | 81 (0.2%) | 4 | Uric acid; glucose-1-phosphate; ADP/dGDP; isoxanthopterin; trehalose-6-phosphate |
| 19 | Unannotated, 100% | Unannotated | 25 (0.1%) | 4 | Xanthurenic acid; ATP; fructose-1,6-bisphosphate; Allura Red_214; D-erythrose-4-phosphate |
| 20 | Unannotated, 100% | Unannotated | 22 (0.1%) | 2 | Asparagine; Allura Red_214; ATP; fructose-1,6-bisphosphate; D-erythrose-4-phosphate |
| 21 | Unannotated, 100% | Unannotated | 41 (0.1%) | 1 | O-phosphorylethanolamine; ATP; fructose-1,6-bisphosphate; Allura Red_214; D-erythrose-4-phosphate |
| 22 | Unannotated, 100% | Unannotated | 10 (<0.1%) | 2 | Uric acid; ATP; fructose-1,6-bisphosphate; Allura Red_214; D-erythrose-4-phosphate |
| 23 | Unannotated, 100% | Unannotated | 9 (<0.1%) | 1 | O-phosphorylethanolamine; ATP; fructose-1,6-bisphosphate; Allura |

| | | | | | |
|---|---|---|---|---|---|
| | | | | | Red_214; D-erythrose-4-phosphate |
| 24 | Unannotated, 100% | Unannotated | 14 (<0.1%) | 1 | O-phosphorylethanolamine; ATP; fructose-1,6-bisphosphate; Allura Red_214; D-erythrose-4-phosphate |
| 25 | Unannotated, 100% | Unannotated | 54 (0.2%) | 1 | O-phosphorylethanolamine; ATP; fructose-1,6-bisphosphate; Allura Red_214; D-erythrose-4-phosphate |
| 26 | Unannotated, 100% | Unannotated | 32 (0.1%) | 1 | O-phosphorylethanolamine; ATP; fructose-1,6-bisphosphate; Allura Red_214; D-erythrose-4-phosphate |
| 27 | Unannotated, 100% | Unannotated | 16 (<0.1%) | 1 | O-phosphorylethanolamine; ATP; fructose-1,6-bisphosphate; Allura Red_214; D-erythrose-4-phosphate |
| 28 | Unannotated, 100% | Unannotated | 16 (<0.1%) | 1 | O-phosphorylethanolamine; ATP; fructose-1,6-bisphosphate; Allura Red_214; D-erythrose-4-phosphate |
| 29 | Unannotated, 100% | Unannotated | 28 (0.1%) | 1 | O-phosphorylethanolamine; ATP; fructose-1,6-bisphosphate; Allura Red_214; D-erythrose-4-phosphate |
| 30 | Unannotated, 100% | Unannotated | 15 (<0.1%) | 1 | O-phosphorylethanolamine; ATP; fructose-1,6-bisphosphate; Allura Red_214; D-erythrose-4-phosphate |
| 31 | Unannotated, 100% | Unannotated | 23 (0.1%) | 1 | Pyroglutamic acid; ATP; fructose-1,6-bisphosphate; Allura Red_214; D-erythrose-4-phosphate |

| | | | | | | |
|---|---|---|---|---|---|---|
| 32 | Gut_lumen_2, 52%; Unannotated, 48% | Unannotated | 58 (0.2%) | 1 | Isoxanthopterin; trehalose; trehalose-6-phosphate; citrate/isocitrate; N-acetylaspartic acid | Left unannotated because it was detected in only one section and contained a mixed composition without a clear metabolite enrichment pattern |

**Table S17.** Round 13 – re-evaluation of subclusters from gut lumen, fat body, and neighboring tissue annotations.

| Subcluster | Round 12 annotation composition | Round 13 annotation | Pixels, n (%) | Sections containing pixels, n | Top enriched metabolites | Description |
|---|---|---|---|---|---|---|
| 0 | Gut_lumen_3, 100% | Unannotated | 1,557 (1.2%) | 52 | Uric acid; xanthurenic acid; kynurenic acid; indole-2-carboxylic acid; taurine | Left unannotated because its spatial and metabolite patterns were not consistent across all flies. Additionally, although it showed enrichment of several MT-associated metabolites, the spatial evidence was insufficient to support a reproducible MT or other tissue assignment. |
| 1 | Muscle, 99%; Gut_lumen_3, 1% | Muscle | 27,585 (21.0%) | 68 | Taurine; AMP/dGMP; 3-phosphoglycerate; ADP/dGDP; sn-glycerol-3-phosphate | Retained as Muscle based on its predominant previous composition, broad muscle-associated spatial distribution, and enrichment of taurine, adenylates, and glycolytic metabolites. |
| 2 | CNS, 58%; Fat_body, 40%; Interstitial_space, 1% | Fat_body_3 | 11,026 (8.4%) | 67 | O-phosphorylethanolamine; N-acetylaspartic acid; C18:2; C18:0; C16:0 | Assigned as Fat_body_3 based primarily on its spatial distribution and enrichment of O-phosphorylethanolamine and long-chain fatty acids. |
| 3 | CNS, 97%; Muscle, 2%; Interstitial_space, 1% | CNS | 6,170 (4.7%) | 59 | N-acetylaspartic acid; hydroxyproline; N-acetyl-L-alanine; N-acetyl-glutamate; C18:0 | Retained as CNS based on its predominant previous composition, CNS-associated spatial localization, and enrichment of N-acetylaspartic acid and associated neural metabolites. |
| 4 | Muscle, 89%; Interstitial_space, 9%; Gut_lumen_2, 1%; Gut_lumen_3, 1% | Muscle_2 | 27,616 (21.0%) | 68 | Histidine; taurine; methionine sulfoxide; glutamine; arginine | Assigned as Muscle_2 because it formed a distinct but muscle-associated spatial pattern in thorax. It was also derived from the previous Muscle annotation. It surrounds major muscle region in thorax. Taurine enrichment supported a muscle identity, whereas the absence of strong |

| | | | | | | |
|---|---|---|---|---|---|---|
| | | | | | | adenylate enrichment distinguished it from the main Muscle region and supported retaining it as a separate working annotation. |
| 5 | Interstitial_space, 100% | Unannotated | 977 (0.7%) | 65 | AMP/dGMP; ADP/dGDP; C18:2; arginine; 3-phosphoglycerate | Left unannotated because its spatial pattern did not provide sufficient evidence for a specific tissue assignment, even if muscle-associated metabolites were detected. The subcluster also represented less than 1% of the pixels included in this round. |
| 6 | Fat_body_3, 52%; Gut_lumen_2, 45%; Fat_body, 1%; Interstitial_space, 1% | Gut_lumen_2 | 9,442 (7.2%) | 67 | Xanthurenic acid; isoxanthopterin; uric acid; indole-2-carboxylic acid; guanosine | Assigned as Gut_lumen_2 as enriched metabolites do not correspond much to fat body and its spatial distribution aligns with gut_lumen_2 region. Because its enriched metabolites were also associated with MT regions, this remained a provisional working annotation. |
| 7 | Interstitial_space, 99%; Muscle, 1% | Hemolymph | 5,983 (4.6%) | 68 | Histidine; citrate/isocitrate; methionine sulfoxide; asparagine; glutathione | Provisionally assigned as Hemolymph based on its broadly distributed spatial pattern rather than a clearly bounded tissue morphology. Because the enriched metabolite profile did not provide strong independent support for a hemolymph assignment, this remained a working annotation for later re-evaluation. |
| 8 | Fat_body_2, 68%; Interstitial_space, 25%; Fat_body, 7% | Fat_body_2 | 16,962 (12.9%) | 68 | Glutathione; glutamate; malate; glucosamine-6-phosphate; UMP | Retained as Fat_body_2 based primarily on its correspondence with the broader Fat_body_2 region and its predominant previous composition. Its mixed composition and lack of a strongly fat-body-associated metabolite |

| | | | | | | |
|---|---|---|---|---|---|---|
| | | | | | | profile made this assignment provisional. |
| 9 | Fat_body, 95%; Fat_body_2, 4%; Fat_body_3, 1% | Fat_body | 18,916 (14.4%) | 67 | O-phosphorylethanolamine; glucose-1-phosphate; C16:0; C18:1; C16:1 | Retained as Fat_body based on its predominant previous composition, spatial distribution, and enrichment of O-phosphorylethanolamine and long-chain fatty acids. |
| 10 | Gut_lumen, 100% | Gut_lumen | 2,172 (1.7%) | 61 | Allura Red; O-phosphorylethanolamine; glucose-1-phosphate; Allura Red_214; isoxanthopterin | Retained as Gut_lumen based on its previous annotation, and enrichment of the ingested dye Allura Red. |
| 11 | Interstitial_space, 100% | Interstitial_space | 2,919 (2.2%) | 54 | Adenine; pyroglutamic acid; guanine; malate; succinate | Retained as Interstitial_space based on its previous annotation and diffuse spatial distribution between tissue regions. |
| 12 | CNS, 100% | Unannotated | 29 (<0.1%) | 3 | N-acetylaspartic acid; uridine; ribose-phosphate; Allura Red_214; D-erythrose-4-phosphate | Left unannotated because it contained very few pixels, occurred in only three sections despite its previous CNS annotation and enrichment of N-acetylaspartic acid. |

**Table S18.** Round 14 - final consolidation and nomenclature standardization of tissue annotations.

| Final annotation | Round 13 annotation composition | Pixels, n (%)* | Final update | Description |
|---|---|---|---|---|
| CNS | CNS, 100% | 6,170 (3.3%) | Retained | Retained based on reproducible CNS-associated spatial distribution and enrichment of N-acetylaspartic acid. |
| Eye | Eye, 100% | 2,883 (1.5%) | Retained | Retained based on its reproducible localization within an eye-associated region across sections. |
| Eye_2 | Eye_2, 100% | 6,754 (3.6%) | Retained | Retained as a second eye-associated subregion because it showed a spatial pattern distinct from the primary Eye region and was located around its periphery. |
| Fat_body | Fat_body, 100% | 18,916 (10.1%) | Retained | Retained based on its fat body spatial distribution and enrichment of O-phosphoethanolamine and long-chain fatty acids. |
| Fat_body_2 | Fat_body_3, 100% | 11,026 (5.9%) | Renamed | Retained as a distinct fat body subregion based on its predominantly non abdomen-associated spatial pattern and lipid-associated metabolite profile. It was renumbered from Fat_body_3 to Fat_body_2 to standardize the final nomenclature. |
| Fat_body_3 | Thoracic_fat_body, 100% | 6,038 (3.2%) | Renamed | Renamed as Fat_body_3 to standardize the nomenclature of the three fat body subregions. Its thoracic distribution and enrichment of O-phosphoethanolamine and long-chain fatty acids supported a fat body identity. |
| Gut_lumen | Gut_lumen, 100% | 2,172 (1.2%) | Retained | Retained based on its spatial location and enrichment of the ingested dye Allura Red. |
| Hemolymph_crop | Hemolymph, 57%; Crop, 43% | 10,514 (5.6%) | Merged and renamed | The Hemolymph and Crop working annotations were combined because they showed similar enrichment of trehalose and glucose. The putative crop region was also detected in only a small number of sections, providing insufficient support for retaining it as a separate annotation. The combined Hemolymph_crop label preserves this remaining uncertainty rather than assigning these pixels definitively to either hemolymph or crop. |
| Hindgut | MT_2, 100% | 10,127 (5.4%) | Reassigned | Reassigned as Hindgut based on its posterior, gut-associated spatial pattern during the final review. |

| | | | | |
|---|---|---|---|---|
| MT | MT, 100% | 10,072 (5.4%) | Retained | Retained based on its consistent spatial pattern and strong enrichment of uric acid, isoxanthopterin, xanthurenic acid, and kynurenic acid. |
| MT_2 | Gut_lumen_2, 100% | 9,442 (5.0%) | Reassigned | Reassigned as MT_2 because it was spatially adjacent to the primary MT region and showed enrichment of uric acid, isoxanthopterin, kynurenine, and related metabolites, which was more consistent with an MT region than with the gut lumen. |
| MT_3 | Hindgut, 100% | 6,315 (3.4%) | Reassigned | Reassigned as MT_3 because it was spatially associated with other MT regions and showed an enriched metabolite profile more consistent with an MT region than with hindgut. The numbered label distinguishes this spatial subregion without assigning it to a specific Malpighian tubule segment. |
| Muscle | Muscle, 100% | 27,585 (14.7%) | Retained | Retained based on its broad muscle-associated spatial distribution in the thorax and enrichment of taurine, adenylates, and glycolytic metabolites, which together supported a muscle identity. |
| Muscle_2 | Muscle_2, 100% | 27,616 (14.7%) | Retained | Retained as a second muscle region because it showed a distinct spatial pattern adjacent to the Muscle region and was enriched in muscle-associated metabolites, including taurine. Differences in its overall metabolite profile, including lower adenylate enrichment, distinguished it from Muscle region and supported its retention as a separate subregion. |
| Reproductive_system | Fat_body_2, 100% | 16,962 (9.0%) | Reassigned | Reassigned as Reproductive_system because its spatial pattern was associated with reproductive-system structures rather than the broad peripheral distribution expected for fat body. Its overall metabolite profile, including enrichment of UMP, glutamate, glutathione, and related metabolites, was also more consistent with the other reproductive-system-associated region than with the principal Fat_body regions. |
| Reproductive_system_2 | Reproductive_system, 100% | 4,355 (2.3%) | Renamed | Retained as a reproductive-system-associated region based on its spatial location and metabolite profile but renamed Reproductive_system_2 to distinguish it from the larger region reassigned as Reproductive_system during the final review. The numbered labels distinguish |

| | | | | |
|---|---|---|---|---|
| | | | | spatial and metabolic subregions rather than established anatomical subdivisions. |
| Unannotated | Unannotated, 52%; Interstitial_space, 27%; Oenocyte, 21% | 10,762 (5.7%) | Consolidated | Existing Unannotated pixels were combined with the Interstitial_space and provisional Oenocyte annotations because these regions lacked sufficient independent spatial and metabolite evidence for distinct tissue assignments. They were conservatively retained as Unannotated rather than overinterpreted. |

*Pixel percentages indicate the proportion of all 187,709 pixels in the integrated atlas. Round 14 involved final annotation review, consolidation, and nomenclature standardization without additional subclustering.
**Numbered suffixes denote subclusters within a broader tissue category (e.g., Muscle, MT, or Reproductive_system).
